# Single-cell somatic copy number variants in brain using different amplification methods and reference genomes

**DOI:** 10.1101/2023.08.07.552289

**Authors:** Ester Kalef-Ezra, Zeliha Gozde Turan, Diego Perez-Rodriguez, Ida Bomann, Sairam Behera, Caoimhe Morley, Sonja W. Scholz, Zane Jaunmuktane, Jonas Demeulemeester, Fritz J Sedlazeck, Christos Proukakis

**Affiliations:** Department of Clinical and Movement Neurosciences, UCL Queen Square Institute of Neurology, London, UK; Aligning Science Across Parkinson’s (ASAP) Collaborative Research Network, Chevy Chase, MD, 20815; Human Genome Sequencing Center, Baylor College of Medicine, One Baylor Plaza, Houston TX 77030, USA; Neurodegenerative Diseases Research Unit, National Institute of Neurological Disorders and Stroke, Bethesda, MD, USA; Department of Neurology, Johns Hopkins University Medical Center, Baltimore, MD, USA; Queen Square Brain Bank for Neurological disorders, UCL Queen Square Institute of Neurology, London, UK; Department of Oncology, KU Leuven, Leuven, Belgium; Cancer Genomics Laboratory, The Francis Crick Institute, London, UK; VIB Center for Cancer Biology, Leuven, Belgium; Department of Molecular and Human Genetics, Baylor College of Medicine, TX, USA; Department of Computer Science, Rice University, 6100 Main Street, Houston, TX, USA

## Abstract

The presence of somatic mutations, including copy number variants (CNVs), in the brain is well recognized. Comprehensive study requires single-cell whole genome amplification, with several methods available, prior to sequencing. We compared PicoPLEX with two recent adaptations of multiple displacement amplification (MDA): primary template-directed amplification (PTA) and droplet MDA, across 93 human brain cortical nuclei. We demonstrated different properties for each, with PTA providing the broadest amplification, PicoPLEX the most even, and distinct chimeric profiles. Furthermore, we performed CNV calling on two brains with multiple system atrophy and one control brain using different reference genomes. We found that 38% of brain cells have at least one Mb-scale CNV, with some supported by bulk sequencing or single-cells from other brain regions. Our study highlights the importance of selecting whole genome amplification method and reference genome for CNV calling, while supporting the existence of somatic CNVs in healthy and diseased human brain.

## Introduction

Mosaicism, due to somatic mutations in the human brain, is increasingly recognised, with likely roles in neurodevelopmental and neurodegenerative diseases ^1^ ^2–4^. As the ‘signal’ of a low-level somatic mutation can be lost in ‘bulk’ tissue homogenates, single-cell whole genome sequencing (scWGS) after whole genome amplification (WGA) has been increasingly utilized in the past decade ^5^. Megabase-scale copy number variants (CNVs) can be detected using read-depth methods even with one or a few million short reads ^6–8^. Several studies of human single cortical neurons showed Megabase-scale CNVs, although the precise frequency of these changes remains unclear ^9^ ^7^ ^10^ ^11^ ^12^. These CNVs may be more frequent in younger than aged healthy brains ^7,8^, arising in embryonic neurogenesis in mouse^8^, with complex structural variants (SVs) also arising in human neurogenesis^13^. We previously performed targeted detection of somatic CNVs in two related neurodegenerative disorders, Parkinson’s disease (PD) and multiple system atrophy (MSA). These diseases are summarized under the umbrella term of synucleinopathies, related to abnormal aggregation of misfolded alpha-synuclein protein, encoded by the gene *SNCA*. Using fluorescent *in situ* hybridisation, we found somatic *SNCA* CNVs in PD and MSA brain tissue^14^ associated with pathology in MSA at the single-cell and regional level^15,16^.

A range of WGA options exist, such as PCR-based, Multiple Displacement Amplification (MDA, isothermal, using phi-29, a DNA polymerase with high fidelity and extreme processivity ideal for amplifying large fragments), and hybrid methods^5^. Still, one needs to better understand the advantages and disadvantages/biases across these WGA methods, as it impacts the detection and interpretation of CNV calls across cells. This is especially needed outside control tissues or cell lines, as often these do not encapsulate the challenges faced across e.g., brain tissue. In our previous work^16^, we used PicoPLEX, a hybrid method related to multiple-annealing and looping-based amplification cycles (MALBAC), to capture genome-wide somatic CNVs in diseased human brain tissue for the first time. We detected CNVs in ∼30% of cells in two MSA cases in affected brain regions^16^, although their relevance remains unclear. We note several previous comparisons of WGA methods, with the broad consensus being that MDA is not suitable for calling CNVs by read depth, as there is bias due to over-amplification of certain regions, although it is more accurate at the single-base level ^17–23^. There have been several attempts to reduce MDA amplification bias by performing reactions in nanoliter-scale volumes^10,24,25^. MDA performed after the single-cell genome is partitioned into ∼50,000 droplets (droplet MDA, or dMDA) in the Samplix X-Drop device was reported to yield more even amplification^26^. The recently developed method of Primary Template-directed Amplification (PTA) also utilizes phi29, but the reaction is terminated early to avoid over-amplification^27^. High coverage (>30x) scWGS after PTA allows detection of single nucleotide variants (SNVs) and indels, already been applied to normal and Alzheimer’s disease brain^28,29^, and also shows lower coverage variability than MDA ^28^.

In this work we provide deeper insights into the advantages and disadvantages of three WGA approaches currently available commercially (PicoPLEX, PTA, dMDA), the first two of which are easily scalable. We perform their first direct comparison using human *post-mortem* frozen brain samples, including disease brain (two MSA cases) and one control donor. We found that in our hands both PicoPLEX and PTA are suitable for CNV calling by shallow WGS, but not dMDA. PTA amplifies more the human genome, but PicoPLEX is the least noisy method. Furthermore, we compared chimeras generated by these methods, which could hinder precise SV detection or introduce other biases in the analysis. We also investigated the utility of alignment to a complete reference genome (T2T-CHM13) as compared to the commonly used hg38 reference genome, and liftover back to hg38.

## Results and Discussion

To optimize the workflow for detection of large CNVs by scWGS, we compared three different WGA methods, PicoPLEX, PTA, dMDA. We used single-nuclei isolated from the cortex of up to three different brains, two with MSA and one control (**Fig. 1a**). We had previously reported scWGS from other regions of the same MSA brains^16^. We also used non-brain control nuclei in selected experiments: fibroblasts carrying a germline *SNCA* triplication, and lymphocyte nuclei from NA12878. In total, we amplified and performed Illumina sequencing (paired-end) across 93 brain cells, as well as 3 fibroblasts and 5 NA12878 Β-lymphocytes (mean coverage ∼0.64x, 0.17x and 0.71x respectively) (**Fig. 1b**, **Supplementary Fig. 1a**).

**Fig. 1.**
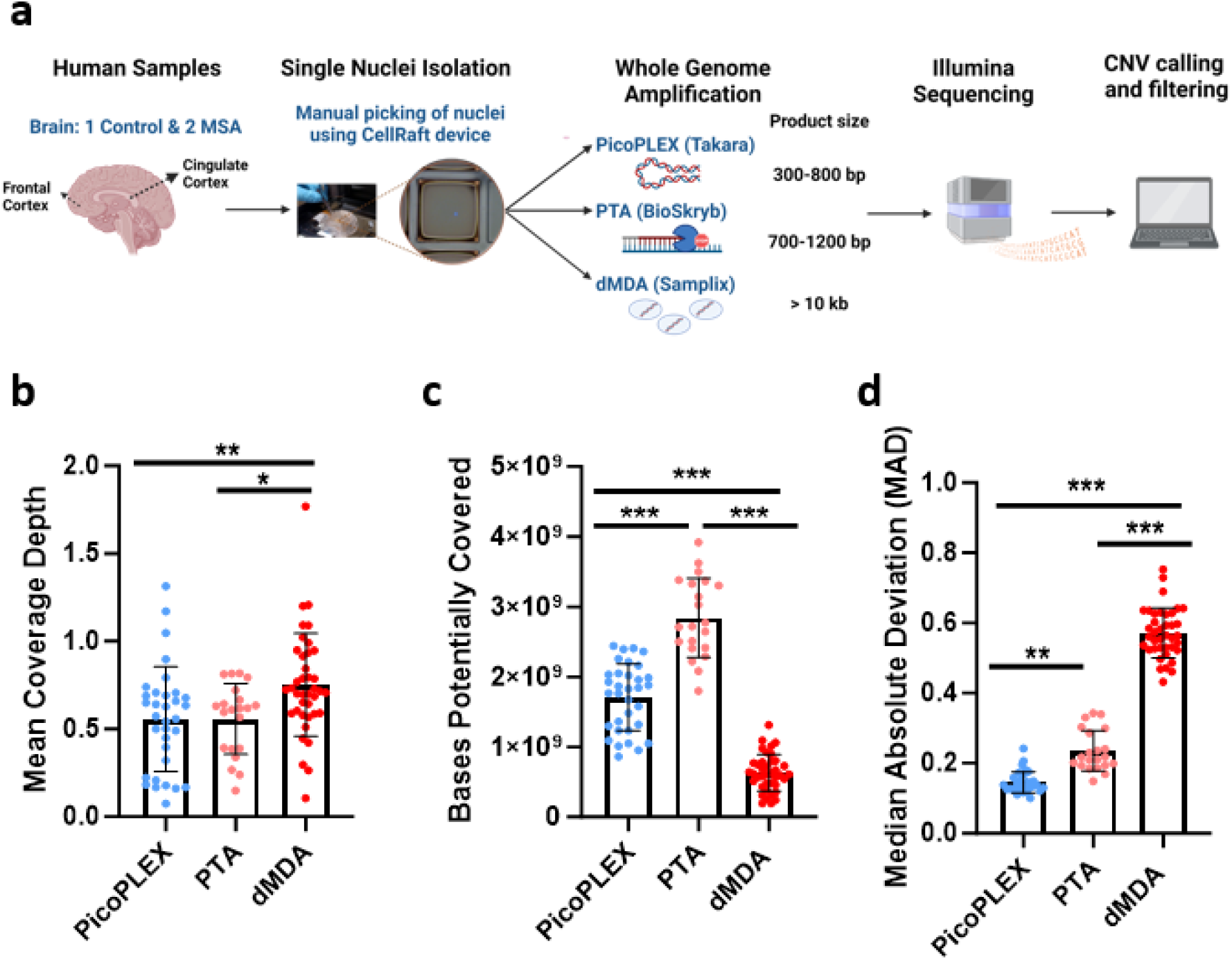
Experimental overview and scWGS preliminary analysis from human *post-mortem* brain samples. **a** Methodology overview created with BioRender.com (agreement KB25LPBTEK). **b** Mean depth of coverage for each WGA method. PicoPLEX vs PTA ns (adj. *p*>0.99), PicoPLEX vs dMDA ** (adj. *p*=0.008) and PTA vs dMDA * (adj. *p*=0.04). Kruskal-Wallis test with Dunn’s multiple comparisons correction. **c** Bases potentially covered if sequenced deeper. P value for all pairwise comparisons <0.001. Brown-Forsythe and Welch ANOVA tests with Dunnett’s T3 multiple comparisons correction. **d** Median Absolute Deviation (MAD) scores. P value for all pairwise comparisons <0.001 (adj. *p*<0.001). Kruskal-Wallis test with Dunn’s multiple comparisons correction. For b-d PicoPLEX (*n*=33), PTA (*n*=21), dMDA (*n*=39); Mean ± SD shown.

### PTA provides the broadest amplification, but PicoPLEX provides the most even

Although these are “whole” genome amplification methods, some regions may not be amplified in a given cell (locus dropouts)^5^. In the absence of high coverage WGS of each cell, the maximum number of bases which could be retrieved by deeper sequencing can be deduced genome-wide using Preseq^30^. We found that PTA provides efficient capture of the genome of brain nuclei, consistent with other data^27,28^ (**Fig. 1c**; mean ± SD: 2.84 ± 0.56 Gb), with PicoPLEX the next best performer (1.71 ± 0.48 Gb). dMDA provided the lowest breadth of coverage (0.75 ± 0.29 Gb), even lower than the report of only one-third of the genome covered with 100 million reads after dMDA^26^.

CNV calling by read depth is hampered by uneven amplification, as regions which are over- or under-amplified could appear as CNVs. We compared a key metric, the median absolute deviation (MAD) between adjacent bins, between the three technologies at 500 kb bin size using *Ginkgo*^6^, after adapting this widely used validated tool to hg38 (see methods). We noted clear differences across methods, with PicoPLEX performing best and dMDA worst (**Fig. 1d**, mean ± SD: PicoPLEX 0.15 ± 0.03, PTA 0.24 ± 0.06, dMDA 0.57 ± 0.07). This is reflected by visual review of sequencing traces (**Fig. 2a**), and Lorenz curves, which indicate the amplification variation by plotting the cumulative fraction of reads as a function of the cumulative fraction of the genome (**Fig. 2b**). In MSA1, where we had used different PicoPLEX versions, and different library preparation for PTA, we verified that the MAD was not affected by this (**Supplementary Fig. 1b-c**). Furthermore, the MAD for each WGA method was similar between different brains, and non-brain samples (**Supplementary Fig. 1d-f**). The MAD values we obtained for PTA are similar to the ones reported (∼0.25 for bin size range 100-1,000 kb)^27^. The uneven coverage of dMDA in particular cannot be explained by GC content variation, although there was a slight drop in coverage of high GC regions for both dMDA and PTA (**Fig. 2c**). Indeed, others have reported no major GC effect on MDA bias^23^. Although previous reports had shown superiority of PicoPLEX or MALBAC over MDA for CNV calling^18–23,31,32^, there had been no prior comparison to dMDA. The previous study of dMDA, conducted in lymphoblasts, reported more even genome coverage compared to MDA performed in parallel but not in droplets, although MAD values were not reported^26^. As the MAD values we obtained using dMDA were poor, we recalculated MAD after gradually increasing the bin size up to 5 Mb, which improved the values as expected^9^. We also found that MAD can be further improved by removing noise using principal component analysis (see methods)^33^ (**Supplementary Fig. 2**). CNV calling in dMDA might therefore be possible for very large aberrations after denoising.

**Fig. 2.**
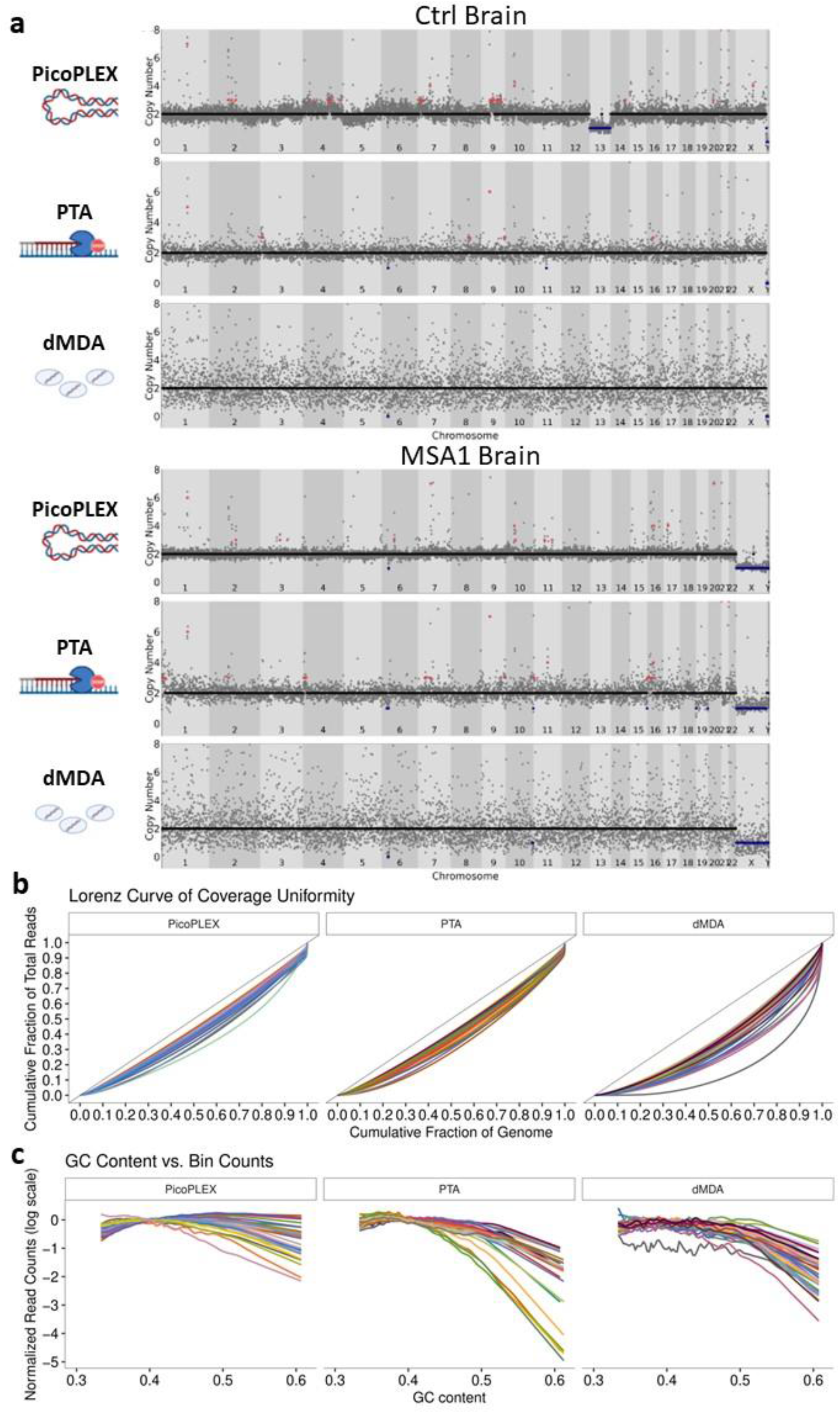
scWGA method comparison from brain analyzed by *Ginkgo* at 500 kb bin size. **a** Visual view of copy number profiles of single-nuclei amplified by PicoPLEX, PTA and dMDA from the control (top) and MSA1 (bottom) brains. **b** Lorenz curves. The black lines with slope 1 represent perfect coverage uniformity. Increasing divergence of the curve of each cell from this indicates lower overage uniformity. **c** Effect of GC content in scWGA. For b-c PicoPLEX *n*=33, PTA *n*=21, dMDA *n*=40.

### Each WGA method has propensity for different types of chimeras

All WGA methods can lead to chimeras. These could be misinterpreted as SVs, but also, if they cluster in certain parts of the genome, could impact CNV calls. We therefore aimed to compare the frequencies of key types between the three WGA methods (**Fig. 3**). For this comparison, PTA cells which had a different library preparation method were analyzed separately, as they had a divergent profile presumably related to this (**Supplementary Fig. 3**). In paired-end sequencing, the read pairs should both be pointing inwards, towards each other. Outward read pairs, indicative of tandem duplications, were most frequent in PicoPLEX, and least frequent in PTA. This observation is consistent with a previous report that over half of PicoPLEX artifacts appear as duplications^17^. Apparent translocations were most frequent in PTA and least frequent in dMDA. Other orientations, which include inversions, were most frequent in PTA, and hardly ever seen in PicoPLEX, but relatively common in dMDA, as previously reported in MDA, where they had comprised almost all the chimera signatures^17^.

**Figure 3.**
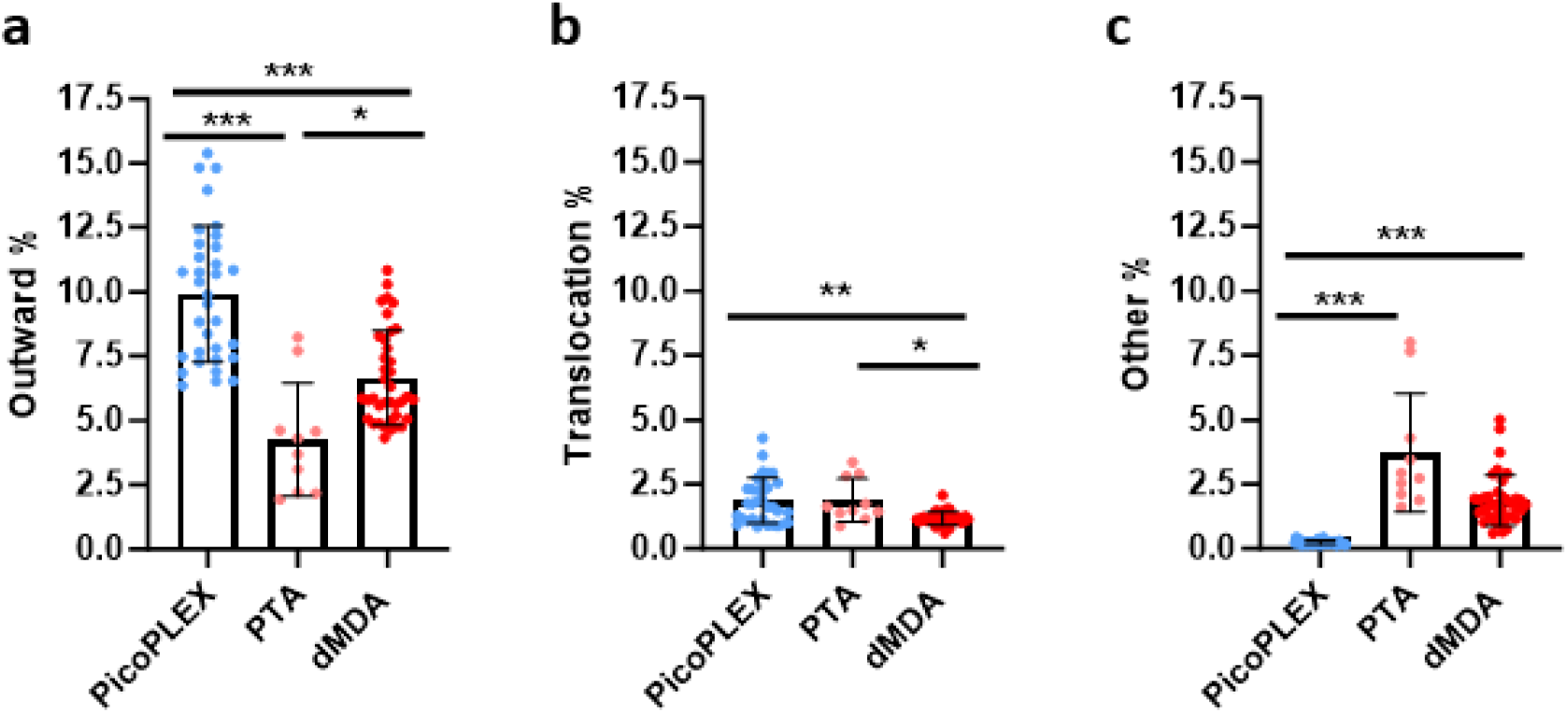
Comparison of discordant read pairs of brain nuclei amplified with each method. **a** Outward pairs. Differences analyzed using Brown-Forsythe and Welch ANOVA test with Dunnett’s T2 multiple comparisons test. PicoPLEX vs PTA *** (adj. *p*<0.001), PicoPLEX vs dMDA *** (adj. *p*<0.001), PTA vs dMDA * (adj. *p*=0.02). **b** Pairs on different chromosomes indicating translocations. Differences analyzed by Kruskal-Wallis test with Dunn’s multiple comparisons test. PicoPLEX vs PTA ns (adj. p>0.99), PicoPLEX vs dMDA ** (adj. *p*=0.002), PTA vs dMDA *** (adj. *p*=0.02). **c** Pairs in other orientations. Differences analyzed by Kruskal-Wallis test with Dunn’s multiple comparisons test. PicoPLEX vs PTA *** (adj. p<0.001), PicoPLEX vs dMDA *** (adj. *p*<0.001), PTA vs dMDA ns (adj. *p*=0.18. a-c PicoPLEX (*n*=33), PTA (*n*=10). dMDA (*n*=38).

### Realignment to T2T-CHM13 and liftover to hg38 affects CNV calling

Next we investigated the impact of the reference genome for the analysis of single-cell CNV. We realigned all data to the T2T-CHM13 genome^34^ and performed liftover of the read alignments back to hg38 using *levioSAM2*, which improves calling on the original reference, both for small variants, and for structural variants using long reads.^35^ We compared relevant metrics between the different alignments. We found a marginal improvement in the percentage of reads aligned, which was highest in T2T-CHM13 and partly maintained in hg38 after liftover (**Supplementary Fig. 4a**). For both amplification methods, the MAD slightly improved in the T2T-CHM13 and liftover alignments compared to the original hg38 (**Supplementary Fig. 4b**; see also methods). The confidence score was not affected by genome changes (**Supplementary Fig. 4c**; see also methods). The breadth of sequencing (bases potentially covered using *Preseq*) was improved using T2T but was lower in the liftover genome (**Supplementary Fig. 4d**).

To determine which brain cells were suitable for CNV calling by *Ginkgo*, we set strict thresholds. We discarded cells with MAD score >0.3 as before^16^, although even higher values have been considered acceptable^8,9^. We also calculated the confidence score, which indicates the extent to which genomic segments have integer copy numbers, rather than intermediate copy number states which may indicate uneven amplification,^11^ and retained only cells with confidence score ≥0.7. We used 250 kb bins for PicoPLEX, for consistency with our previous work, which left 23 of 34 cells (**Table 1)**. We initially compared unfiltered PicoPLEX and PTA CNV calls between the different genome alignments (**Supplementary Table 1**). T2T-CHM13 alignment followed by liftover to hg38 led to 65% fewer losses called than hg38. Liftover also increased gains by >10%, but the T2T-CHM13 gains were even higher (an additional 14%, or 2.15 CNVs per cell). We reviewed hg38-specific losses and noted that they were mostly shared between cells and individuals, did not appear robust (as they appeared to be non-integer, intermediate between copy numbers 1 and 2), and were often around centromeres. The *SNCA* germline triplication (1.85 Mb), which we had also used as a positive control before, was detected in all three PicoPLEX-amplified fibroblasts, including one which failed confidence score filter, regardless of reference used (**Supplementary Fig. 5**).

**Table 1.** Success rates and overview of filtered CNV calls across all brain samples (hg38-liftover).

|  | Cells passing QC (%) |  | Cells with at least 1 CNV (%) |  | CNVs called |  |  |
| --- | --- | --- | --- | --- | --- | --- | --- |
|  | PicoPLEX | PTA | PicoPLEX | PTA | Total | Gains | Losses |
| MSA1 | 11/15 (73.3%) | 5/13 (38.5%) | 3<br>(27.3%) | 4<br>(80%) | 26 | 16 | 10 |
| MSA2 | 8/12 (66.7%) | x | 3<br>(37.5%) | x | 4 | 3 | 1 |
| Control | 4/7<br>(57.1%) | 6/12 (50.0%) | 1<br>(25.0%) | 2<br>(33.3%) | 6 | 4 | 2 |

For PTA-amplified cells, we used 500 kb bins, as even at this size only 11 of 25 cells passed our criteria. We thus decided to focus CNV-calling on the hg38-liftover genomes, limited to autosomes, as this is better annotated, and the results are easier to compare than T2T-CHM13.

### Megabase-scale CNVs are detected in over a third of brain cells

We proceeded to somatic CNV calling in PicoPLEX and PTA-amplified brain cells; the profiles of all cells and CNVs called are given in **Supplementary Fig. 6**. To perform stringent filtering, we first removed all calls that were shared between at least half of the cells, or any cells between different individuals. To assign a numerical threshold, we calculated the median copy number of each segment assigned by *Ginkgo* and plotted the distribution of segments with each copy number, from which we set copy number thresholds of 1.29 for a loss and 2.80 for a gain (methods and **Supplementary Fig. 7**). We confirmed that in males, excluding one cell with possible chrX aneusomy, CN was below that number in both PicoPLEX and PTA.

This led to a total of 36 CNVs across the three brains, with gains predominant (63.9%; **Table 1**; all CNVs listed in **Supplementary Table 2**, profiles shown in **Supplementary Fig. 6**), CNVs were found in 38.2% of cells (13/34 overall; 8/23 PicoPLEX, 6/11 PTA; *p*=0.46). The proportion of cells with CNVs were similar in MSA (10/24) and controls (3/10; *p*=0.7). This is very similar to the ∼30% of cells we estimated to carry somatic CNVs in other regions of the same MSA brains^16^. For controls it appears higher than previous estimates in cortical neurons, with 10-25% reported to carry Mb-scale CNVs^7^, and more recently only 2 of 52 PTA-amplified cortical neurons carrying CNVs >∼5 Mb ^28^. In the cells which had CNVs, the mean number per cell was 2.8, although notably one MSA cell had 17 (11 gains and 6 losses; cell A24; **Supplementary Fig. 6**). To ensure that CNVs are not associated with low mapping quality, we conducted the same analyses after filtering for reads with mapping quality below 10. All CNVs were detected, except one in A24.

Examining CNVs further, these were slightly larger in the PTA data, at least partly due to the higher bin size used (median 5.15 Mb v 3.36 Mb, Mann-Whitney *p*=0.22; **Supplementary Table 2**). Gains and losses had similar sizes in PicoPLEX data (3.31 and 3.21 Mb respectively), although in PTA gains were larger (11.44 v 5.65 Mb). One control brain cell had two distinct losses which essentially added up to a loss of chromosome 13, with only 2.3 Mb spared, which presumably is an error, and this chromosome is lost in its entirety. This was the only aneusomy seen in a brain cell, consistent with estimates of brain aneuploidy of 0.7-5% derived by scWGS^1^, although one of three fibroblasts did have a chromosome gain (**Supplementary Fig. 5**). We noted that 9 brain CNVs (27.8%) were sub-telomeric. This observation is in agreement with previous work suggesting enrichment in these regions which are rich in segmental duplications, but considerably higher than the 9.15% in MSA neurons from other brain regions which we previously reported^16^. To understand the nature of genes affected by CNVs, we performed gene ontology analysis using PANTHER^36^. We noted divergent enrichment in MSA and control, but due to the sample size we cannot draw any conclusions (**Supplementary Fig. 8**).

We also investigated whether CNV calls are supported by another algorithm for scWGS CNV calling, *Copykit*,^37^ also based on circular binary segmentation, which uses hg38 as default (**Supplementary Table 2**). The *SNCA* triplication was detected in all three fibroblasts, although the copy number was given as 3 (rather than 4) in the one that failed confidence score. We allowed for a smaller CNV called by *Copykit* to be classed as supportive of a CNV called by *Ginkgo* if it was encapsulated by the *Ginkgo* CNV region. The majority of *Ginkgo* CNV calls in brain were supported (75.8% for PicoPLEX and 71.4% for PTA). The CNVs in PicoPLEX cells not reported by *Copykit* were all <3.5 Mb, except for a centromeric one. The median CNV size was identical (3.52 Mb PicoPLEX, 3.51 Mb *Copykit*). For CNVs called by both, *Copykit* sizes were slightly smaller (median difference −0.54 Mb), with three notable large CNVs extending to the short arm telomere being much smaller (by 19-35 Mb) in *Copykit*. As *Copykit* has not been formally benchmarked to our knowledge, while *Ginkgo* has been found to be accurate in calling breakpoints^38^ we gave it the benefit of the doubt for estimating CNV sizes. *Ginkgo* appears to have an advantage in detecting relatively small CNVs, and centromeric and telomeric regions.

### Some CNVs have support in bulk or other brain regions

If a somatic CNV is limited to a single-cell, orthogonal validation is by definition impossible. A clonal CNV, however, could be detectable in other cells, from the same or other brain regions. It may also be detectable by bulk sequencing of adequate depth. This could, however, be compromised by a number of issues, including imprecise boundaries in single-cell CNV calling, possible non-amplification of the breakpoints due to allelic drop-out, and the intrinsic limitations in SV calling if short-read sequencing is used. To investigate this in the two MSA brains, we first reviewed the presence of CNVs with similar boundaries in previously reported different regions from the same brains: substantia nigra in both (*n*=99), and the pons and putamen in MSA1 (*n*=70)^16^. One 3.36 Mb gain on chromosome 9 in MSA2 (cell A76; **Supplementary Fig. 6)** was essentially identical to a gain which had already been suspected of being clonal, as one breakpoint had been shared and one located nearby in two cells from the substantia nigra of the same brain, one neuron and one non-neuron. This includes the *TLR4* gene, which encodes a microglial and astroglial receptor with a role in alpha-synuclein clearance and pathology propagation^39^, and is upregulated in MSA^40^. Although we did not detect a gain of this gene in the other MSA brain, its presence in a clonal CNV makes further study worthwhile.

To seek support for somatic CNV calls in the MSA brains, we also analyzed deep bulk short-read WGS from the MSA brains, specifically the cingulate cortex (where the single-cells were obtained from) and the adjacent cingulate white matter for both cases, as well as the cerebellar cortex and white matter from MSA2 (mean coverage across all 83.6x; **Supplementary Table 3**). We used Samplot^41^ to visualize any read pairs consistent with each reported CNV in all bam files, scanning 250 kb on each side of *Ginkgo*-reported breakpoints. We considered any read pair in the WGS from the same sample and region consistent with the called CNV as tentative support, as long as it was in a region with good coverage, and, assuming that somatic CNVs would not be shared between individuals, nothing similar was found in the other brain. We found tentative support (at least one read pair each) for 4 CNVs, 2 gains and 2 losses, all <10 Mb in size, with support also in DNA from the adjacent white matter of the same brain in one (**Fig. 4**; the images of these genomic regions from all WGS samples are shown in **Supplementary Fig. 9**).

**Fig. 4:**
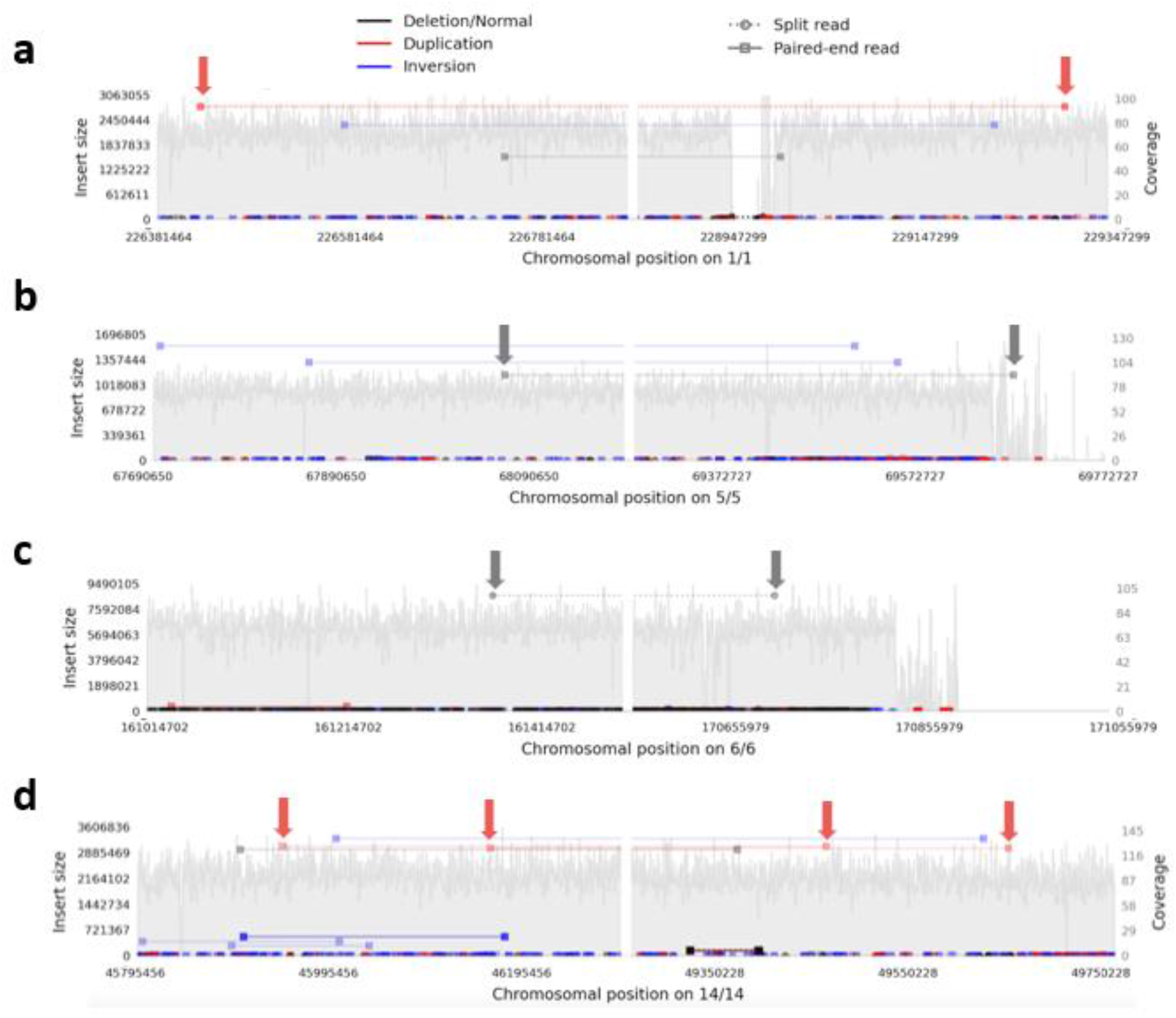
Samplot views demonstrating tentative support for single cell CNV calls in bulk short read WGS. These show all read pairs in the designated regions in cingulate cortex, with pairs supporting a particular type of CNV / SV indicated according to the scheme at the top, and those supporting each single-cell CNV arrowed. The left and right panel of each plot show the regions around the reported proximal and distal breakpoint respectively, which is at the middle of each panel, with the chromosomal numbers and positions on the x axis below. The y axis indicates the calculated insert size for the read pairs of interest on the left, and the local coverage on the right. **a-c** MSA1 and **d** MSA2. **a** duplication 2.47 Mb (cell A24), **b** deletion 1.58 Mb (cell A24), **c** deletion 9.54 Mb (cell L21), **d** duplication 3.45 Mb (cell A82). A read pair identical to **c** was found in the cingulate white matter, suggesting mosaicism across both brain regions.

### Putative gains in T2T-CHM13 specific regions may be due to low mapping quality but merit further assessment

We finally addressed the T2T-CHM13 specific gains (gains called only in the T2T-CHM13 alignments, with at least 50% of their span being novel T2T-CHM13 sequence). Again, we filtered those shared between >50% of cells or different individuals, to focus on most likely somatic events. We identified three gains, two involving centromeric active α-sat higher order repeat (HOR) arrays^42^. A chromosome 13 acrocentric arm gain had surprisingly high copy numbers (15-16). We reviewed the copy number of the region covered in all other cells, and we noted that the PicoPLEX data were much more noisy than PTA in two of these regions (**Fig. 5**). We investigated the likelihood of these calls by restraining the mapping quality and assessing how this impacts the CN estimation. As previously reported, the alignment to the T2T-CHM13 reference results in less noisy and more reliable alignments for short reads^35^. Nevertheless, these T2T-CHM13 unique regions have specific challenges due to their repetitiveness^34^. We filtered the reads based on mapping quality (MQ) 0, 1 and 10 to assess different stringencies. For MQ1, we observed that the single copy number gain was removed, but the two high CN gains persisted at MQ1, and were removed at MQ10. At MQ10, however, many cells had apparent losses in these regions, due to the poor mappability. This illustrates the difficulty at calling even large CNVs in these regions. We approached this by filtering more stringently across samples and CN directly, since false positives should manifest in all samples if it is due to mapping or reference biases. Since this is not the case in these CN candidates, the possibility of true somatic gains cannot be excluded.

**Fig. 5.**
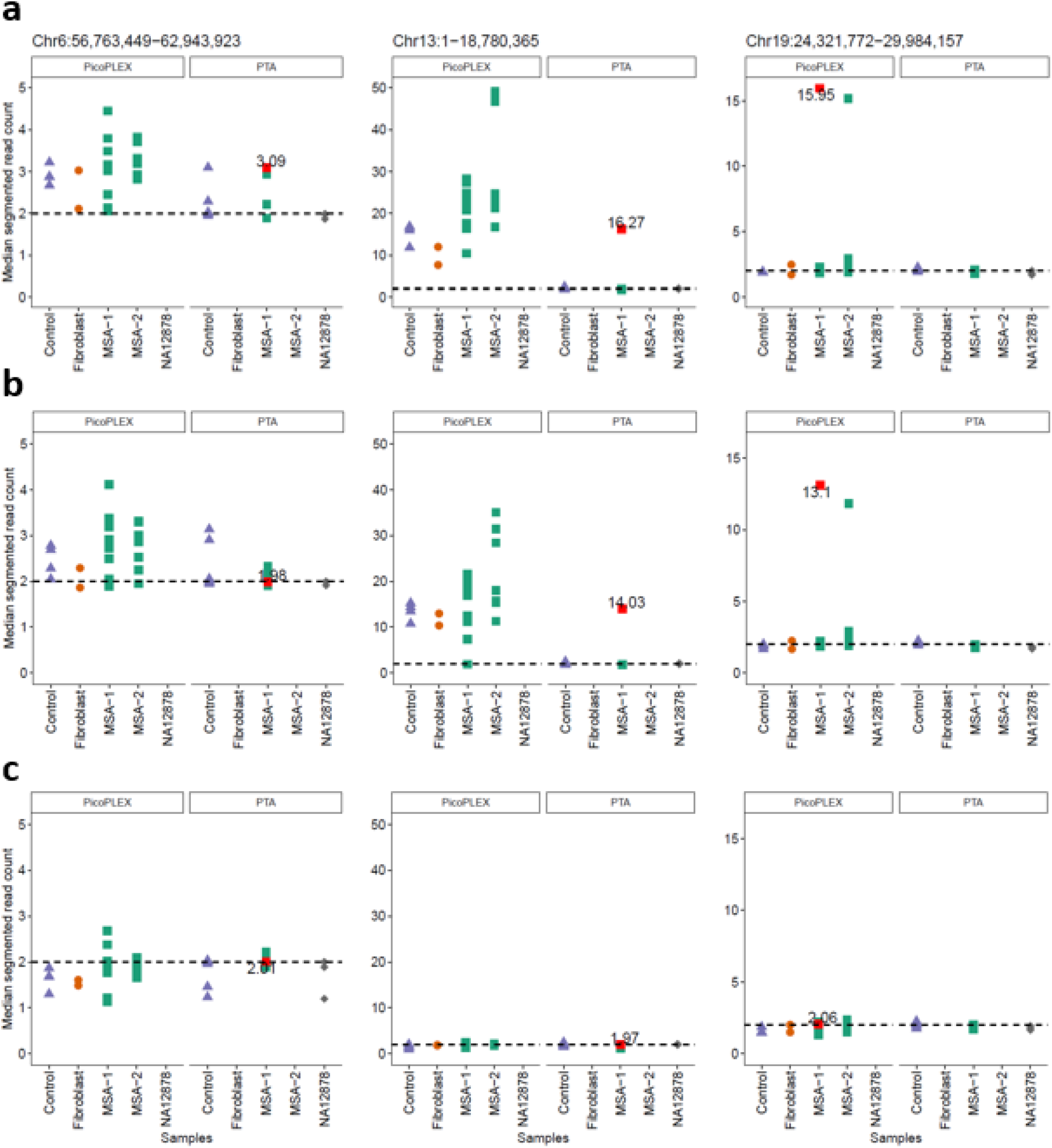
Evaluation of T2T-CHM13 specific CNV calls using different WGA methods (PTA and PicoPLEX) and mapping quality filtering. **a** no mapping quality filter, **b** mapping quality 1, **c** mapping quality 10.

### Conclusions

Whole genome amplification allows the analysis of single-cell genomes for somatic mutations. Nowadays there are several available methods to perform this which vary in their advantages (e.g. evenness of amplification) and disadvantages (e.g. coverage across the genome, scalability, costs), and even in applicability to neurons, as in the case of single-cell trichannel processing which requires dividing cells^43^. In this work, we have directly compared three existing technologies, the well-established PicoPLEX and two very recently developed adaptations of phi-29-based isothermal amplification, PTA and dMDA, using human *post-mortem* brain samples, and highlighted their characteristics. Furthermore, we have updated the popular single-cell CNV caller Gingko for hg38, and for the first time assessed the advantage of utilizing alternative reference genomes besides hg19 or hg38. Indeed, the T2T-CHM13 genome seems to offer potential novel CNV candidates, although the intrinsically low mappability of the novel regions complicates detection. Thus, this study provides key insights for experimental considerations for whole genome single-cell studies and analysis with clear recommendations.

The consistency of amplification across the genome by PicoPLEX makes it preferable for CNV calling, although PTA can also be used. The latter has the advantage of more complete genome coverage, albeit at the expense of requiring a larger bin size to allow CNV calling when a strict MAD cutoff is used. dMDA was not suitable for CNV calling by read depth, but further developments (e.g., more stringent lysis and increased number of droplets per reaction) could improve this. Furthermore, the long MDA amplicons make it potentially well suited for long reads and direct detection of SV breakpoints. This will require effective filtering of pervasive chimeras, with tools already becoming available^44^.

Findings of interest in MSA is the apparent clonality of a gain which encompasses *TLR4*, a possibly disease-relevant gene^39^, as well as tentative support for some CNVs by focused analysis of deep bulk short read WGS. Detection of breakpoints with deep long read WGS and dedicated analysis for somatic SV’s^45^ is needed to fully validate these, although breakpoints of CNVs found in one or a small number of single cells will be impossible to confirm even with sensitive digital PCR methods. Determining the possible relevance of the *TLR4* CNV, and of the apparently high proportion of brain cells with somatic CNVs in MSA (∼40% in the cingulate cortex, and ∼30% in our earlier study of different regions of the same two brains^16^), will require larger studies with well-matched controls. Further single-cell genomic studies in sporadic neurodegenerative disorders are needed to fully elucidate the potential role of somatic mutations in their etiology and pathogenesis.

## Methods

### Human Tissue and cell lines

Fresh frozen *post-mortem* brain samples were provided by the Queen Square Brain Bank, London, UK. All donors had given informed consent for the use of their brain in research and the study was approved by the National Research Ethics service London – Hampstead (10/H0729/21) and from the brain tissue bank by the UK National Research Ethics Service (07/MRE09/72). Samples from 1 control (frontal cortex) and 2 MSA (cingulate cortex) donors were used in this study. We selected MSA cases from which we already have scWGS from other regions^26^. Demographics are presented in **Table 2**. As positive control, we used human skin fibroblasts with a known germline *SNCA* triplication that we previously used with FISH^14^ and scWGS. For PTA, we also used fluorescence-activated nuclei (FANS) sorting NA12878 (B-lymphocyte cell line, RRID:CVCL_7526) single-nuclei provided by BioSkryB as amplification controls.

**Table 2.** Demographics of brains used in the study.

|  | Control | MSA1 | MSA2 |
| --- | --- | --- | --- |
| Age of death (years) | 91 | 50 | 67 |
| Disease duration (years) | NA | 2 | 7 |
| *Post-mortem* interval (hours) | 38 | 30 | 28 |
| Sex | Female | Male | Female |

### Manual isolation of single-nuclei using CellRaft device

We isolated single-nuclei using the CellRaft system (Cell Microsystems) mounted on a Nikon Eclipse TE300 inverted microscope coupled to a CCD camera (KERN optics), as described in detail elsewhere^16,18,46^. Briefly, we prepared nuclear fractions from 30-50 mg frozen tissues or cell pellets, and counterstained nuclei with 1 μg/ml DAPI for 20 min on ice. Then, we seeded 5000 nuclei onto a 10,000-raft array pre-treated with Cell-Tak (Corning), and nuclei were allowed to settle on the raft at 4°C at least overnight. The nuclei were observed under the microscope, and we selected rafts that contained a single-nucleus with a neuronal appearance (large diameter and presence of low condensed chromatin). We isolated individual nuclei of interest manually using a magnetic wand, and subsequently released each nucleus into a 0.2 ml tube containing 5 μl TE or Cell Extraction Buffer for PicoPLEX, 3 μl of Cell Buffer for PTA, and in 2.8 μl lysis buffer (200 mM KOH, 5 mM EDTA pH 8 and 40 mM 1.4 DTT) supplemented with 2 μl dH2O (DNase/RNase-free) for dMDA and kept them on ice until further use. To avoid cross-contamination, we rinsed the wand sequentially with DNase I solution (1x, Corning DLW354242), absolute EtOH and dH2O before and between the individual nuclear collections. In each experiment, we used at least one negative control (a tube with no raft or a raft with no nucleus) and a positive control (15 pg of bulk gDNA).

### Single-cell Whole Genome Amplification (WGA)

We performed single-cell WGA using the following methods:

1. PicoPLEX Single Cell WGA Kit (Takara, R300672 v2 or R300722 v3) according to the manufacturer protocol. In brief, lysis mix was added to the samples and lysis reaction carried out on a thermal cycler. Then pre-amplification reagents were added, and the pre-amplification reaction was carried out on a thermal cycler. Lastly, amplification mix was added. including 1x EvaGreen (Biotium) as reporter dye and the reaction was monitored using qPCR (StepOne, Applied BioSystems). The scWGA products were then cleaned with AMPure XP beads (Beckman Coulter; 1:1 ratio).
2. PTA using ResolveDNA™ Whole Genome Amplification Kit (BioSkryB PN100136) according to the manufacturer protocol. All the PTA reagents were added step by step according to manufactory protocol, but on step 7 the samples were incubated for 20 min at RT, instead of 10 min in a PCR cooler, for improved lysis. The lysed samples with the enzymes and terminators were incubated on thermal cycler at 30°C for 10 h before enzyme deactivation at 65°C for 3 min, followed by bead purification of the amplicons according to the manufacturer protocol.
3. dMDA kit (Samplix) according to the manufacturer protocol, but with an addition of a lysis step 95°C for 3 min followed by 10 min cool down at RT) after alkaline lysis of the samples ^46^. Briefly, samples underwent alkaline and heat lysis, followed by droplet generation in the Xdrop^TM^ instrument (Samplix) which encapsulated the single-cell DNA fragments and dMDA enzyme mix. The droplets were incubated in a thermal block for 16 h on 30°C before inactivating the enzyme at 65°C for 10 min. Droplets were then broken by the addition of break solution and color reagents.

All amplified samples were assessed using Qubit dsDNA BR or HS Assay kits (Thermo Fisher Scientific) and TapeStation (Agilent) using HS D5000 DNA Tapes or HS D1000 DNA Tapes (Agilent). All amplicons were stored at −20°C and quantified by Qubit prior to library preparation.

### Library preparation and short-read sequencing

Unless otherwise indicated, scWGA products were manually fragmented using SureSelect XT HS Enzymatic Fragmentation Kit (Agilent), and libraries for Illumina sequencing were created using SureSelect XT HS2 DNA Reagent Kit (Agilent) manually or using automation with Agilent Bravo, according to manufacturer’s guidelines (dx.doi.org/10.17504/protocols.io.x54v9p3qzg3e/v1) as used without fragmentation to create libraries using ResolveDNA Library Preparation Kit (BioSkryB) kit, according to BioSkryB guidelines. Each library was quantified by Qubit dsDNA BR or HS Assay kits and assessed by TapeStation using D1000 or HS D1000 DNA tapes (Agilent). was determined using Qubit dsDNA HS Assay Kit, HS D1000 tapes on TapeStation, and qPCR (QuantaBio qPCR Library Quantification or NEBNext Library Quant Kit for Illumina). The pooled libraries were sequenced on NextSeq 2000 (100 or 200 cycles, Illumina) or NovaSeq SP v1.5 (300 Cycles, Illumina) using paired-end configuration and including 2% PhiX. For three cells (one dMDA, two PTA), library preparation was repeated and each one sequenced separately, but the bam files were merged for *eq*, *Ginkgo*, and *Copykit*.

## Bioinformatic analyses

### Read alignment and alignment summary

The sequencing reads from PicoPLEX included a 14 base amplification adapter, and the resulting fastq files were therefore trimmed using Trimmomatic-v.0.36 (RRID:SCR_011848; http://www.usadellab.org/cms/?page=trimmomatic)^47^. The data were then aligned to hg38 and T2T-CHM13-v2.0 reference genome with *bowtie2-v2.5.1 (RRID:SCR_016368;* http://bowtie-bio.sourceforge.net/bowtie2/index.shtml)^48^. The sequencing data were then sorted using *Samtools-v.1.1*4 (RRID:SCR_002105; http://www.htslib.org)^49^ and duplicates were marked and coverage metrics collected using *Picard-v2.18.4 (RRID:SCR_006525;* https://broadinstitute.github.io/picard)^50^. BAM files were converted to bed files using *Bedtools v2.25.0* (RRID:SCR_006646; https://bedtools.readthedocs.io/en/latest)^51^ Liftover of T2T-CHM13 to hg38 was performed using *LevioSAM2-v0.2.2*^35^ after which duplicates were marked again. *Preseq-v.3.1.1* (RRID:SCR_018664; http://smithlabresearch.org/software/preseq) was run on bed files across different genome assemblies using “gc_extrap” option^30^. The alignment statistics of all mapping files were generated using *Samtools-v1.14* (RRID:SCR_002105; http://www.htslib.org)^49^ (see code availability).

### Data quality metrics for each amplification method

Data quality assessment and CNV calling were performed using the command-line version of *Ginkgo* (https://github.com/robertaboukhalil/ginkgo). To allow use of *Ginkgo* beyond the hg19 genome, we generated *v*ariable and constant sized bins for hg38 and T2T-CHM13 using the buildGenome scripts provided within *Ginkgo* (https://github.com/robertaboukhalil/ginkgo/tree/master/genomes/scripts), adapted to run on the compute infrastructure of the Flemish Supercomputing Center. The data quality for each amplification method was assessed by calculating MAD, GC content and Lorenz curve with a variable bin size initially of 500 kb for the hg38 genome. We adapted and ran functions within *Ginkgo* to compute locally these three statistics across the autosomes. MAD was calculated between neighboring bins using normalized read counts (the count of each bin divided by the mean read count per cell). As a robust statistic, MAD is resilient to abrupt changes in read counts resulting from copy number changes. The confidence score was calculated as before. GC extreme regions could cause read dropout, one of the reasons for uneven coverage. To model the relationship between normalized read counts and GC content, *Ginkgo* uses the ‘LOWESS’ function in R (RRID:SCR_001905; https://www.r-project.org)^52^, followed by scaling the read counts accordingly. We also computed Lorenz curves as indicators of uniform amplification for each amplification method. The Lorenz curves and GC content plots were visually assessed to compare the data quality of PicoPLEX, PTA and dMDA.

### Denoising

To reduce high within-cell variation in dMDA data, we applied a principal component analysis (PCA) based denoising approach. The approach is based on the idea that PCA can capture common variation in read depth across cells that are mostly likely caused by technical artifacts. To remove the effect of common variation, multiple regression is applied followed by PCA. CNV calling can then be performed using the residuals from the regression, instead of using normalized read counts. Somatic CNVs that are randomly distributed in the genome should remain unaffected^33^. We ran *Ginkgo* for different bin sizes (500 kb, 1 Mb, 2.5 Mb, 5 Mb), and for each bin size, we removed common variation, starting from 40% and increasing up to 90% in 10% increments.

### CNV calling using Ginkgo

*Ginkgo* was used for CNV calling across different genome assemblies^6^. As read lengths varied in some sequencing experiments, the read length of each dataset was rounded down to the next smallest value to get more conservative mappability. *Ginkgo* settings are presented in **Table 3** CNV calling. *Ginkgo* employs circular binary segmentation (CBS), which is implemented in *DNAcopy* in R^53^. *DNAcopy* (v.1.68.0) (RRID:SCR_012560; http://www.bioconductor.org/packages/2.12/bioc/html/DNAcopy.html) was run with *Ginkgo*-implemented parameters which are: alpha=0.01, min.width=5. CBS uses a permutation reference distribution to identify the change points^54^. To ensure reproducibility, all codes were run with *set.seed (1)* in R. Independent segmentation was used throughout. We removed all CNVs smaller than 5 bins.

**Table 3.** Read length and corresponding *Ginkgo* settings.

| scWGA method | Read length used for alignment | *Ginkgo* read length setting |
| --- | --- | --- |
| PTA | 58 bp | 48 bp |
| PTA | 111 bp | 101 bp |
| PicoPLEX | 97 bp | 76 bp |
| PicoPLEX | 137 bp | 101 bp |
| dMDA | 111 bp | 101 bp |

To identify common CNVs shared among multiple cells, we considered two CNVs as shared if their start and end positions are within 2.5 Mb of each other for PicoPLEX cells, and within 5 Mb for PTA cells. We removed these calls from downstream analyses, as they could result from either dry lab/wet lab artifacts or germline mutations. ^33,55^

### Filtering of Ginkgo CNV calls

The output of the segmentation algorithm is integer-like CN values such as 1.2 and 2.1. These deviations from integer values are due to variations in the data caused by both biological and technical factors^6,7^. Therefore, a CN threshold is required to stringently identify gains and losses. For this, we calculated the median CN of diploid segments across autosomes using PicoPLEX data, as the number of cells passing QC (MAD ≤0.3 and confidence score ≥0.7) is greater than that in the PTA data set (25 v 14). The lower limit for the segment size was set to 5, consistent with our use of a minimum of 5 bins for CNV detection, and the upper limit at the size of the smallest chromosome (*n*=127 bins)^7^. We plotted the distribution of the segments CN values on a histogram, from which we set thresholds of 1.29 for a loss and 2.80 for a gain (**Supplementary Fig. 7**).

### CNV calling using Copykit

The newly developed algorithm, *Copykit*^37^ was used for comparison against filtered *Ginkgo* CNV calls in all cells passing QC. For consistency with *Ginkgo* parameters used, Alpha was adjusted to 0.01, and the bin size used for PicoPLEX and PTA amplified cells was 220 kb (as 250 kb is not available) and 500 kb, respectively. *Copykit* is compatible with the genome assemblies hg19 and hg38, therefore CNV calling was performed on hg38 and liftover (not T2T-CHM13). CNV coordinates were extracted from *Copykit* using a custom R script.

### Short read Illumina bulk WGS analysis

Illumina reads for DNA extracted from both MSA brains were mapped to hg38 using *bwa mem (v.0.7.17-r1188) (RRID:SCR_010910;* http://bio-bwa.sourceforge.net) with default parameters including -M to mark split reads as secondary alignments. To visualize possible read support for single-cell gains and losses in bulk Illumina WGS data, we used *Samplot (v.1.3.0*)^41^, which creates images that display the read depth and sequence alignments across specified regions, and reveal any reads supporting a specified CNV. The coordinates of all filtered CNVs (gains and losses) were input into *Samplot*, to detect and reads supporting them in all MSA samples available (2 brain regions from MSA1, 4 regions from MSA2). High-coverage (250x) HG002 data was used for comparison. We allowed 250 kb on either side of each reported breakpoint, due to the inherent inaccuracy of defining CNVs using large bin sizes.

### T2T-CHM13 specific gains

To identify the characteristics of T2T-CHM13 specific gains, we used samples that were aligned to T2T-CHM13 and lifted over to hg38, passing both the MAD (≤0.3) and confidence score (≥0.7) filters. One PicoPLEX cell (“A14_v2_Exp9_1_sn1_P70_06_CC_S14_R”) was excluded from the analyses due to the absence of a T2T-CHM13 version. The number of cells for PicoPLEX is 25 and for PTA, it’s 14. bedtools subtract -v (v2.30.0)^51^ was used to identify T2T-CHM13 specific gains compared to the lifted-over version. T2T-CHM13 unique regions in comparison to hg38 were downloaded from the UCSC Table Browser (http://genome.ucsc.edu/cgi-bin/hgTables, track: CHM13 unique, table: hub_3671779_hgUniqueHg38 on 2023-06-09)^56^. If the T2T-CHM13 specific gains for each cell showed a minimum of 50% overlap with the bed file obtained from the UCSC Table Browser, those positions were kept for the analysis (using Bedtools v2.30.0 with the intersect -f 0.50 -wo).

### Gene Ontology (GO) Analysis

A list of genes covered by significant CNVs was separately identified for MSA patients and controls using the Ensembl BioMart package (RRID:SCR_010714; http://www.ensembl.org/biomart/martview)^57^. The gene lists were submitted to PANTHER (RRID:SCR_004869; http://www.pantherdb.org)^36^ to identify statistical overrepresentation in any GO categories. The remaining analyses were conducted as described^16,18^.

### Statistics and reproducibility

Statistical tests were performed, and graphs were plotted using GraphPad Prism version 10 (RRID:SCR_002798, https://www.graphpad.com/features) and RStudio IDE (v.2023.6.1.524) (RRID:SCR_000432; https://posit.co). Normal distribution was assessed using the D’Agostino & Pearson test. Data are presented as mean ± standard deviation (SD) and a p-value of <0.05 was considered statistically significant. Comparisons between groups were assessed using Kruskal-Wallis test with Dunn’s multiple comparisons test, Brown-Forsythe and Welch ANOVA tests with Dunnett’s T3 multiple comparisons test, paired-matched Friedman test with Dunn’s multiple comparisons test, Mann–Whitney, unpaired student’s t-test with Welch’s correction, Geisser-Greenhouse correction and Tukey’s multiple comparisons test or RM one-way ANOVA with the Geisser-Greenhouse correction and Tukey’s multiple comparisons test, as indicated in the Figure legends. All statistical tests were two-sided.

## Data availability

Sequence data has been deposited at the European Genome-phenome Archive (EGA), which is hosted by the EBI and the CRG, under accession number EGAS50000000020 (Dataset ID: EGAD50000000030). Further information about EGA can be found on https://ega-archive.org “The European Genome-phenome Archive in 2021” (https://academic.oup.com/nar/advance-article/doi/10.1093/nar/gkab1059/6430505). Sample information and statistics related to analyzed cells can be found in **Supplementary Table 4**. Illumina bulk data from the MSA brains will be submitted to an appropriate repository as part of a larger MSA study. The *Ginkgo* bins for hg38 and T2T-CHM13 are available at Zenodo (RRID: SCR_00412, https://doi.org/10.5281/zenodo.8225214).

## Code availability

Custom scripts written in bash (v.5.1.16) or R (v.4.1.2) using *RStudio IDE (v.2023.6.1.524)* (RRID:SCR_000432, https://rstudio.com) are available on https://github.com/zgturan/scWGA_comp.git (https://zenodo.org/doi/10.5281/zenodo.10019431). Custom scripts for re-alignment and alignment analysis are available on https://github.com/srbehera/Brain_scWGA_comparision.

## IP rights notice

For the purpose of open access, the author has applied a CC-BY public copyright license to the Author Accepted Manuscript (AAM) version arising from this submission.

## Author contributions

CP and FJS planned and supervised the study. Single cell wet lab work was performed by EKE with assistance from DRP. Bioinformatic analysis was done by ZGT with input from IB, SB, CM, JD, FJS, CP. Statistical analysis was performed by EKE and ZGT. Ginkgo adaptation was performed by JD. Tissue selection and provision was performed by ZJ. Bulk WGS was provided by SWS. The first draft was written by EKE, ZGT, CP. All authors contributed to and approved the final manuscript.

## Funding and acknowledgements

The study is funded partly by the joint efforts of The Michael J. Fox Foundation for Parkinson’s Research (MJFF) and the Aligning Science Across Parkinson’s (ASAP) initiative. MJFF administers the grant Grant ID 000430] on behalf of ASAP and itself. SB and FS were also supported by NIH (1UG3NS132105-01,1U01HG011758-01). This research was supported in part by the Intramural Research Program of the National Institutes of Health (National Institute of Neurological Disorders and Stroke; project number: 1ZIANS003154). This research was supported in part by the MSA Trust to CP. CGM is supported by a PhD Fellowship from the MSA Trust.

We would like to thank all individuals who donated their brains for research, Dr Darlan Mintussi for help with *CopyKit*, Dr Jan-Willem Taanman for providing fibroblasts, Thorarrin Blondal for assistance in optimizing dMDA and UCL Genomics for sequencing support.

## Supplementary information

**Supplementary Fig. 1.**
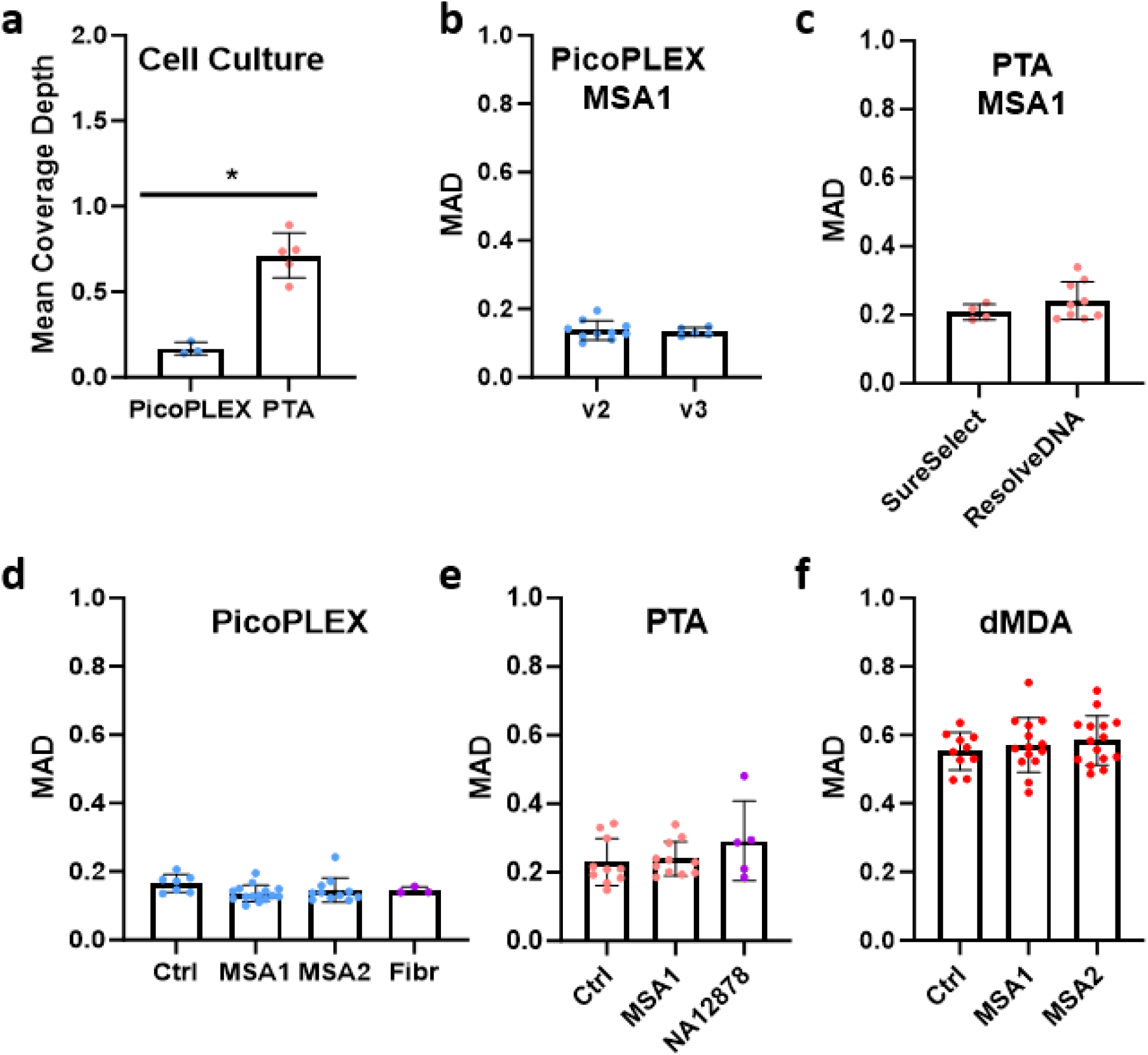
Sequencing coverage of non-brain samples and detailed MAD comparison between samples and method variations. **a** Mean coverage depth distribution of nuclei from cell culture amplified by either PicoPLEX (fibroblasts with known *SNCA* triplication isolated by CellRaft, *n*=3) and PTA (NA12878 cells, isolated by nuclei sorting, *n*=5) *; *p*=0.04 unpaired Mann-Witney test. **b-f** Median Absolute Deviation (MAD scores) of nuclei aligned to hg38 at 500 kb bins. **b** MAD scores PicoPLEX amplified nuclei from MSA1 donor amplified by either PicoPLEX v2 (Takara R300672, *n*=10) or PicoPLEX v3 (Takara R300722, *n*=5) analyzed by unpaired Mann-Whitney test. v2 vs v3 ns (*n*=0.95). **c** MAD from PTA-amplified brain nuclei when used either SureSelect (Agilent) or ResolveDNA (BioSkryB) libraries analyzed by unpaired Mann-Whitney test. SureSelect vs ResolveDNA ns (*n*=0.33). **d** PicoPLEX-amplified nuclei from brain donors (Ctrl *n*=7, MSA1 *n*=15, MSA2 *n*=11) or fibroblast (*n*=3) cells analyzed by Kruskal-Wallis test with Dunn’s multiple comparisons test Ctrl vs MSA1 ns (adj. *p*=0.09), Ctrl vs MSA2 ns (adj. *p*=0.28), Ctrl vs Fibr ns (adj. *p*>0.99), MSA1 vs MSA2 ns (adj. *p*>0.99), MSA1 vs Fibr ns (adj. *p*>0.99), MSA2 vs Fibr ns (adj. *p*>0.99). **e** PTA-amplified nuclei from different brain donors (Ctrl *n*=10), MSA1 *n*=11) or NA12878 cells (*n*=5) analysed by Kruskal-Wallis test with Dunn’s multiple comparisons test. Ctrl vs MSA1 ns (adj. *p*>0.99), Ctrl vs NA12878 ns (adj. *p*=0.88), MSA1 vs NA12878 ns (adj *p*>0.99). **f** MAD of dMDA-amplified from different brain donors (Ctrl *n*=10), MSA1 *n*=14, MSA2 *n*=15) analysed by Brown-Forsythe and Welch ANOVA tests with Dunnett’s T3 multiple comparisons tests. All comparisons ns (adj. p>0.99). Data represent Mean ± SD.

**Supplementary Fig. 2.**
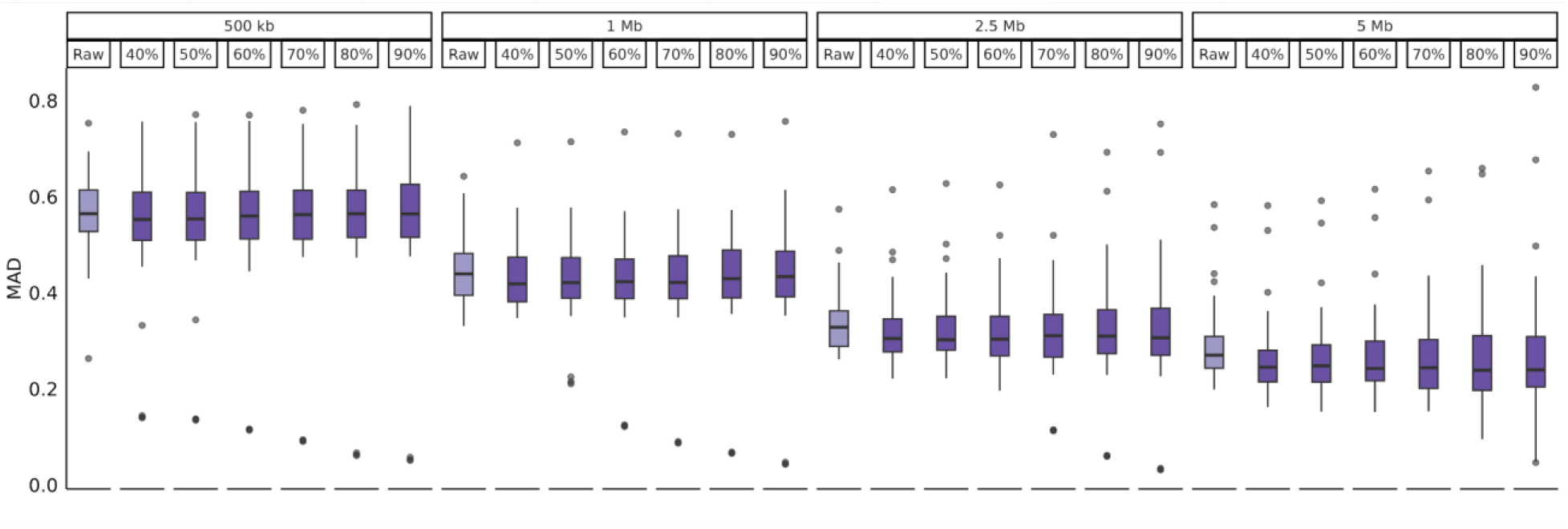
MAD of dMDA at different bin sizes. The results are shown before denoising (“raw”) and after denoising at different proportions. For each bin size, we tested the difference between the raw data and after denoising (40%) using the Wilcoxon test with “paired=TRUE” and alternative=“greater”. For all bin sizes, except for 500 kb (*p=*0.06), we found that 40% denoising significantly improved the MAD score: for bin sizes 1 Mb, 2.5 Mb, and 5 Mb, the *p*-value was < 0.01.

**Supplementary Fig. 3.**
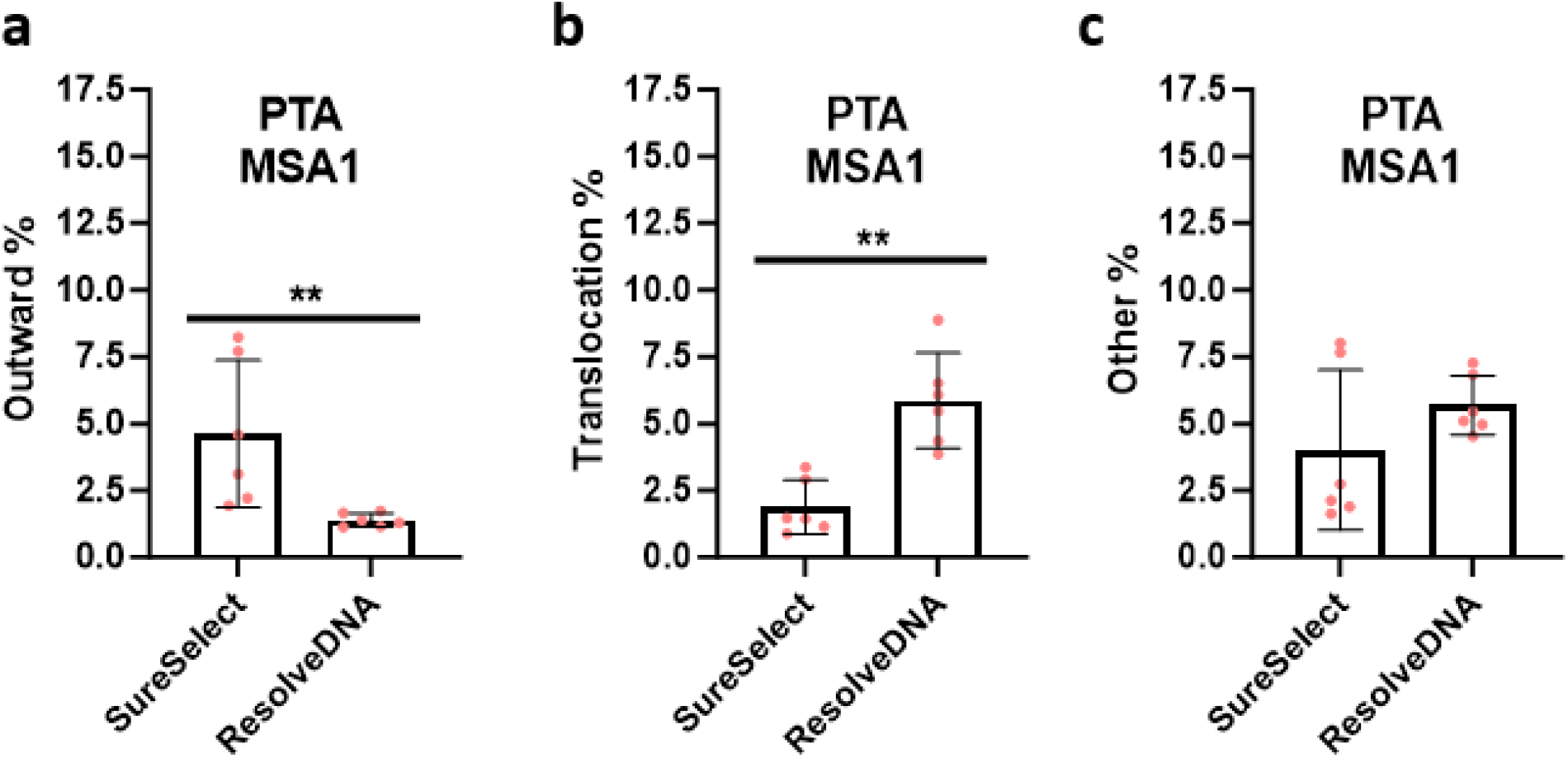
Effect of different library preparations for PTA samples on discordant read pair orientation. PTA-amplified cells from MSA1 brain sequenced before either SureSelect (*n*=6) or ResolveDNA (*n*=6) library preparation. Analyzed by Mann–Whitney, unpaired student’s t-test, mean ± SD shown. **a** Outward pairs. SureSelect vs ResolveDNA ** (*p*=0.002). **b** Translocations. SureSelect vs ResolveDNA ** (*p*=0.002). **c** Other. SureSelect vs ResolveDNA ns (*p*=0.39).

**Supplementary Fig. 4.**
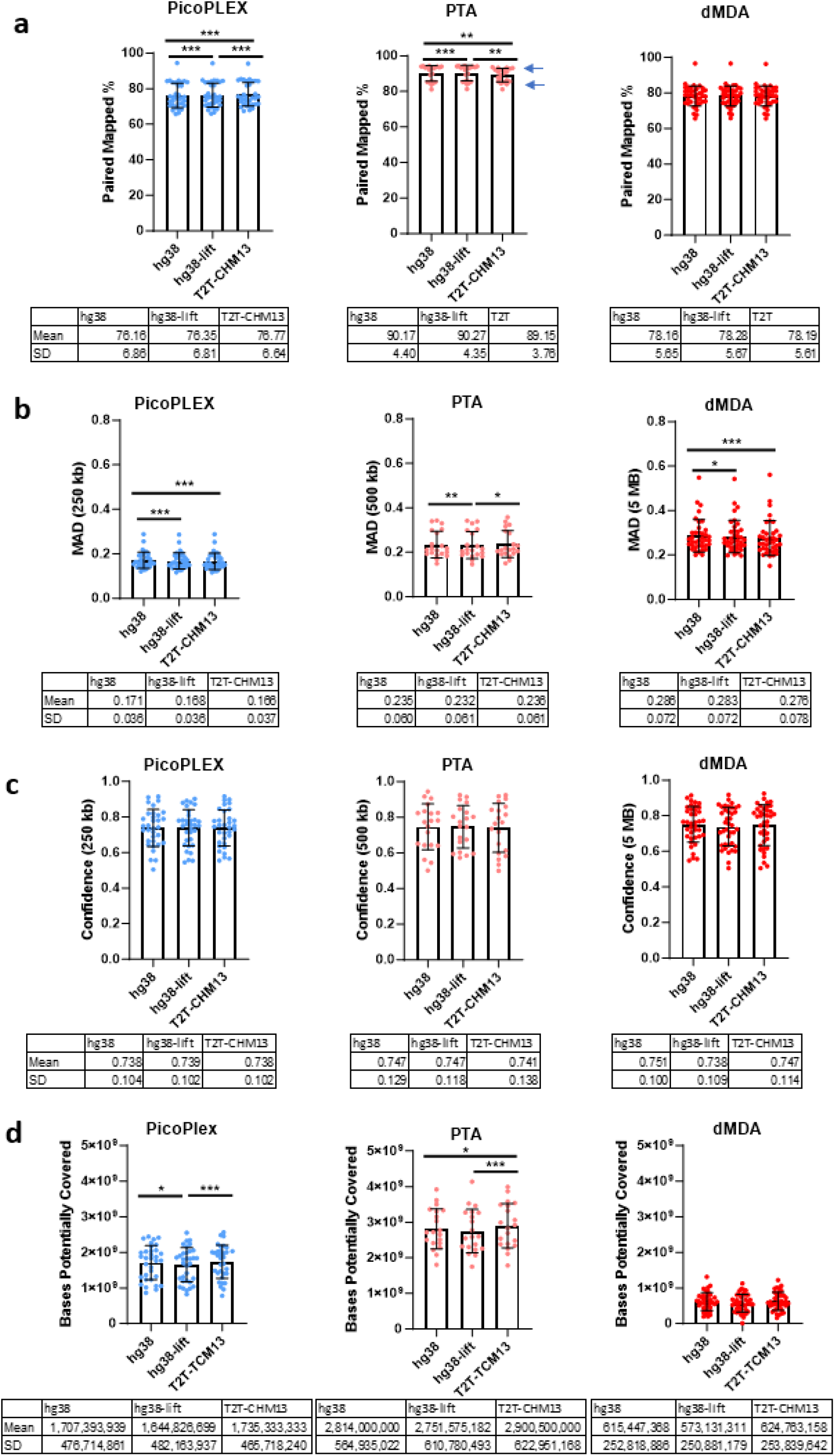
Single-cell metrics across different reference genomes. **a** % of reads aligned. RM one-way ANOVA with the Geisser-Greenhouse correction and Tukey’s multiple comparisons test. PicoPLEX (*n*=33): all comparisons *** (*p*<0.001). PTA (*n*=21): arrows indicate library prep; top=ResolveDNA (n=12), bottom=SureSelect (n=9). hg38 vs hg38-lift ***(*p*<0.001), hg38 vs T2T-CHM23 **(*p*=0.007), hg38-lift vs T2T-CHM23 ** (*p*=0.003). dMDA (*n*=39) : hg38 vs hg38-lift ns (*p*=0.14), hg38 vs T2T-CHM23 ns (*p*=0.27), hg38-lift vs T2T-CHM23 ns (*p*=0.32). **b** MAD. Friedman test with Dunn’s multiple comparisons test. PicoPLEX (n= 33): hg38 vs hg38-lift ***(p<0.001), hg38 vs T2T-CHM23 ***(p<0.001), hg38-lift vs T2T-CHM23 ns(p=0.42). PTA (n=20): hg38 vs hg38-lift **(p=0.005), hg38 vs T2T-CHM23 ns(p>0.99), hg38-lift vs T2T-CHM23 *(p=0.03). dMDA (n= 38): hg38 vs hg38-lift *(p=0.02), hg38 vs T2T-CHM23 ***(p<0.001), hg38-lift vs T2T-CHM23 ns (p=0.26). **c** Confidence scores. RM one-way ANOVA with the Geisser-Greenhouse correction and Tukey’s multiple comparisons test. PicoPLEX (*n*=33): all comparisons ns, hg38 vs hg38-lift (*p*=0.98), hg38 vs T2T-CHM23 (*p*>0.99), hg38-lift vs T2T-CHM23 (*p*=0.95), PTA (*n*=20): all comparisons ns, hg38 vs hg38-lift (*p*>0.99), hg38 vs T2T-CHM23 (*p*=0.85), hg38-lift vs T2T-CHM23 (*p*=0.75), dMDA (n=38): all comparisons ns, hg38 vs hg38-lift (*p*=0.18), hg38 vs T2T-CHM23 (*p*=0.92), hg38-lift vs T2T-CHM23 (*p*=0.77). **d** PreSeq. RM one-way ANOVA with the Geisser-Greenhouse correction and Tukey’s multiple comparisons test. PicoPLEX (*n*=33): hg38 vs hg38-lift *(*p*=0.04), hg38 vs T2T-CHM23 ns(p=0.51), hg38-lift vs T2T-CHM23 ***(p<0.001), PTA (*n*=20): hg38 vs hg38-lift ns (*p*=0.06), hg38 vs T2T-CHM23 *(*p*=0.05), hg38-lift vs T2T-CHM23 ***(*p*<0.001), dMDA (*n*=38): all comparisons ns, hg38 vs hg38-lift (*p*=0.11), hg38 vs T2T-CHM23 (*p*=0.80), hg38-lift vs T2T-CHM23 (*p*=0.10). Data represent Mean ± SD.

**Supplementary Fig. 5.**
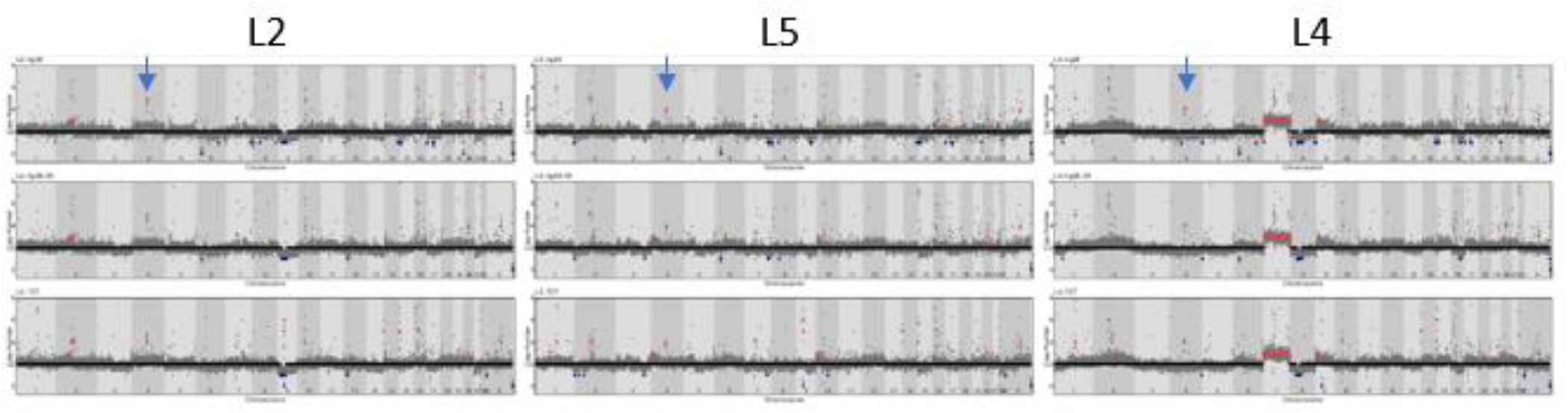
*SNCA* germline triplication (copy number 4) in three single fibroblasts. PicoPLEX WGA, 250 kb bins. Arrow points to CNV. Reference genomes from top to bottom: hg38, liftover, T2T. Note that cell L2 has copy number assigned as 5 in the hg38 and liftover, although visual inspection indicates that the bin copy numbers are between 4 and 5. Cell L4 fails confidence score (0.6). Cell L4 also has a chromosome 7 gain.

**Supplementary Fig. 6.** CN profile of cells passing QC. Each page shows the profile of one cell, including MAD, confidence score, bin size, and CNV calls. All CNVs called by *Ginkgo* are shown in the Table above the CN plot. Filtered CNVs are highlighted (pink = gain, blue = loss).

**Supplementary Fig. 7.**
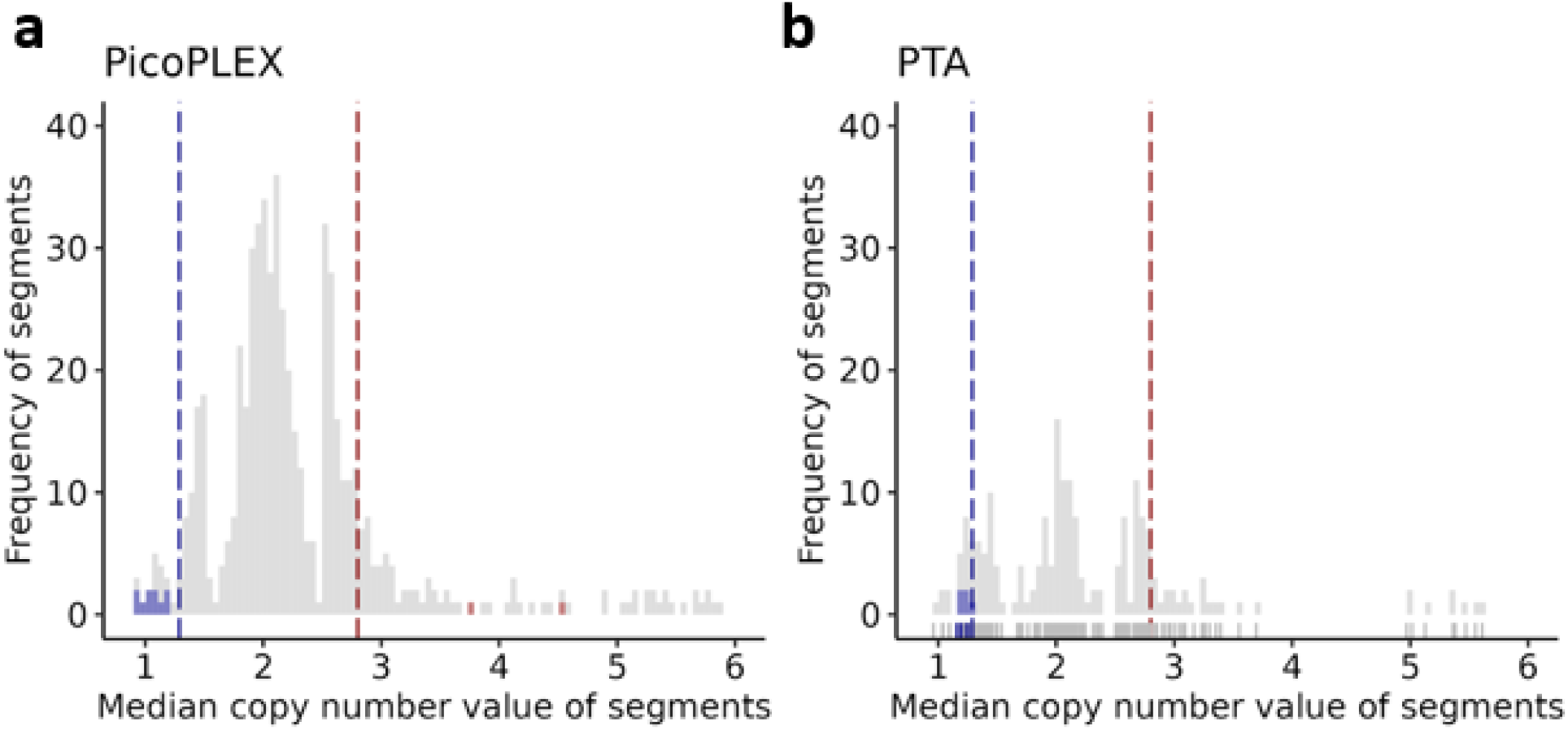
Determination of thresholds for CNV filtering. Plots show the median CN of segments in cells passing QC. A. PicoPLEX. For losses, we set a threshold of 1.29 (dotted purple line), as the median CN of chromosome X was lower than that in all males (*n*=11). For gains, as the CN 2.8 was found in more segments than the immediately higher and lower values, we used that as the cut-off (dotted red line). The two segments in red are the *SNCA* CNV in the fibroblasts passing QC. B. PTA. As the data were sparse, we applied the same thresholds as in PicoPLEX, visually shown as purple and red lines. The median CN of chromosome X in males was below 1.29 in all 5 cells.

**Supplementary Fig. 8.**
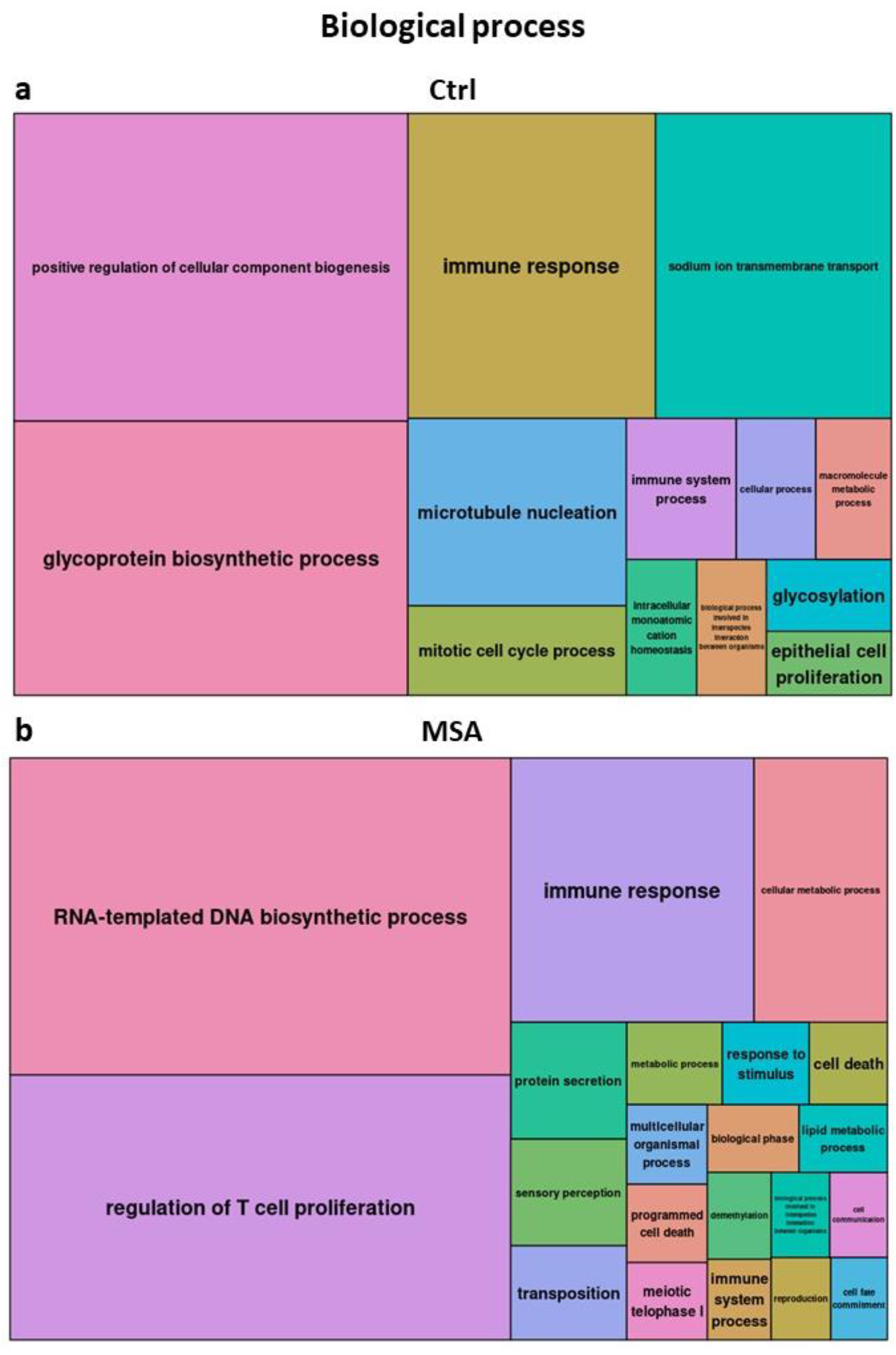
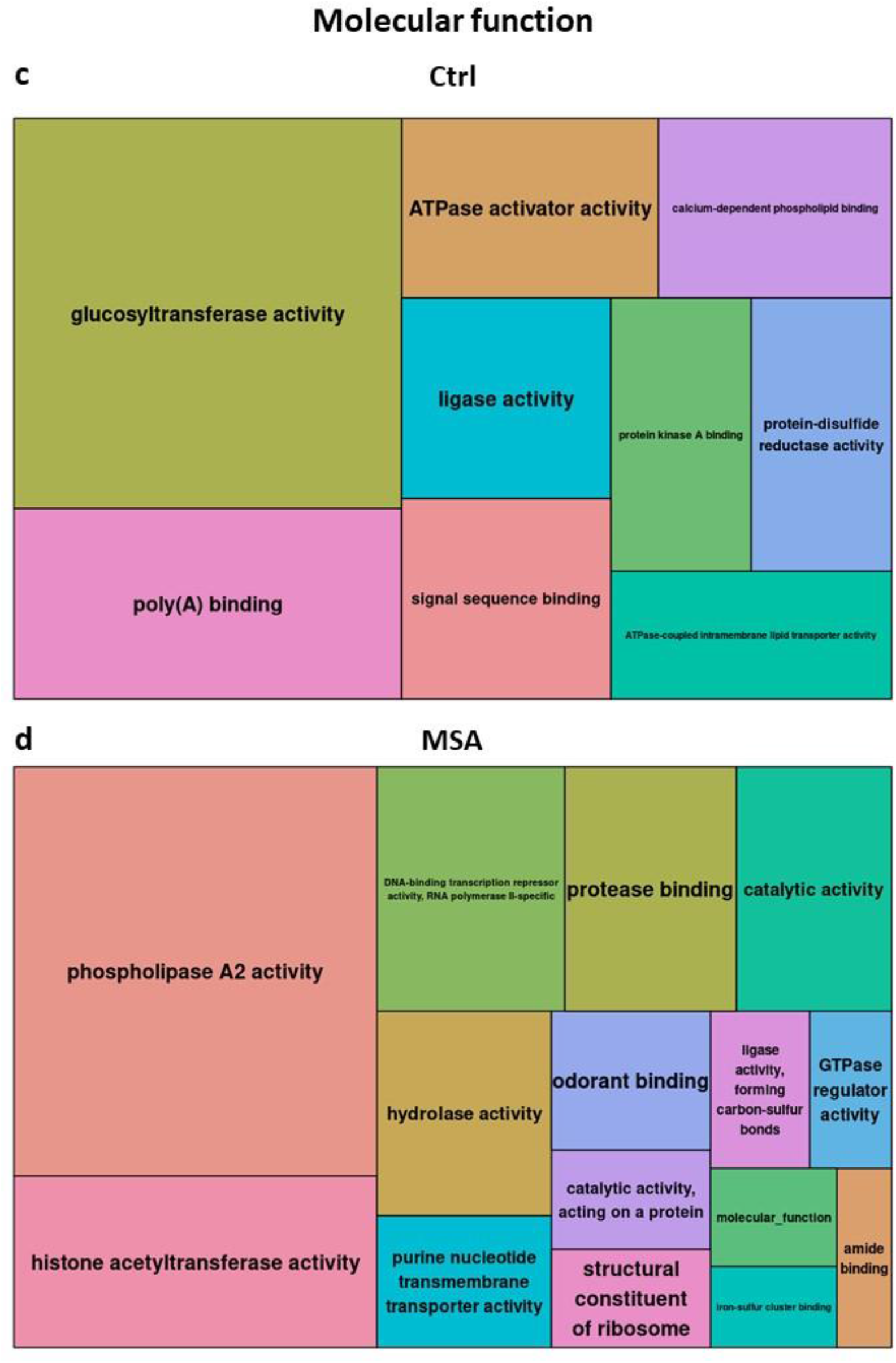
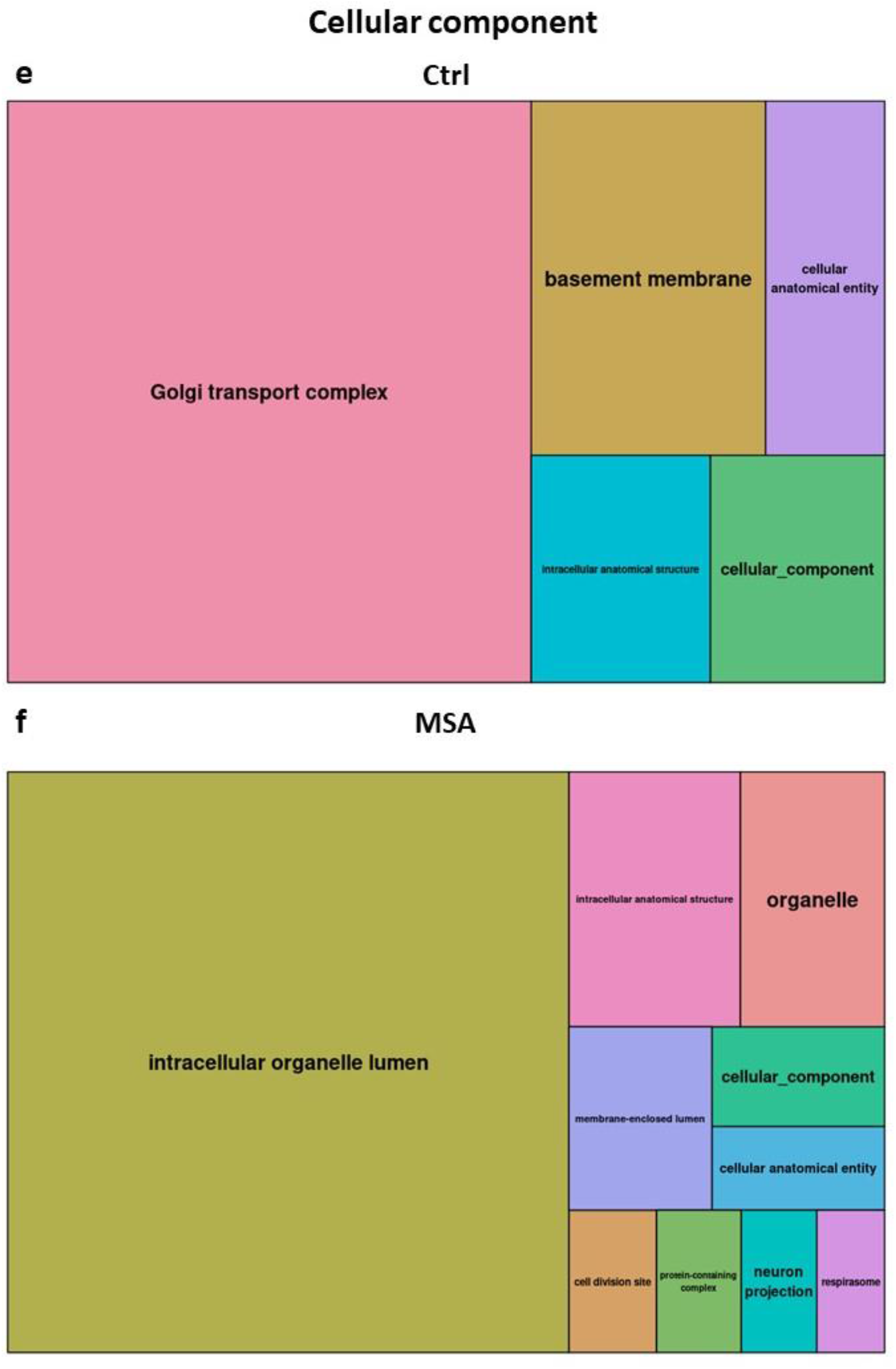
Gene ontology analysis. **a-b** biological process for **a** control and **b** MSA brains, **c-d** molecular function for **c** control and **d** MSA brains. **e-f** cellular component for **e** control and **f** MSA brains.

**Supplementary Fig. 9.**
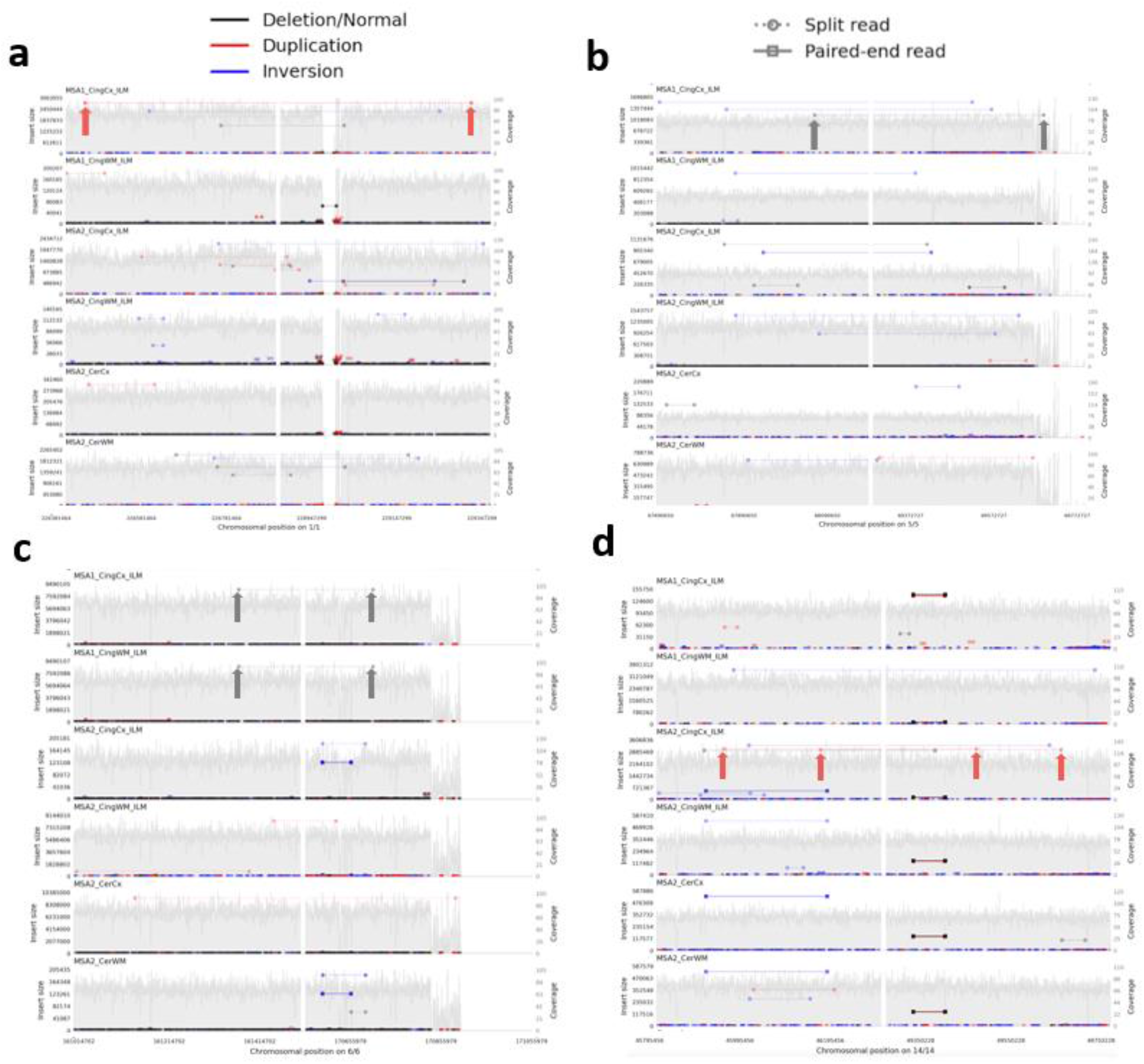
Samplot visualization of bulk Illumina (ILM) WGS in all brain samples focusing on genomic regions of single-cell CNVs supported by bulk reads. All MSA1 and MSA2 brain regions with data available are shown; CingCx= cingulate cortex (also shown in **Fig. 4**), CingCx= cingulate white matter, CerCx= cerebellar cortex, CerWM= cerebellar WM. Read pairs supporting a particular type of CNV / SV indicated according to the scheme at the top, and those supporting each called single-cell CNV arrowed. The left and right panel of each plot show the regions around the reported proximal and distal breakpoint respectively, which is at the middle of each panel, with the chromosomal positions on the x axis below. The y axis indicates the calculated insert size for the read pairs of interest on the left, and the local coverage on the right. Note that each supporting read pair is only seen in the cingulate cortex of the MSA brain where the relevant single-cell was reported, except for **c** which is also shown in the adjacent white matter of the same brain.

### Supplementary Tables

**Supplementary Table 1.** CNV calls (unfiltered) for PicoPLEX and PTA data across different genomes.

|  | Gains / cell | Losses / cell |
| --- | --- | --- |
| hg38 | 12.62 | 11.26 |
| hg38-lift | 14.41 | 4.00 |
| T2T | 16.98 | 3.40 |

**Supplementary Table 2.** Information About Significant CNVs. This shows detailed information about all CNVs which were called after filtering. CN=copy number. Ginkgo results are shown in columns F-L, and Copykit (if called) L-O. The size difference for CNV called by both is shown in P.

**Supplementary Table 3.**
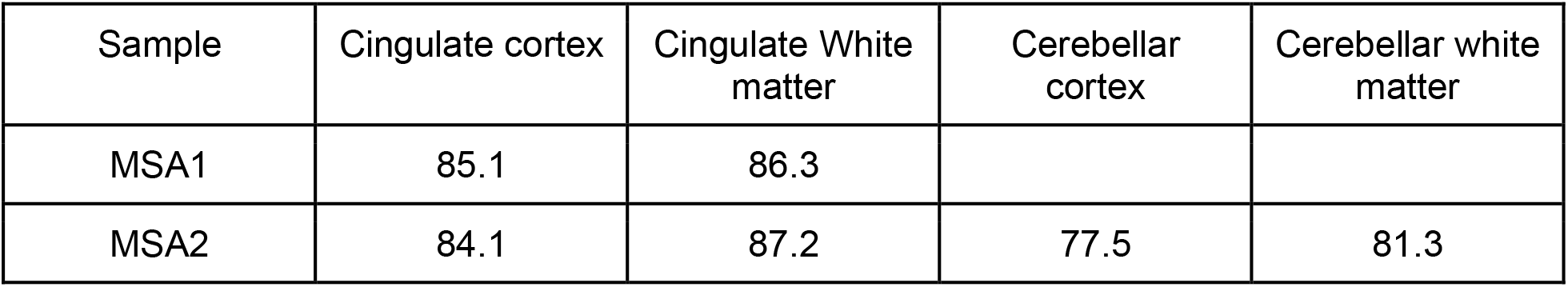
Coverage of bulk WGS.

**Supplementary Table 4.** Sample information submitted in EGA and quality control detailed information. This shows detailed information about the analyzed cells. **a** Nuclei used in this study and detailed information about them. **b-d** Sample statistics about different amplification method, **a** PicoPLEX, **b** PTA, **c** dMDA.

