## Supplementary Figure 6 for "Single-cell somatic copy number variants in brain using different amplification methods and reference genomes"

### PicoPLEX_76bp_250kb_Liftover; MAD= 0.14 Confidence_score= 0.78

| samples | ID | shared_ind_number | chr | cn | cn_median | start | end | width |
|---|---|---|---|---|---|---|---|---|
| MSA−1 | A11_v3_Exp6.1.sn3 | 4 | chr1 | 5 | 5.44 | 119788580 | 149878253 | 30089674 |
| | | 1 | chr4 | 3 | 2.58 | 102822109 | 105001087 | 2178979 |
| | | 4 | chr9 | 14 | 13.65 | 38640746 | 68419166 | 29778421 |
| | | 4 | chr20 | 4 | 4.41 | 25724654 | 30899236 | 5174583 |
| | | 1 | chrX | 1 | 1.04 | 1 | 89017669 | 89017669 |
| | | 1 | chrX | 1 | 1.17 | 92901881 | 156040895 | 63139015 |

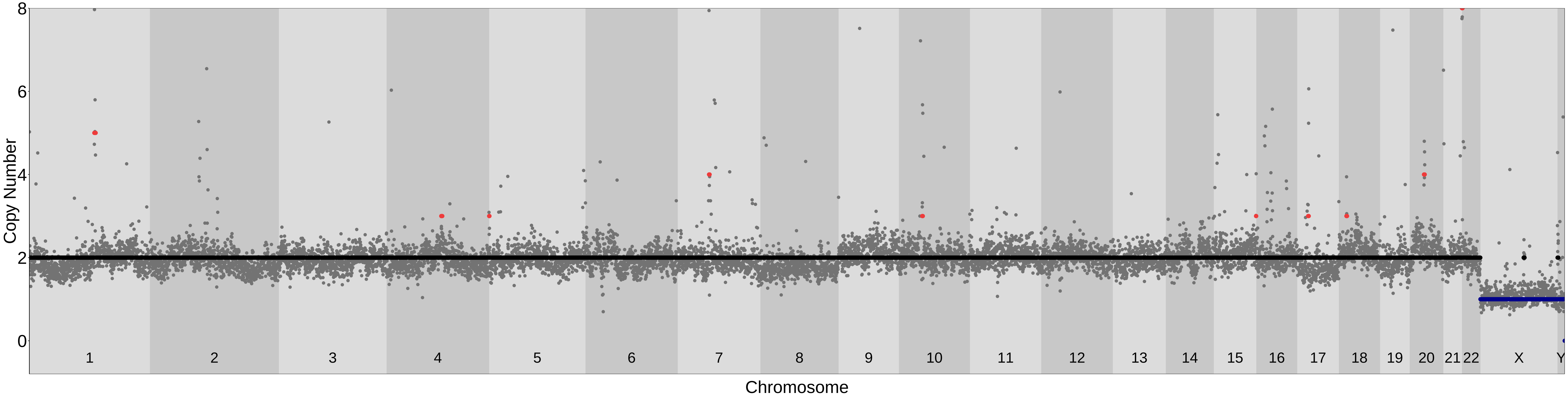

### PicoPLEX_76bp_250kb_Liftover; MAD= 0.16 Confidence_score= 0.72

| samples | ID | shared_ind_number | chr | cn | cn_median | start | end | width |
|---|---|---|---|---|---|---|---|---|
| MSA−1 | A12_v3_Exp6.1_sn4 | 4 | chr1 | 5 | 5.42 | 119788580 | 149878253 | 30089674 |
| | | 1 | chr2 | 1 | 1.45 | 183120714 | 197577018 | 14456305 |
| | | 4 | chr9 | 12 | 11.78 | 38640746 | 68419166 | 29778421 |
| | | 1 | chr9 | 1 | 1.50 | 98583039 | 107807672 | 9224634 |
| | | 1 | chr10 | 3 | 2.54 | 112325062 | 114633716 | 2308655 |
| | | 4 | chr20 | 5 | 4.58 | 25724654 | 30899236 | 5174583 |
| | | 1 | chrX | 1 | 1.16 | 1 | 89017669 | 89017669 |
| | | 1 | chrX | 1 | 1.21 | 93293918 | 156040895 | 62746978 |

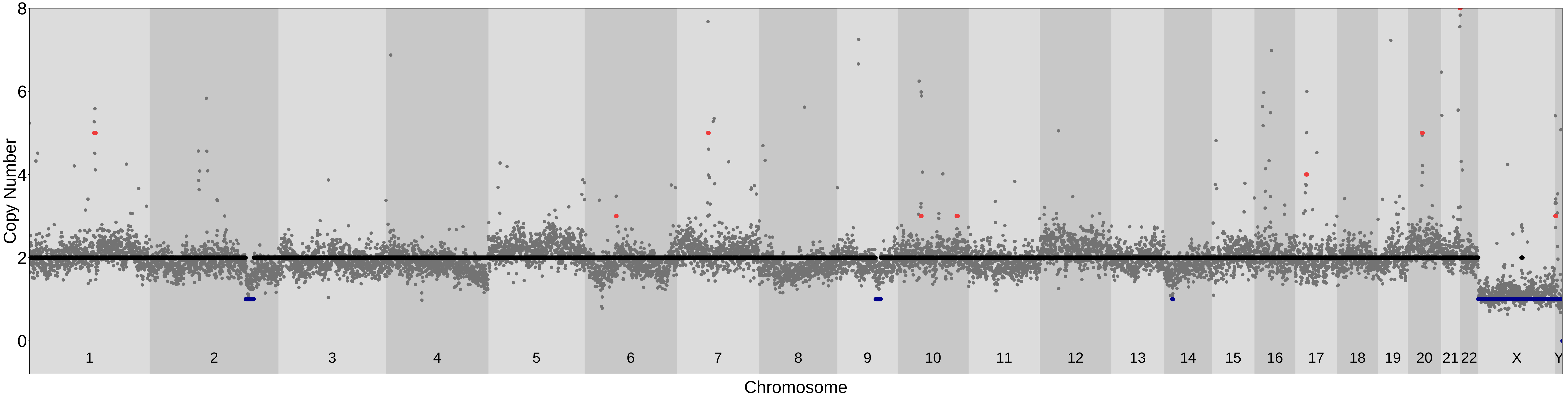

### PicoPLEX_76bp_250kb_Liftover; MAD= 0.18 Confidence_score= 0.75

| samples | ID | shared_ind_number | chr | cn | cn_median | start | end | width |
|---|---|---|---|---|---|---|---|---|
| MSA−1 | A13_v3_Exp6.1_sn6 | 4 | chr1 | 5 | 4.91 | 119788580 | 149878253 | 30089674 |
| | | 4 | chr2 | 4 | 4.28 | 86824353 | 94916453 | 8092101 |
| | | 1 | chr2 | 1 | 1.48 | 202060106 | 203660707 | 1600602 |
| | | 1 | chr6 | 3 | 2.84 | 1 | 2496842 | 2496842 |
| | | 1 | chr6 | 3 | 2.51 | 57492507 | 72138448 | 14645942 |
| | | 1 | chr6 | 3 | 2.55 | 93301029 | 109691169 | 16390141 |
| | | 1 | chr7 | 3 | 2.58 | 63785714 | 77394825 | 13609112 |
| | | 1 | chr8 | 3 | 2.51 | 78533691 | 92973922 | 14440232 |
| | | 1 | chr8 | 3 | 2.58 | 121057404 | 139838730 | 18781327 |
| | | 4 | chr9 | 12 | 12.32 | 38640746 | 68419166 | 29778421 |
| | | 1 | chr11 | 3 | 2.67 | 69978290 | 72070343 | 2092054 |
| | | 1 | chr17 | 3 | 2.86 | 18157522 | 27290069 | 9132548 |
| | | 1 | chr17 | 3 | 2.51 | 72139705 | 81301536 | 9161832 |
| | | 4 | chr20 | 4 | 4.18 | 25724654 | 30899236 | 5174583 |
| | | 1 | chr22 | 3 | 2.50 | 1 | 42675627 | 42675627 |
| | | 1 | chrX | 1 | 1.01 | 1 | 89017669 | 89017669 |
| | | 1 | chrX | 1 | 0.98 | 93293918 | 156040895 | 62746978 |

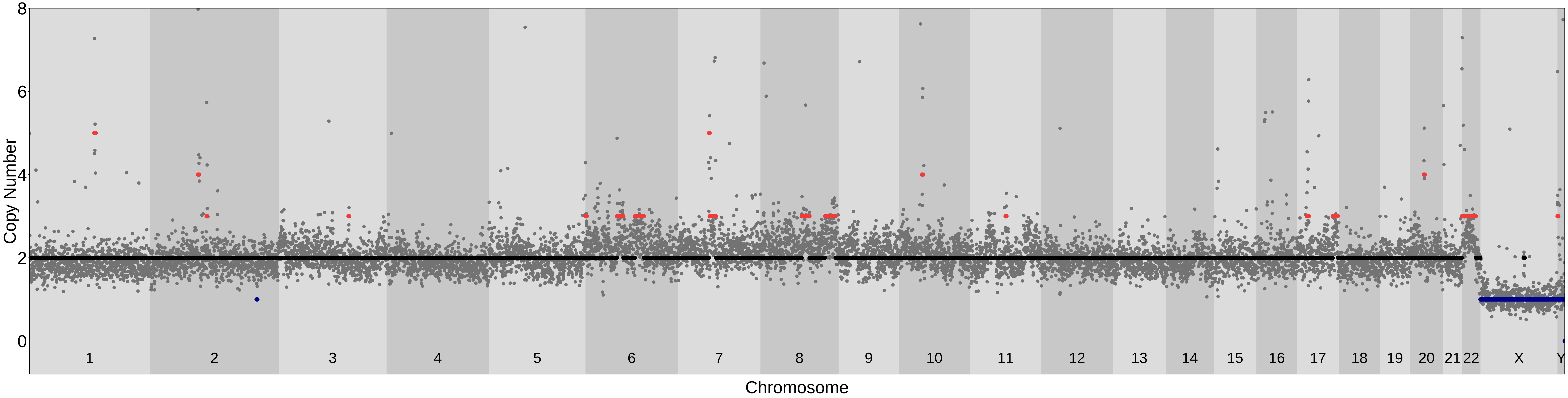

### PicoPLEX_76bp_250kb_Liftover; MAD= 0.13 Confidence_score= 0.74

| samples | ID | shared_ind_number | chr | cn | cn_median | start | end | width |
|---|---|---|---|---|---|---|---|---|
| MSA−1 | A14_v2_Exp9.1_sn1 | 1 | chr1 | 3 | 2.79 | 1 | 2845080 | 2845080 |
| | | 1 | chr1 | 3 | 2.61 | 15082134 | 16926419 | 1844286 |
| | | 4 | chr1 | 6 | 5.74 | 119788580 | 149878253 | 30089674 |
| | | 3 | chr2 | 3 | 2.60 | 129947291 | 132549472 | 2602182 |
| | | 1 | chr5 | 3 | 2.58 | 6893757 | 11603992 | 4710236 |
| | | 1 | chr5 | 3 | 2.60 | 16261610 | 50704308 | 34442699 |
| | | 1 | chr5 | 3 | 2.66 | 67122502 | 73169334 | 6046833 |
| | | 1 | chr5 | 3 | 2.54 | 95598353 | 101626773 | 6028421 |
| | | 1 | chr5 | 3 | 2.53 | 151625606 | 169958689 | 18333084 |
| | | 1 | chr6 | 1 | 1.41 | 61542307 | 63940094 | 2397788 |
| | | 1 | chr7 | 3 | 2.62 | 72569993 | 77394825 | 4824833 |
| | | 4 | chr9 | 12 | 12.38 | 38640746 | 68419166 | 29778421 |
| | | 1 | chr12 | 3 | 2.58 | 47984790 | 52004046 | 4019257 |
| | | 1 | chr14 | 1 | 1.44 | 86567954 | 89183850 | 2615897 |
| | | 1 | chr21 | 3 | 2.65 | 25677414 | 34703974 | 9026561 |
| | | 3 | chr22 | 3 | 3.19 | 1 | 19375084 | 19375084 |
| | | 1 | chrX | 1 | 0.95 | 1 | 89017669 | 89017669 |
| | | 1 | chrX | 1 | 0.94 | 92901881 | 156040895 | 63139015 |

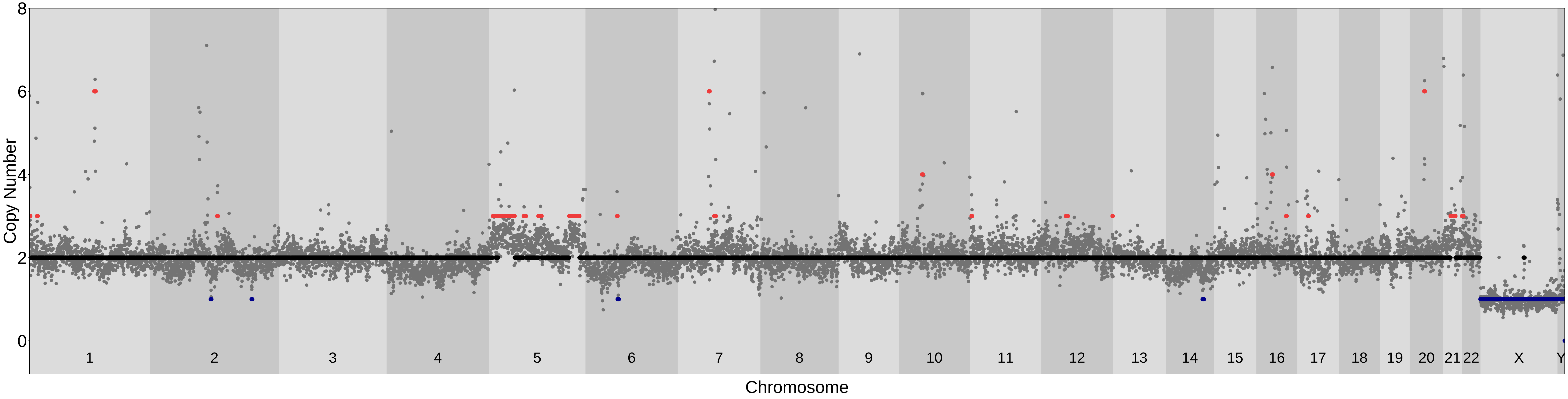

### PicoPLEX_76bp_250kb_Liftover; MAD= 0.15 Confidence_score= 0.75

| samples | ID | shared_ind_number | chr | cn | cn_median | start | end | width |
|---|---|---|---|---|---|---|---|---|
| MSA−1 | A22_v2_Exp9.1_sn3 | 4 | chr1 | 6 | 6.09 | 119788580 | 149878253 | 30089674 |
| | | 1 | chr1 | 3 | 2.64 | 149878254 | 152012333 | 2134080 |
| | | 3 | chr2 | 3 | 3.09 | 129947291 | 132549472 | 2602182 |
| | | 1 | chr2 | 3 | 2.72 | 142985338 | 145563695 | 2578358 |
| | | 1 | chr3 | 3 | 3.01 | 14987425 | 16571696 | 1584272 |
| | | 1 | chr3 | 3 | 2.54 | 48501774 | 58562212 | 10060439 |
| | | 1 | chr3 | 3 | 2.51 | 68484328 | 71099680 | 2615353 |
| | | 1 | chr3 | 3 | 2.61 | 99309312 | 101689574 | 2380263 |
| | | 1 | chr4 | 1 | 1.47 | 107923288 | 111094382 | 3171095 |
| | | 1 | chr5 | 1 | 1.46 | 27421682 | 30046107 | 2624426 |
| | | 1 | chr5 | 3 | 2.51 | 44854060 | 50374296 | 5520237 |
| | | 1 | chr5 | 3 | 2.56 | 136118742 | 139303709 | 3184968 |
| | | 1 | chr5 | 1 | 1.49 | 173612992 | 175688230 | 2075239 |
| | | 2 | chr7 | 3 | 2.59 | 55636895 | 57799139 | 2162245 |
| | | 1 | chr9 | 3 | 2.58 | 15173045 | 33224995 | 18051951 |
| | | 4 | chr9 | 14 | 13.63 | 38640746 | 68419166 | 29778421 |
| | | 2 | chr10 | 3 | 2.91 | 37872529 | 42772360 | 4899832 |
| | | 1 | chr10 | 3 | 2.60 | 125821322 | 127390087 | 1568766 |
| | | 1 | chr13 | 3 | 2.70 | 103696556 | 108341293 | 4644738 |
| | | 1 | chr16 | 3 | 2.51 | 14605263 | 46698329 | 32093067 |
| | | 1 | chr17 | 3 | 3.08 | 74995515 | 76870296 | 1874782 |
| | | 2 | chr21 | 3 | 2.51 | 42743069 | 46709983 | 3966915 |
| | | 1 | chrX | 1 | 0.94 | 1 | 89017669 | 89017669 |
| | | 1 | chrX | 1 | 0.94 | 93293918 | 156040895 | 62746978 |

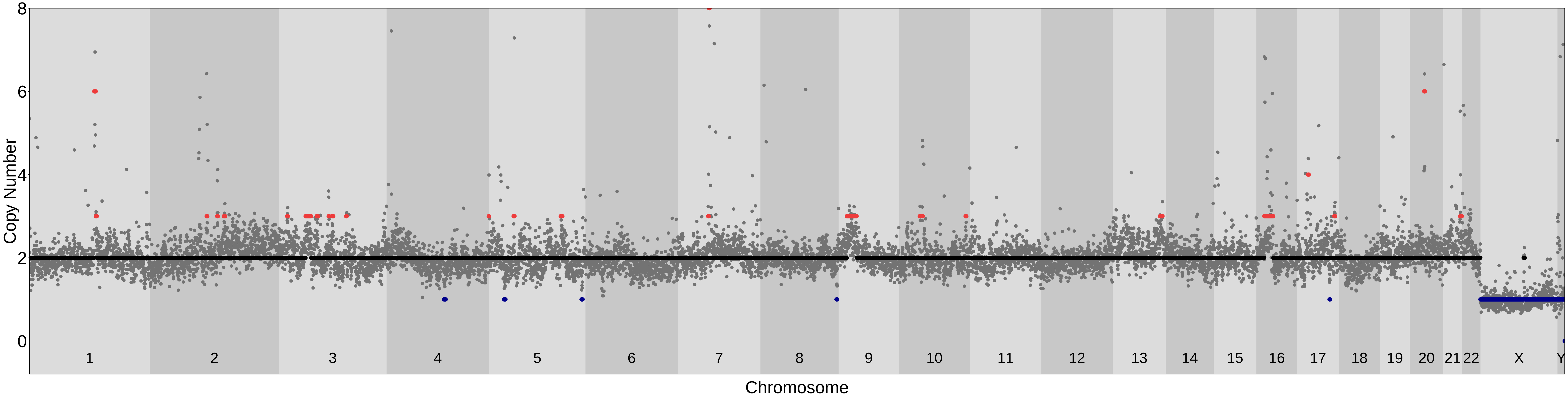

### PicoPLEX_76bp_250kb_Liftover; MAD= 0.14 Confidence_score= 0.76

| samples | ID | shared_ind_number | chr | cn | cn_median | start | end | width |
|---|---|---|---|---|---|---|---|---|
| MSA−1 | A23_v2_Exp9.1_sn4 | 1 | chr1 | 1 | 1.43 | 17192376 | 18985365 | 1792990 |
| | | 4 | chr1 | 5 | 5.37 | 119788580 | 149878253 | 30089674 |
| | | 4 | chr2 | 3 | 2.98 | 86824353 | 97590603 | 10766251 |
| | | 3 | chr2 | 3 | 2.91 | 129947291 | 132549472 | 2602182 |
| | | 1 | chr3 | 1 | 1.49 | 125992138 | 128569150 | 2577013 |
| | | 4 | chr6 | 1 | 1.49 | 30813820 | 34064176 | 3250357 |
| | | 4 | chr6 | 3 | 2.90 | 57145229 | 62604501 | 5459273 |
| | | 4 | chr9 | 13 | 13.48 | 38640746 | 68419166 | 29778421 |
| | | 2 | chr10 | 3 | 3.32 | 37872529 | 42484908 | 4612380 |
| | | 3 | chr11 | 3 | 2.74 | 88690994 | 91347300 | 2656307 |
| | | 1 | chr17 | 3 | 2.50 | 62193911 | 72917441 | 10723531 |
| | | 1 | chrX | 1 | 1.04 | 1 | 89017669 | 89017669 |
| | | 1 | chrX | 1 | 1.07 | 93293918 | 156040895 | 62746978 |

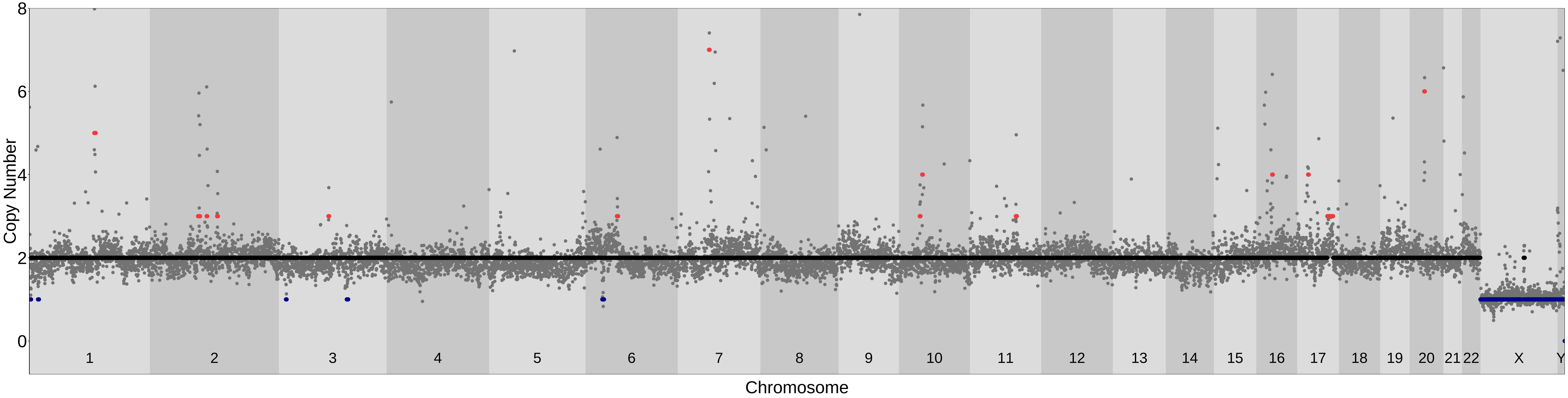

### PicoPLEX_76bp_250kb_Liftover; MAD= 0.22 Confidence_score= 0.72

| samples | ID | shared_ind_number | chr | cn | cn_median | start | end | width |
|---|---|---|---|---|---|---|---|---|
| MSA−1 | A24_v2_Exp9.1_sn5 | 1 | chr1 | 1 | 1.06 | 2471997 | 4133387 | 1661391 |
| | | 1 | chr1 | 1 | 1.41 | 15082134 | 34121854 | 19039721 |
| | | 1 | chr1 | 1 | 1.45 | 44109606 | 54518715 | 10409110 |
| | | 3 | chr1 | 4 | 4.15 | 119251453 | 147189751 | 27938299 |
| | | 1 | chr1 | 3 | 2.93 | 226631464 | 229097299 | 2465836 |
| | | 3 | chr2 | 3 | 3.32 | 129947291 | 132549472 | 2602182 |
| | | 1 | chr3 | 3 | 2.66 | 1 | 14218862 | 14218862 |
| | | 1 | chr3 | 3 | 2.52 | 26068496 | 30036029 | 3967534 |
| | | 1 | chr3 | 3 | 2.79 | 32626622 | 47430118 | 14803497 |
| | | 1 | chr3 | 3 | 2.84 | 51468018 | 75984384 | 24516367 |
| | | 1 | chr3 | 3 | 2.52 | 174041212 | 198295559 | 24254348 |
| | | 1 | chr5 | 1 | 1.15 | 67940650 | 69522727 | 1582078 |
| | | 1 | chr6 | 1 | 1.30 | 1 | 3804287 | 3804287 |
| | | 1 | chr6 | 1 | 1.47 | 17124060 | 22644544 | 5520485 |
| | | 1 | chr6 | 1 | 1.29 | 167731689 | 170805979 | 3074291 |
| | | 3 | chr7 | 8 | 7.83 | 56573819 | 62787680 | 6213862 |
| | | 1 | chr7 | 3 | 2.56 | 62787681 | 73313775 | 10526095 |
| | | 1 | chr7 | 3 | 2.63 | 105676524 | 118297130 | 12620607 |
| | | 1 | chr8 | 3 | 3.15 | 126794482 | 128369357 | 1574876 |
| | | 1 | chr9 | 3 | 2.73 | 12295102 | 21023548 | 8728447 |
| | | 4 | chr9 | 19 | 18.93 | 38640746 | 68419166 | 29778421 |
| | | 2 | chr9 | 3 | 3.41 | 68419167 | 89373829 | 20954663 |
| | | 1 | chr9 | 3 | 2.53 | 90952040 | 102530544 | 11578505 |
| | | 1 | chr9 | 3 | 3.06 | 105969935 | 130296262 | 24326328 |
| | | 1 | chr9 | 4 | 3.55 | 132387545 | 138394717 | 6007173 |
| | | 1 | chr10 | 1 | 1.33 | 67595351 | 73633853 | 6038503 |
| | | 1 | chr11 | 1 | 1.36 | 8994325 | 10592093 | 1597769 |
| | | 1 | chr11 | 1 | 1.48 | 46537233 | 48440665 | 1903433 |
| | | 2 | chr11 | 3 | 2.65 | 48440666 | 55914767 | 7474102 |
| | | 1 | chr11 | 1 | 1.14 | 63511135 | 68422287 | 4911153 |
| | | 1 | chr12 | 1 | 1.21 | 1 | 1637065 | 1637065 |
| | | 1 | chr12 | 1 | 1.43 | 111693075 | 114050522 | 2357448 |
| | | 1 | chr14 | 1 | 1.50 | 90491583 | 95186270 | 4694688 |
| | | 1 | chr15 | 3 | 2.53 | 60090690 | 73701521 | 13610832 |
| | | 1 | chr15 | 4 | 3.89 | 73701522 | 75925597 | 2224076 |
| | | 1 | chr16 | 3 | 2.80 | 1 | 46698329 | 46698329 |
| | | 1 | chr16 | 3 | 2.78 | 58951483 | 61857346 | 2905864 |
| | | 1 | chr16 | 1 | 1.10 | 85026918 | 90338345 | 5311428 |
| | | 1 | chr17 | 3 | 2.70 | 1 | 48689823 | 48689823 |
| | | 1 | chr17 | 1 | 1.46 | 68758181 | 74734069 | 5975889 |
| | | 1 | chr17 | 4 | 3.52 | 74734070 | 78178169 | 3444100 |
| | | 1 | chr17 | 4 | 3.88 | 79994262 | 83257441 | 3263180 |
| | | 1 | chr18 | 1 | 1.44 | 55252983 | 58379931 | 3126949 |
| | | 1 | chr19 | 3 | 2.88 | 1 | 11906078 | 11906078 |
| | | 1 | chr21 | 1 | 1.44 | 44854666 | 46709983 | 1855318 |
| | | 1 | chr22 | 1 | 1.35 | 33361450 | 37555750 | 4194301 |
| | | 1 | chr22 | 3 | 2.84 | 37555751 | 50818468 | 13262718 |
| | | 1 | chrX | 1 | 1.47 | 121000952 | 146707847 | 25706896 |

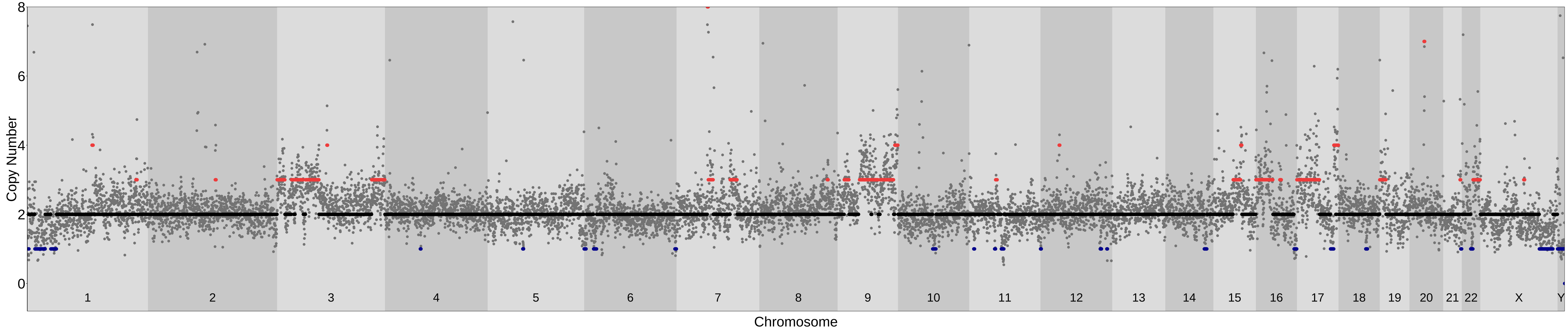

### PicoPLEX_76bp_250kb_Liftover; MAD= 0.16 Confidence_score= 0.72

| samples | ID | shared_ind_number | chr | cn | cn_median | start | end | width |
|---|---|---|---|---|---|---|---|---|
| MSA−1 | A26_v2_Exp9.1_sn7 | 4 | chr1 | 6 | 5.56 | 119788580 | 149878253 | 30089674 |
| | | 1 | chr3 | 3 | 2.55 | 109004922 | 115861487 | 6856566 |
| | | 1 | chr5 | 3 | 2.76 | 134041208 | 138771632 | 4730425 |
| | | 1 | chr5 | 3 | 2.71 | 177244949 | 181538259 | 4293311 |
| | | 1 | chr6 | 3 | 2.75 | 47664390 | 57145228 | 9480839 |
| | | 1 | chr6 | 3 | 2.57 | 61542307 | 67122374 | 5580068 |
| | | 4 | chr9 | 13 | 12.51 | 38640746 | 68419166 | 29778421 |
| | | 2 | chr11 | 3 | 2.71 | 26873455 | 30825636 | 3952182 |
| | | 2 | chr11 | 3 | 3.17 | 48440666 | 54757193 | 6316528 |
| | | 1 | chr11 | 3 | 2.76 | 121496035 | 123335148 | 1839114 |
| | | 1 | chr21 | 1 | 1.50 | 13909163 | 32237669 | 18328507 |
| | | 1 | chrX | 1 | 0.93 | 1 | 89017669 | 89017669 |
| | | 1 | chrX | 1 | 0.96 | 93293918 | 156040895 | 62746978 |

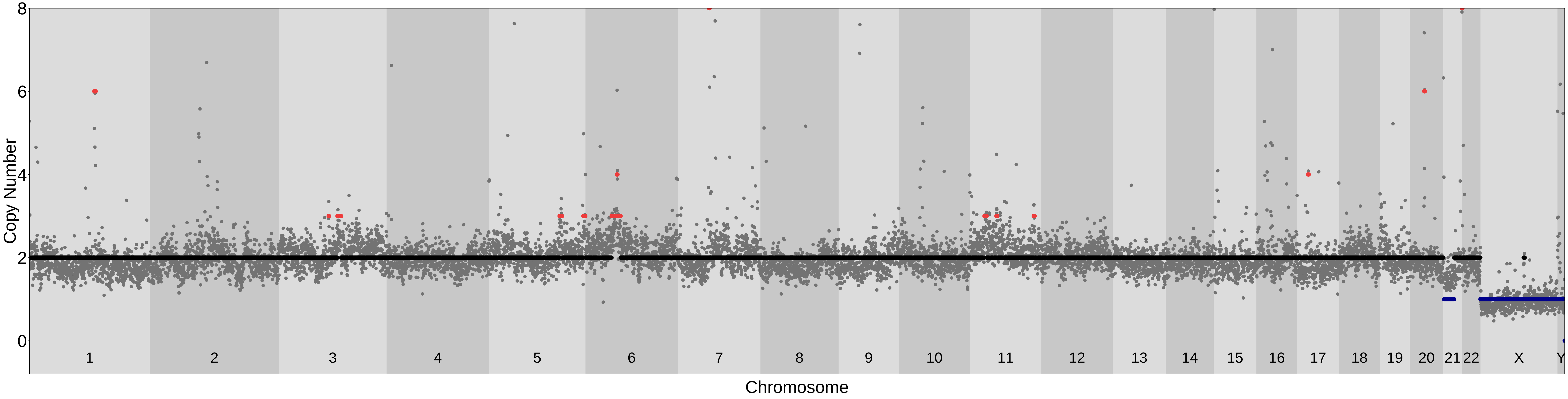

### PicoPLEX_76bp_250kb_Liftover; MAD= 0.15 Confidence_score= 0.77

| samples | ID | shared_ind_number | chr | cn | cn_median | start | end | width |
|---|---|---|---|---|---|---|---|---|
| MSA−1 | A27_v2_Exp9.1_sn8 | 4 | chr1 | 6 | 6.41 | 119788580 | 149878253 | 30089674 |
| | | 1 | chr2 | 3 | 2.52 | 13305039 | 14901752 | 1596714 |
| | | 4 | chr6 | 1 | 1.39 | 30813820 | 33277394 | 2463575 |
| | | 4 | chr9 | 13 | 12.81 | 38640746 | 68419166 | 29778421 |
| | | 2 | chr10 | 3 | 2.80 | 37872529 | 42772360 | 4899832 |
| | | 1 | chrX | 1 | 1.19 | 1 | 89017669 | 89017669 |
| | | 1 | chrX | 1 | 1.23 | 92901881 | 156040895 | 63139015 |

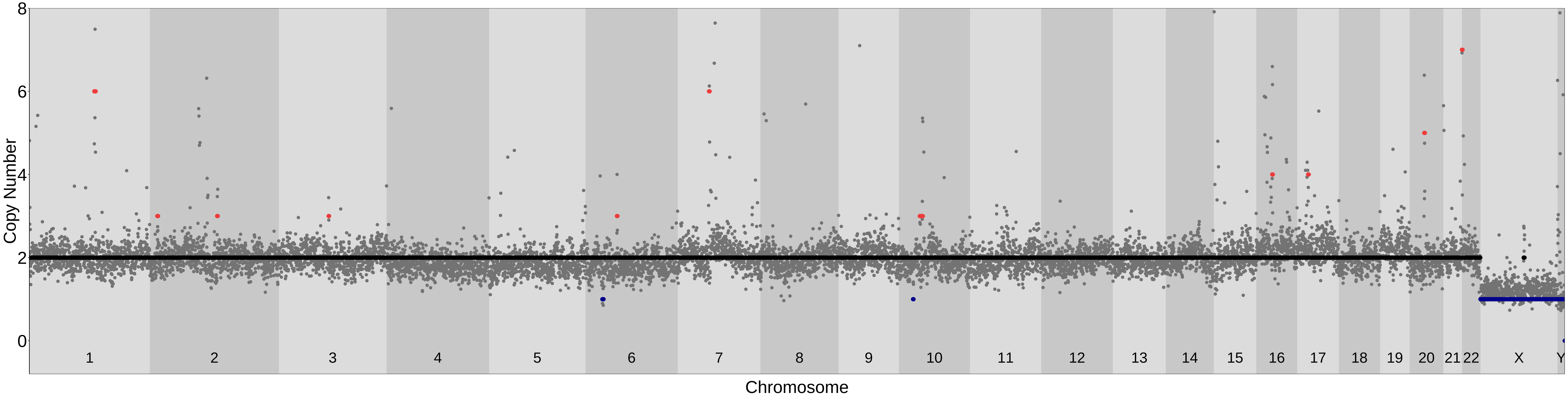

### PicoPLEX_76bp_250kb_Liftover; MAD= 0.12 Confidence_score= 0.9

| samples | ID | shared_ind_number | chr | cn | cn_median | start | end | width |
|---|---|---|---|---|---|---|---|---|
| MSA−1 | A29_v2_Exp9.1_sn10 | 4 | chr1 | 6 | 5.69 | 119788580 | 149878253 | 30089674 |
| | | 3 | chr2 | 3 | 2.74 | 129947291 | 132549472 | 2602182 |
| | | 4 | chr6 | 1 | 1.33 | 31077463 | 33277394 | 2199932 |
| | | 4 | chr9 | 14 | 13.64 | 38640746 | 68419166 | 29778421 |
| | | 2 | chr11 | 3 | 2.74 | 48131572 | 54757193 | 6625622 |
| | | 1 | chrX | 1 | 1.09 | 1 | 89017669 | 89017669 |
| | | 1 | chrX | 1 | 0.95 | 92901881 | 156040895 | 63139015 |

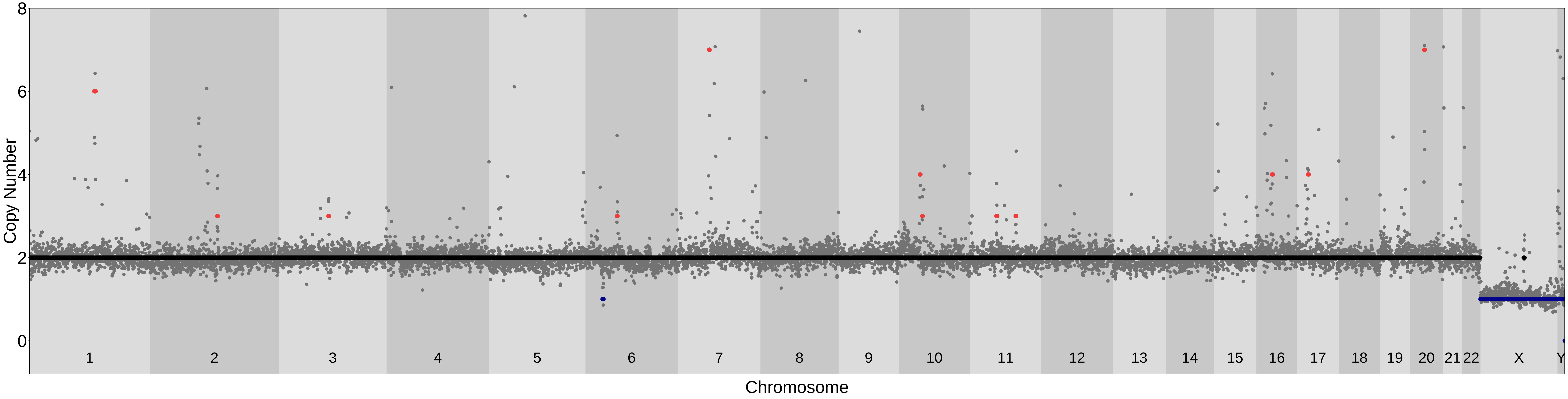

### PicoPLEX_76bp_250kb_Liftover; MAD= 0.14 Confidence_score= 0.81

| samples | ID | shared_ind_number | chr | cn | cn_median | start | end | width |
|---|---|---|---|---|---|---|---|---|
| MSA−1 | A9_v3_Exp6.1_sn1 | 4 | chr1 | 6 | 6.17 | 119788580 | 149878253 | 30089674 |
| | | 1 | chr2 | 1 | 1.34 | 202060106 | 203660707 | 1600602 |
| | | 4 | chr6 | 3 | 2.58 | 57145229 | 61809411 | 4664183 |
| | | 4 | chr9 | 12 | 12.37 | 38640746 | 68419166 | 29778421 |
| | | 1 | chr13 | 3 | 2.54 | 111463618 | 114364328 | 2900711 |
| | | 4 | chr20 | 4 | 4.12 | 25724654 | 30899236 | 5174583 |
| | | 1 | chrX | 1 | 1.16 | 1 | 89017669 | 89017669 |
| | | 1 | chrX | 1 | 1.16 | 93293918 | 156040895 | 62746978 |

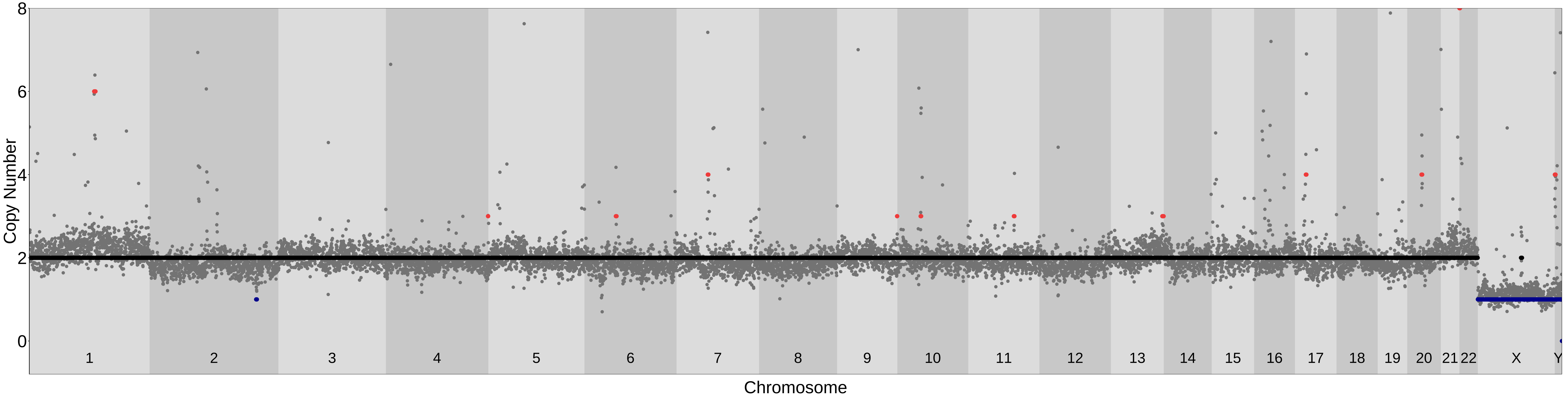

### PicoPLEX_76bp_250kb_Liftover; MAD= 0.16 Confidence_score= 0.88

| samples | ID | shared_ind_number | chr | cn | cn_median | start | end | width |
|---|---|---|---|---|---|---|---|---|
| MSA−2 | A76_Exp7_2_sn1 | 4 | chr1 | 6 | 5.87 | 119788580 | 149878253 | 30089674 |
| | | 1 | chr1 | 3 | 2.77 | 231171100 | 234837530 | 3666431 |
| | | 4 | chr2 | 3 | 3.46 | 86824353 | 96084567 | 9260215 |
| | | 3 | chr2 | 3 | 2.62 | 129947291 | 134634072 | 4686782 |
| | | 1 | chr2 | 3 | 2.50 | 153709214 | 155269401 | 1560188 |
| | | 1 | chr2 | 3 | 2.54 | 157112230 | 161103912 | 3991683 |
| | | 1 | chr3 | 3 | 2.56 | 149844365 | 158291629 | 8447265 |
| | | 1 | chr7 | 3 | 2.56 | 17154953 | 25840156 | 8685204 |
| | | 1 | chr7 | 3 | 2.62 | 27413621 | 30298607 | 2884987 |
| | | 3 | chr7 | 6 | 6.38 | 56573819 | 62787680 | 6213862 |
| | | 4 | chr9 | 12 | 12.30 | 38640746 | 68419166 | 29778421 |
| | | 1 | chr9 | 3 | 2.80 | 116968958 | 120331164 | 3362207 |
| | | 1 | chr10 | 3 | 2.62 | 36738929 | 39017432 | 2278504 |
| | | 1 | chr11 | 3 | 2.79 | 3470720 | 7393485 | 3922766 |
| | | 1 | chr11 | 3 | 2.51 | 17912077 | 25832177 | 7920101 |
| | | 1 | chr11 | 3 | 2.51 | 33722635 | 55914767 | 22192133 |
| | | 3 | chr11 | 3 | 2.87 | 88690994 | 91083415 | 2392422 |

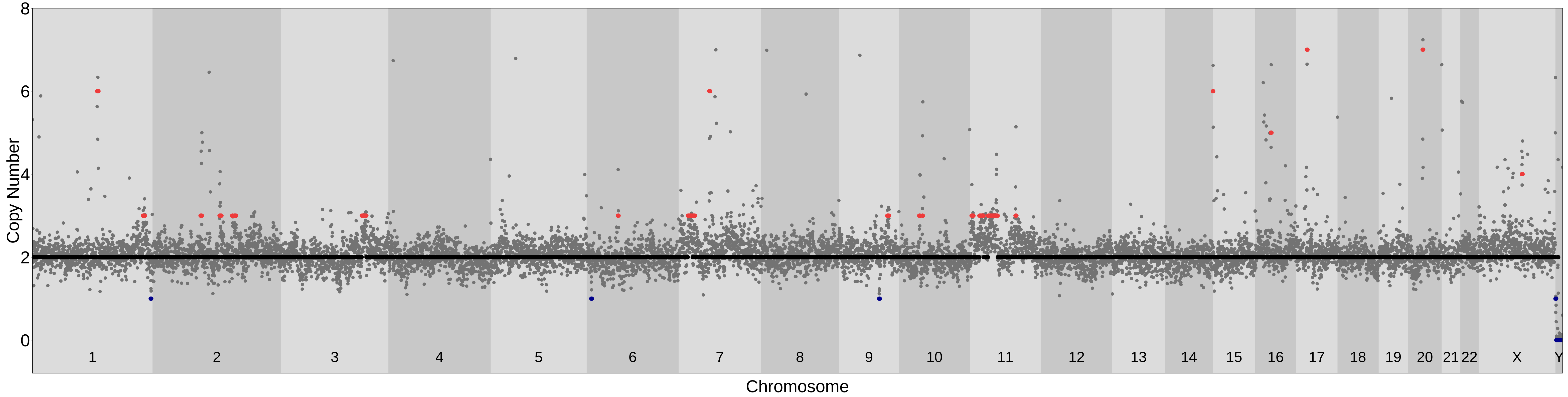

#### PicoPLEX_76bp_250kb_Liftover; MAD= 0.19 Confidence_score= 0.76

| samples | ID | shared_ind_number | chr | cn | cn_median | start | end | width |
|---|---|---|---|---|---|---|---|---|
| MSA−2 | A77_Exp7_2_sn2 | 4 | chr1 | 5 | 5.13 | 119788580 | 149878253 | 30089674 |
| | | 1 | chr3 | 1 | 1.39 | 16571697 | 18926587 | 2354891 |
| | | 1 | chr4 | 1 | 1.48 | 120794027 | 126819986 | 6025960 |
| | | 1 | chr9 | 3 | 2.53 | 9148749 | 14122032 | 4973284 |
| | | 4 | chr9 | 14 | 14.18 | 38640746 | 68419166 | 29778421 |
| | | 1 | chr11 | 1 | 1.45 | 127756934 | 129316730 | 1559797 |
| | | 4 | chr15 | 4 | 3.58 | 1 | 24585546 | 24585546 |

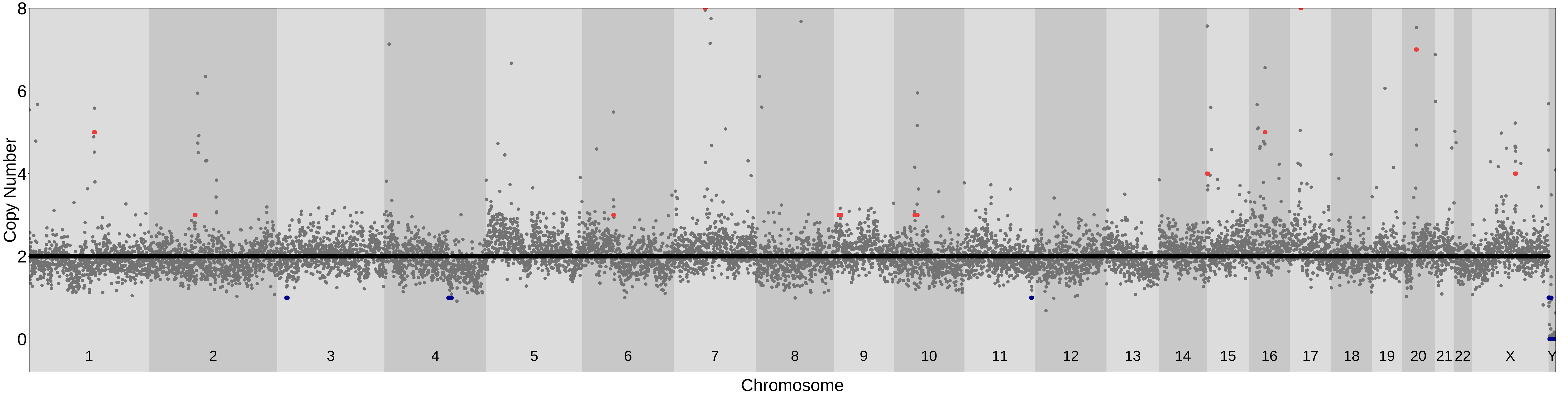

### PicoPLEX_76bp_250kb_Liftover; MAD= 0.13 Confidence_score= 0.89

| samples | ID | shared_ind_number | chr | cn | cn_median | start | end | width |
|---|---|---|---|---|---|---|---|---|
| MSA−2 | A78_Exp7_2_sn3 | 4 | chr1 | 5 | 5.43 | 119788580 | 149878253 | 30089674 |
| | | 3 | chr2 | 3 | 2.95 | 129947291 | 132549472 | 2602182 |
| | | 4 | chr9 | 12 | 11.53 | 38640746 | 68419166 | 29778421 |
| | | 2 | chr11 | 3 | 2.66 | 48131572 | 55342112 | 7210541 |

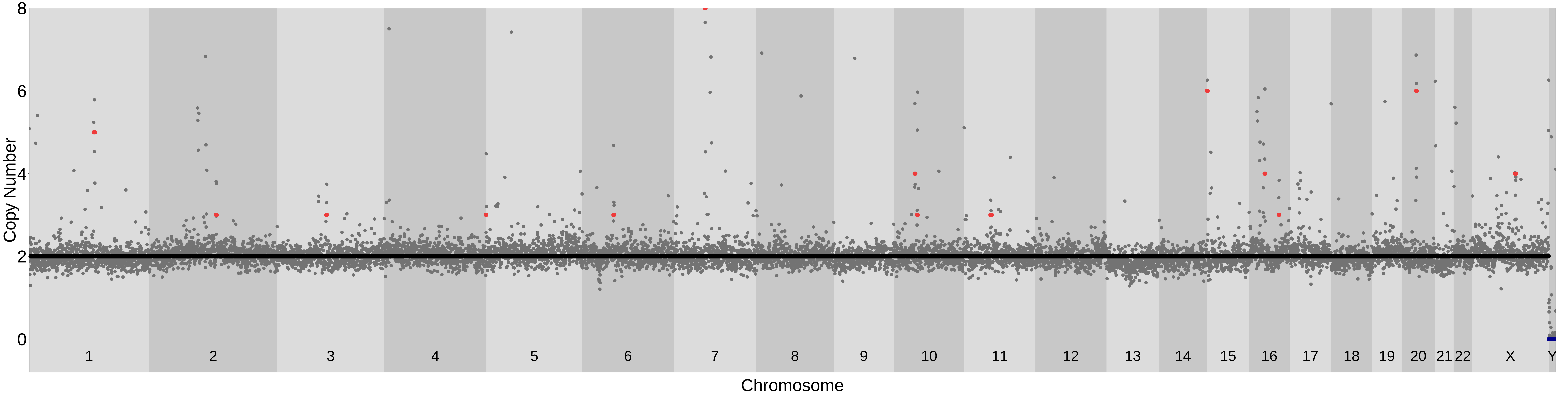

#### PicoPLEX_76bp_250kb_Liftover; MAD= 0.14 Confidence_score= 0.83

| samples | ID | shared_ind_number | chr | cn | cn_median | start | end | width |
|---|---|---|---|---|---|---|---|---|
| MSA−2 | A79_Exp7_2_sn4 | 4 | chr1 | 6 | 5.80 | 119788580 | 149878253 | 30089674 |
| | | 4 | chr2 | 3 | 3.42 | 86824353 | 96084567 | 9260215 |
| | | 3 | chr2 | 3 | 3.07 | 129947291 | 132549472 | 2602182 |
| | | 4 | chr9 | 12 | 11.70 | 38640746 | 68419166 | 29778421 |

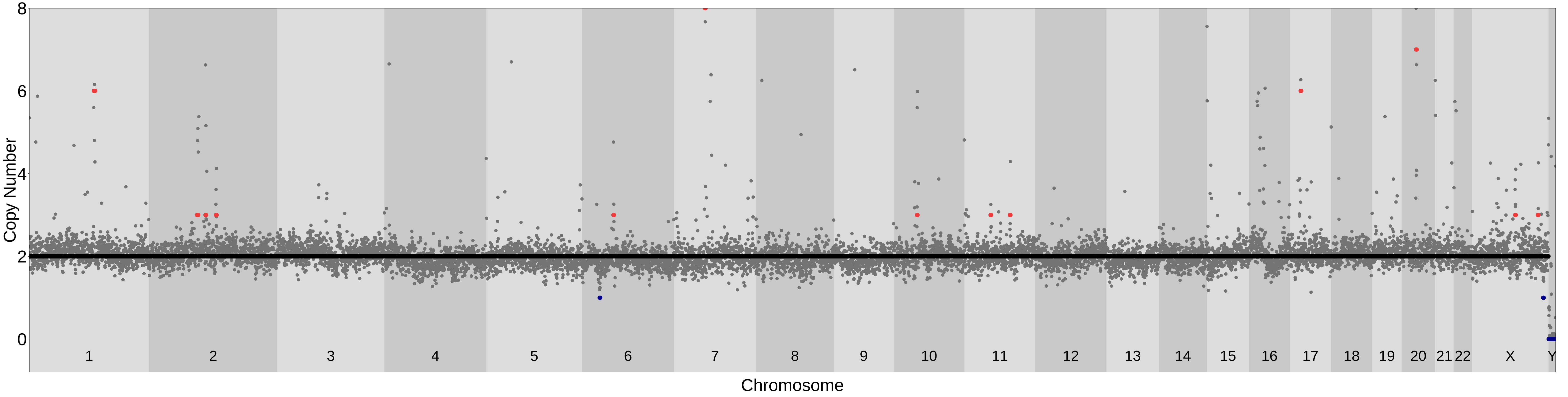

### PicoPLEX_76bp_250kb_Liftover; MAD= 0.15 Confidence_score= 0.88

| samples | ID | shared_ind_number | chr | cn | cn_median | start | end | width |
|---|---|---|---|---|---|---|---|---|
| MSA−2 | A80_Exp7_2_sn5 | 4 | chr1 | 5 | 5.24 | 119788580 | 149878253 | 30089674 |
| | | 3 | chr2 | 3 | 3.04 | 129947291 | 132805370 | 2858080 |
| | | 4 | chr9 | 12 | 12.49 | 38640746 | 68419166 | 29778421 |
| | | 2 | chr10 | 3 | 3.11 | 37872529 | 42484908 | 4612380 |
| | | 1 | chr10 | 1 | 1.41 | 113864762 | 116722933 | 2858172 |
| | | 2 | chr11 | 3 | 3.30 | 48440666 | 54757193 | 6316528 |
| | | 1 | chr20 | 3 | 2.53 | 41866980 | 43422771 | 1555792 |

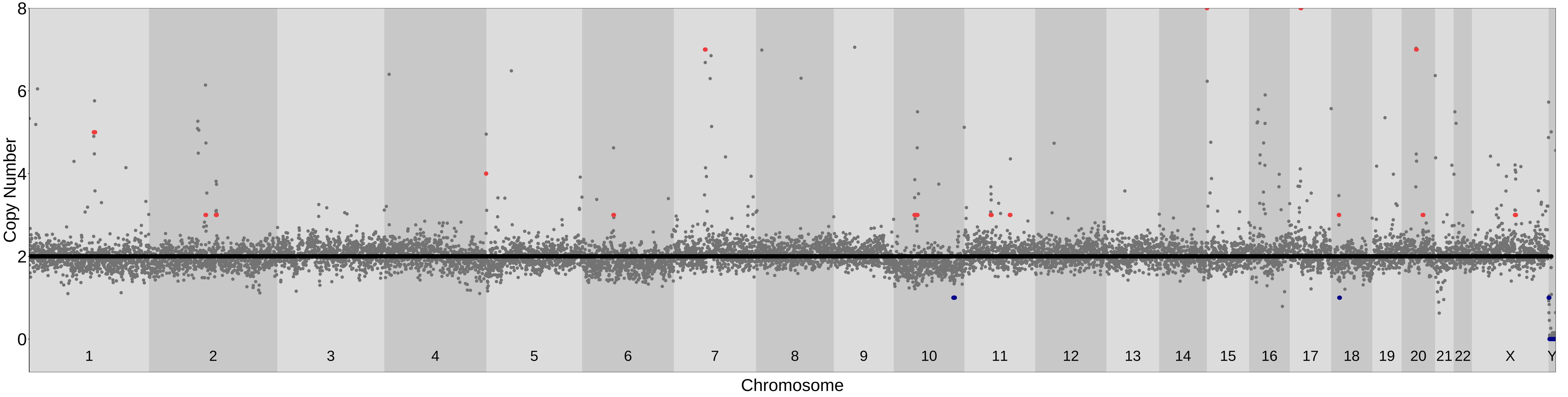

### PicoPLEX_76bp_250kb_Liftover; MAD= 0.15 Confidence_score= 0.78

| samples | ID | shared_ind_number | chr | cn | cn_median | start | end | width |
|---|---|---|---|---|---|---|---|---|
| MSA−2 | A81_Exp7_2_sn6 | 4 | chr1 | 5 | 5.25 | 119788580 | 149878253 | 30089674 |
| | | 3 | chr2 | 3 | 2.92 | 129947291 | 132549472 | 2602182 |
| | | 1 | chr4 | 1 | 1.40 | 87739794 | 91436980 | 3697187 |
| | | 1 | chr4 | 1 | 1.46 | 123426317 | 135686477 | 12260161 |
| | | 1 | chr5 | 3 | 2.52 | 142468649 | 145614906 | 3146258 |
| | | 4 | chr6 | 3 | 3.27 | 57145229 | 61809411 | 4664183 |
| | | 2 | chr7 | 3 | 2.51 | 55636895 | 57799139 | 2162245 |
| | | 4 | chr9 | 12 | 12.23 | 38640746 | 68419166 | 29778421 |
| | | 1 | chr10 | 1 | 1.29 | 1408677 | 4752292 | 3343616 |
| | | 1 | chr11 | 3 | 2.65 | 48131572 | 50247851 | 2116280 |
| | | 1 | chr20 | 1 | 1.48 | 39518785 | 41599795 | 2081011 |
| | | 1 | chr20 | 1 | 1.49 | 59635662 | 61172580 | 1536919 |

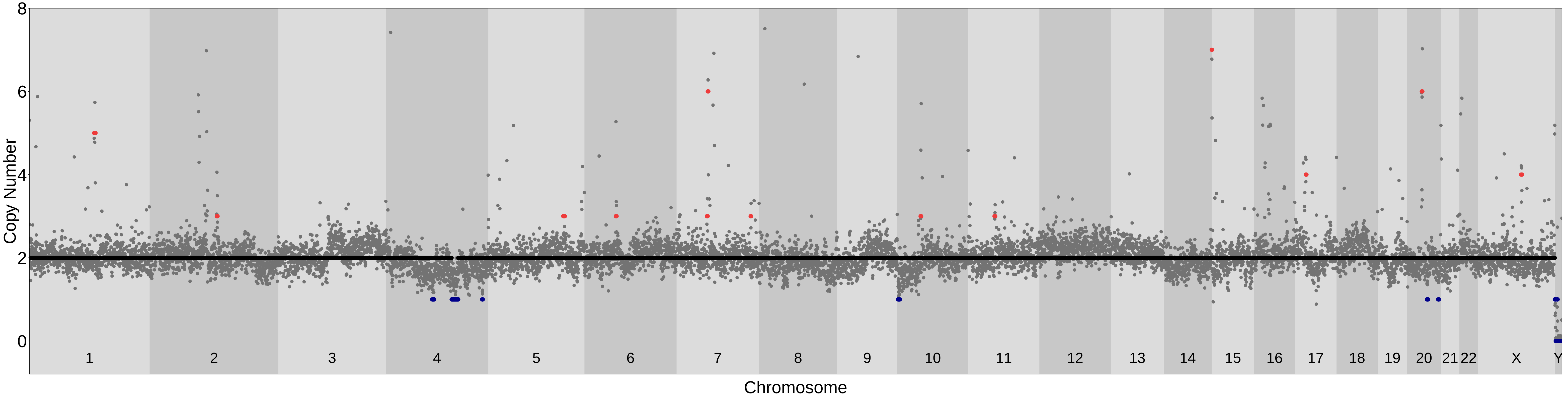

### PicoPLEX_76bp_250kb_Liftover; MAD= 0.15 Confidence_score= 0.78

| samples | ID | shared_ind_number | chr | cn | cn_median | start | end | width |
|---|---|---|---|---|---|---|---|---|
| MSA−2 | A82_Exp7_2_sn7 | 1 | chr1 | 3 | 2.52 | 11727877 | 15341038 | 3613162 |
| | | 4 | chr1 | 6 | 5.74 | 119788580 | 149878253 | 30089674 |
| | | 1 | chr1 | 3 | 2.60 | 180311568 | 185548051 | 5236484 |
| | | 3 | chr2 | 3 | 2.84 | 129947291 | 132549472 | 2602182 |
| | | 1 | chr3 | 3 | 2.89 | 1 | 30817485 | 30817485 |
| | | 1 | chr3 | 3 | 2.67 | 53298781 | 55410569 | 2111789 |
| | | 1 | chr3 | 3 | 2.57 | 74463667 | 77282095 | 2818429 |
| | | 2 | chr3 | 3 | 2.67 | 128569151 | 131341052 | 2771902 |
| | | 1 | chr3 | 3 | 2.64 | 142794700 | 150358564 | 7563865 |
| | | 1 | chr7 | 1 | 1.48 | 38189519 | 39758093 | 1568575 |
| | | 3 | chr7 | 5 | 5.30 | 56573819 | 62787680 | 6213862 |
| | | 1 | chr7 | 1 | 1.42 | 113885301 | 128232704 | 14347404 |
| | | 4 | chr9 | 14 | 13.54 | 38640746 | 68419166 | 29778421 |
| | | 2 | chr11 | 3 | 2.84 | 48440666 | 54757193 | 6316528 |
| | | 1 | chr11 | 1 | 1.46 | 123591210 | 129586379 | 5995170 |
| | | 1 | chr14 | 3 | 2.87 | 46045456 | 49500228 | 3454773 |
| | | 1 | chr14 | 3 | 2.56 | 77200397 | 82421609 | 5221213 |
| | | 1 | chr17 | 3 | 2.65 | 64679165 | 77917904 | 13238740 |
| | | 1 | chr20 | 3 | 2.79 | 5868836 | 8723014 | 2854179 |

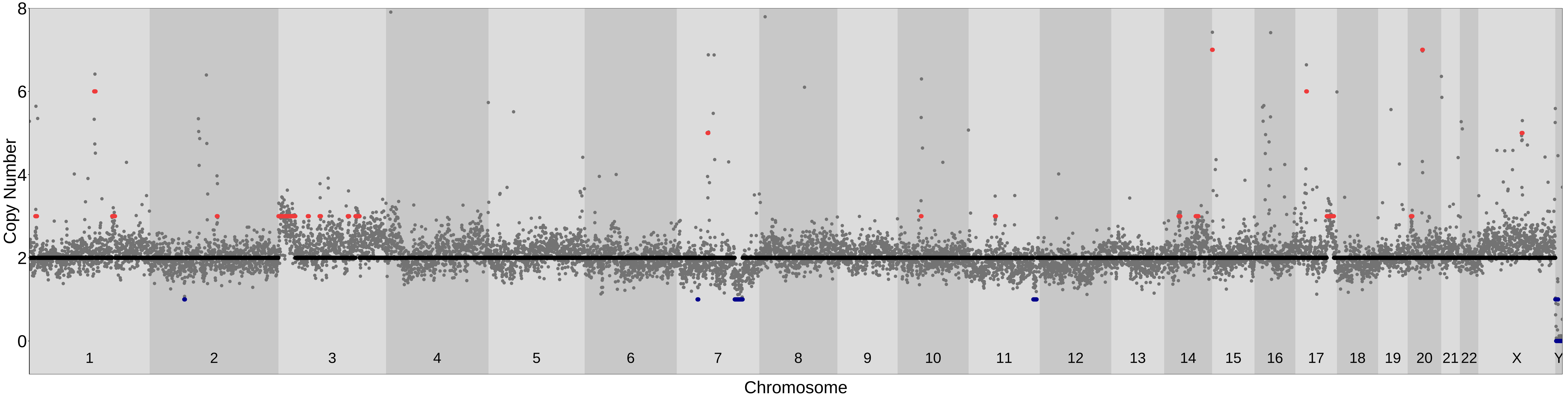

### PicoPLEX_76bp_250kb_Liftover; MAD= 0.13 Confidence_score= 0.86

| samples | ID | shared_ind_number | chr | cn | cn_median | start | end | width |
|---|---|---|---|---|---|---|---|---|
| MSA−2 | A83_Exp7_2_sn9 | 4 | chr1 | 6 | 5.77 | 119788580 | 149878253 | 30089674 |
| | | 3 | chr2 | 3 | 2.68 | 129947291 | 132549472 | 2602182 |
| | | 2 | chr7 | 3 | 2.56 | 55636895 | 57799139 | 2162245 |
| | | 4 | chr9 | 12 | 12.07 | 38640746 | 68419166 | 29778421 |
| | | 2 | chr11 | 3 | 2.92 | 48440666 | 54757193 | 6316528 |

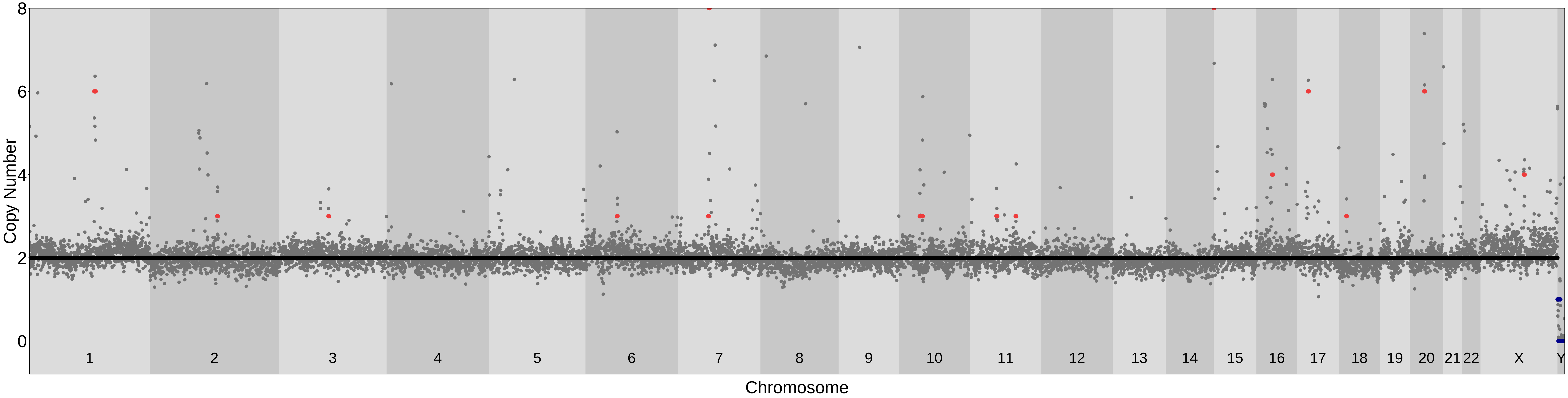

### PicoPLEX_101bp_250kb_Liftover; MAD= 0.2 Confidence_score= 0.78

| samples | ID | shared_ind_number | chr | cn | cn_median | start | end | width |
|---|---|---|---|---|---|---|---|---|
| Control | L11_Exp1.8.1B_sn3 | 3 | chr1 | 7 | 7.14 | 119872097 | 147088134 | 27216038 |
| | | 4 | chr2 | 3 | 2.58 | 86637708 | 97657618 | 11019911 |
| | | 3 | chr2 | 3 | 2.83 | 129864209 | 132266362 | 2402154 |
| | | 1 | chr4 | 3 | 3.01 | 33233263 | 35268589 | 2035327 |
| | | 1 | chr4 | 3 | 2.63 | 118298745 | 138009017 | 19710273 |
| | | 1 | chr4 | 3 | 2.98 | 180678819 | 182472287 | 1793469 |
| | | 1 | chr7 | 3 | 2.51 | 8796991 | 25566626 | 16769636 |
| | | 3 | chr7 | 4 | 4.05 | 57544879 | 64022564 | 6477686 |
| | | 4 | chr9 | 12 | 12.34 | 39017121 | 68523575 | 29506455 |
| | | 2 | chr9 | 3 | 2.57 | 68523576 | 88753990 | 20230415 |
| | | 1 | chr9 | 3 | 2.55 | 98977391 | 104662269 | 5684879 |
| | | 1 | chr9 | 3 | 2.72 | 112363522 | 120826819 | 8463298 |
| | | 1 | chr13 | 1 | 1.09 | 1 | 73082997 | 73082997 |
| | | 1 | chr13 | 1 | 0.90 | 75391006 | 114364328 | 38973323 |
| | | 1 | chr14 | 3 | 2.55 | 84972577 | 88070044 | 3097468 |
| | | 4 | chr20 | 3 | 3.23 | 25752818 | 30908344 | 5155527 |
| | | 4 | chr21 | 8 | 7.62 | 1 | 13983405 | 13983405 |

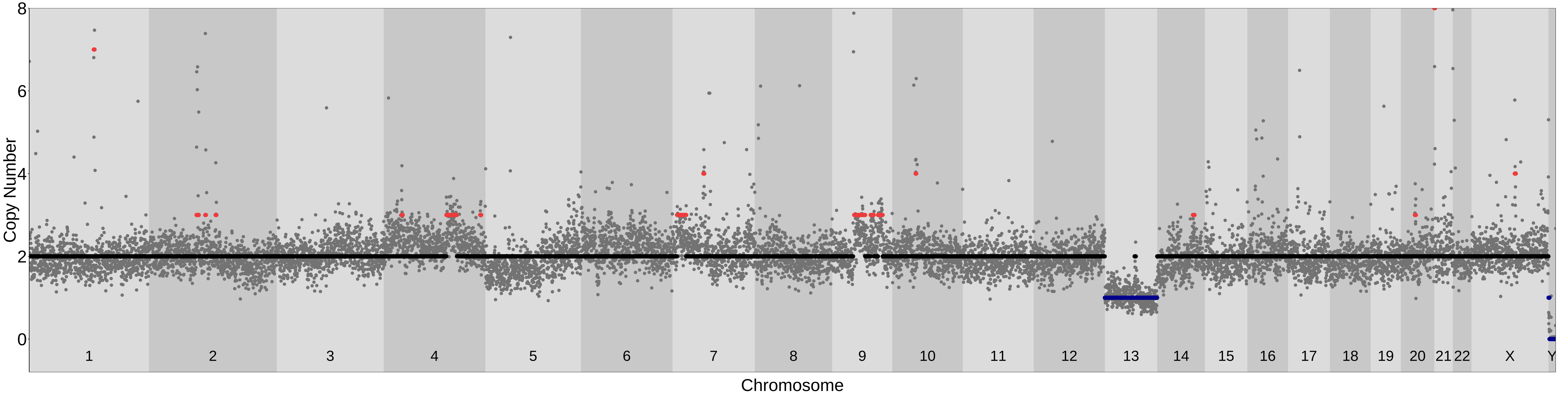

### PicoPLEX_101bp_250kb_Liftover; MAD= 0.17 Confidence_score= 0.87

| samples | ID | shared_ind_number | chr | cn | cn_median | start | end | width |
|---|---|---|---|---|---|---|---|---|
| Control | L12_Exp1.8.1B_sn4 | 4 | chr1 | 5 | 5.09 | 119872097 | 149971236 | 30099140 |
| | | 4 | chr2 | 5 | 4.51 | 86637708 | 94762314 | 8124607 |
| | | 3 | chr2 | 3 | 2.79 | 129864209 | 132563696 | 2699488 |
| | | 3 | chr6 | 1 | 1.42 | 29652898 | 33594914 | 3942017 |
| | | 3 | chr7 | 3 | 3.42 | 56742046 | 64022564 | 7280519 |
| | | 4 | chr9 | 12 | 11.89 | 39017121 | 68523575 | 29506455 |
| | | 4 | chr20 | 4 | 3.54 | 25752818 | 30908344 | 5155527 |
| | | 4 | chr21 | 9 | 8.72 | 1 | 13983405 | 13983405 |

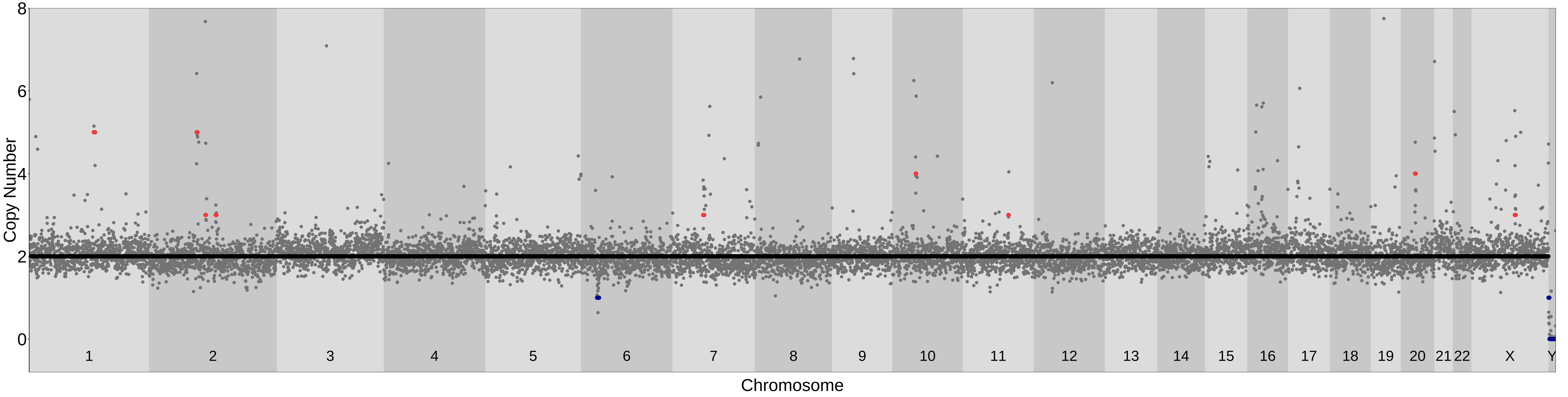

### PicoPLEX_101bp_250kb_Liftover; MAD= 0.17 Confidence_score= 0.75

| samples | ID | shared_ind_number | chr | cn | cn_median | start | end | width |
|---|---|---|---|---|---|---|---|---|
| Control | L8_Exp1.8.1A_sn1 | 4 | chr1 | 5 | 4.92 | 119872097 | 149971236 | 30099140 |
| | | 2 | chr2 | 1 | 1.36 | 1 | 1576659 | 1576659 |
| | | 1 | chr2 | 1 | 1.46 | 6719800 | 8497841 | 1778042 |
| | | 4 | chr2 | 5 | 5.17 | 86637708 | 94762314 | 8124607 |
| | | 4 | chr2 | 3 | 2.50 | 109674602 | 113611097 | 3936496 |
| | | 3 | chr6 | 1 | 1.48 | 28610557 | 33594914 | 4984358 |
| | | 4 | chr9 | 13 | 12.92 | 39017121 | 68523575 | 29506455 |
| | | 1 | chr9 | 3 | 2.57 | 77260975 | 84048984 | 6788010 |
| | | 1 | chr19 | 1 | 1.48 | 51508278 | 55218025 | 3709748 |
| | | 4 | chr20 | 4 | 4.12 | 25752818 | 30908344 | 5155527 |

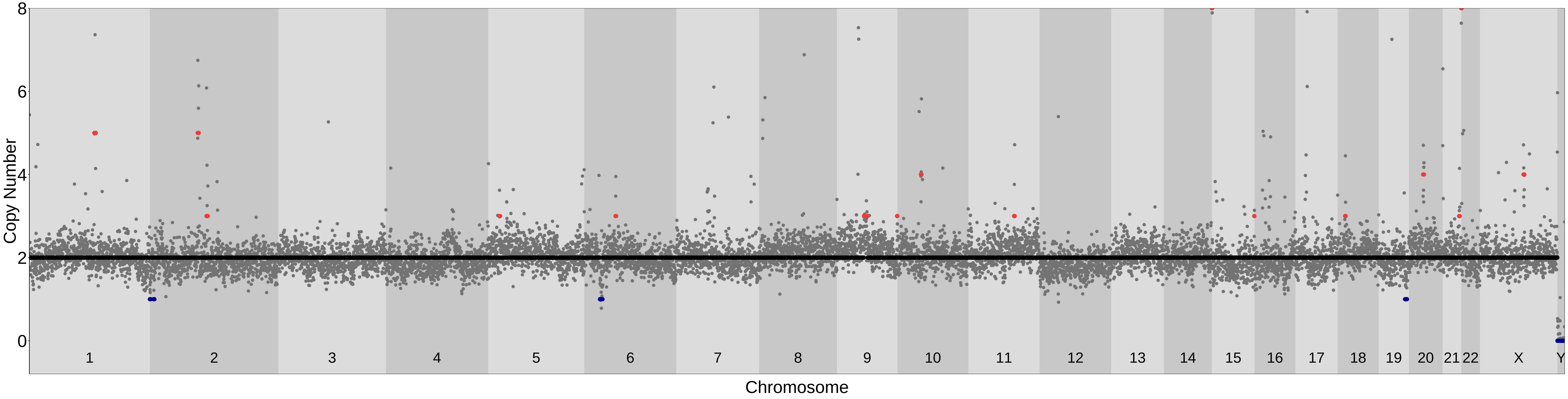

### PicoPLEX_101bp_250kb_Liftover; MAD= 0.18 Confidence_score= 0.78

| samples | ID | shared_ind_number | chr | cn | cn_median | start | end | width |
|---|---|---|---|---|---|---|---|---|
| Control | L9_Exp1.8.1A_sn3 | 3 | chr1 | 6 | 6.48 | 119872097 | 147088134 | 27216038 |
| | | 4 | chr2 | 5 | 5.31 | 86637708 | 94762314 | 8124607 |
| | | 3 | chr2 | 3 | 2.74 | 129864209 | 132563696 | 2699488 |
| | | 3 | chr6 | 1 | 1.36 | 28610557 | 32827875 | 4217319 |
| | | 4 | chr9 | 12 | 12.45 | 39017121 | 68523575 | 29506455 |
| | | 2 | chr11 | 3 | 2.53 | 27954308 | 30305539 | 2351232 |
| | | 4 | chr17 | 3 | 3.05 | 21341319 | 27065465 | 5724147 |
| | | 1 | chr18 | 1 | 1.48 | 63101084 | 72063267 | 8962184 |
| | | 4 | chr20 | 4 | 4.40 | 25752818 | 30908344 | 5155527 |
| | | 4 | chr21 | 9 | 8.54 | 1 | 13983405 | 13983405 |
| | | 3 | chr22 | 3 | 2.83 | 1 | 19092117 | 19092117 |

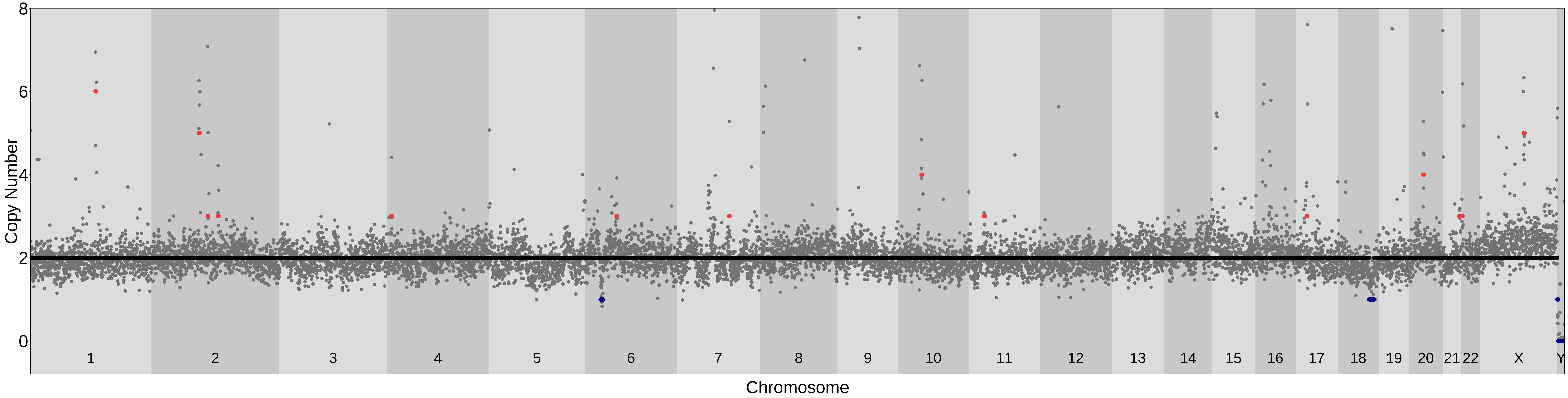

### PicoPLEX_101bp_250kb_Liftover; MAD= 0.17 Confidence_score= 0.72

| samples | ID | shared_ind_number | chr | cn | cn_median | start | end | width |
|---|---|---|---|---|---|---|---|---|
| Fibr | L2_4min_Exp1.7A_sn2 | 3 | chr1 | 9 | 8.72 | 119872097 | 147088134 | 27216038 |
| | | 1 | chr2 | 3 | 2.64 | 75035148 | 77920033 | 2884886 |
| | | 4 | chr2 | 3 | 2.74 | 86637708 | 97657618 | 11019911 |
| | | 1 | chr4 | 5 | 4.54 | 88454582 | 90255973 | 1801392 |
| | | 4 | chr6 | 1 | 1.14 | 30948846 | 32827875 | 1879030 |
| | | 1 | chr9 | 1 | 1.36 | 16951353 | 39017120 | 22065768 |
| | | 4 | chr9 | 8 | 8.02 | 39017121 | 68523575 | 29506455 |
| | | 1 | chr9 | 1 | 1.39 | 68523576 | 77260974 | 8737399 |
| | | 1 | chr12 | 1 | 1.42 | 20208904 | 23608940 | 3400037 |
| | | 1 | chr14 | 3 | 2.51 | 26121613 | 28482193 | 2360581 |
| | | 4 | chr20 | 4 | 3.63 | 25752818 | 30908344 | 5155527 |

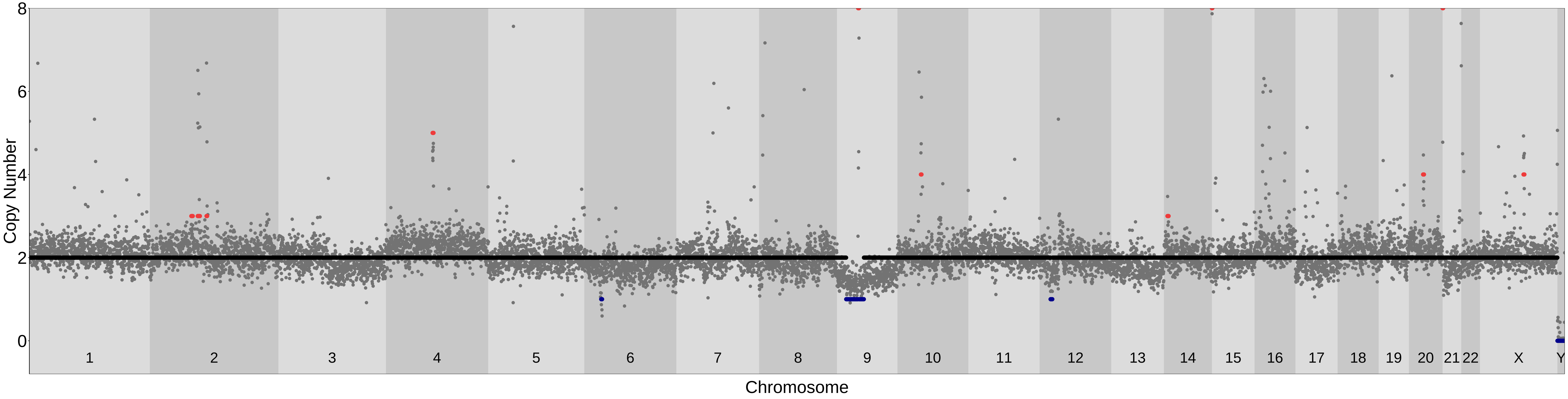

### PicoPLEX_101bp_250kb_Liftover; MAD= 0.16 Confidence_score= 0.71

| samples | ID | shared_ind_number | chr | cn | cn_median | start | end | width |
|---|---|---|---|---|---|---|---|---|
| Fibr | L5_Exp1.7A_sn1 | 4 | chr1 | 6 | 5.64 | 119872097 | 149971236 | 30099140 |
| | | 2 | chr2 | 1 | 1.41 | 1 | 3894901 | 3894901 |
| | | 4 | chr2 | 5 | 5.04 | 86637708 | 94762314 | 8124607 |
| | | 4 | chr2 | 3 | 2.70 | 109674602 | 113907226 | 4232625 |
| | | 1 | chr3 | 1 | 1.23 | 161792006 | 163340109 | 1548104 |
| | | 1 | chr4 | 3 | 2.60 | 8145024 | 17847217 | 9702194 |
| | | 1 | chr4 | 4 | 3.74 | 88454582 | 90255973 | 1801392 |
| | | 3 | chr6 | 1 | 1.43 | 28874539 | 33594914 | 4720376 |
| | | 4 | chr6 | 3 | 2.97 | 56725150 | 61039414 | 4314265 |
| | | 1 | chr7 | 1 | 1.39 | 134960093 | 139582530 | 4622438 |
| | | 1 | chr7 | 1 | 1.39 | 141129694 | 143192129 | 2062436 |
| | | 1 | chr8 | 1 | 1.49 | 41603079 | 43676579 | 2073501 |
| | | 4 | chr9 | 10 | 10.18 | 39017121 | 68523575 | 29506455 |
| | | 1 | chr10 | 3 | 2.70 | 5999132 | 8826810 | 2827679 |
| | | 4 | chr10 | 3 | 2.57 | 45594822 | 50291880 | 4697059 |
| | | 4 | chr17 | 3 | 3.05 | 21341319 | 27065465 | 5724147 |
| | | 4 | chr20 | 3 | 3.45 | 25752818 | 30908344 | 5155527 |

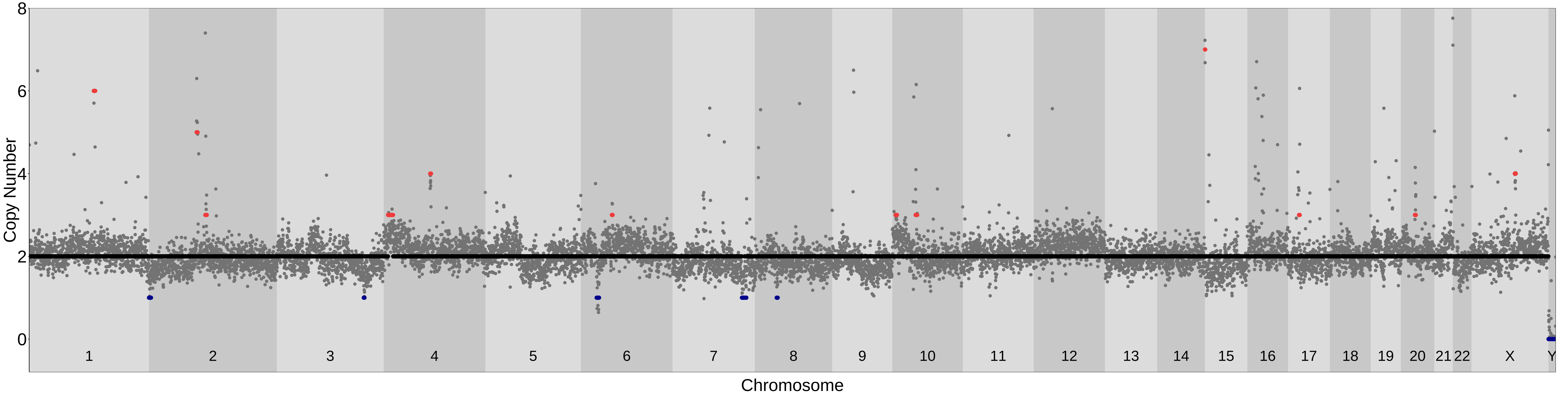

### PTA_101bp_500kb_Liftover; MAD= 0.23 Confidence_score= 0.83

| samples | ID | shared_ind_number | chr | cn | cn_median | start | end | width |
|---|---|---|---|---|---|---|---|---|
| MSA−1 | A4_PTA_Exp9.2_sn_2 | 1 | chr1 | 3 | 2.74 | 1 | 18496462 | 18496462 |
| | | 3 | chr1 | 6 | 5.61 | 119614360 | 150230984 | 30616625 |
| | | 2 | chr2 | 3 | 2.95 | 86384595 | 97657618 | 11273024 |
| | | 1 | chr4 | 3 | 2.59 | 3146119 | 17166542 | 14020424 |
| | | 1 | chr6 | 1 | 1.39 | 23738219 | 33335086 | 9596868 |
| | | 1 | chr7 | 3 | 2.57 | 31385982 | 37849499 | 6463518 |
| | | 3 | chr9 | 7 | 6.62 | 37235975 | 68523573 | 31287599 |
| | | 2 | chr9 | 3 | 3.09 | 133563179 | 138394717 | 4831539 |
| | | 2 | chr11 | 3 | 2.84 | 1 | 5143011 | 5143011 |
| | | 1 | chr16 | 3 | 2.55 | 1609450 | 29857565 | 28248116 |
| | | 1 | chrX | 1 | 1.20 | 1 | 156040895 | 156040895 |

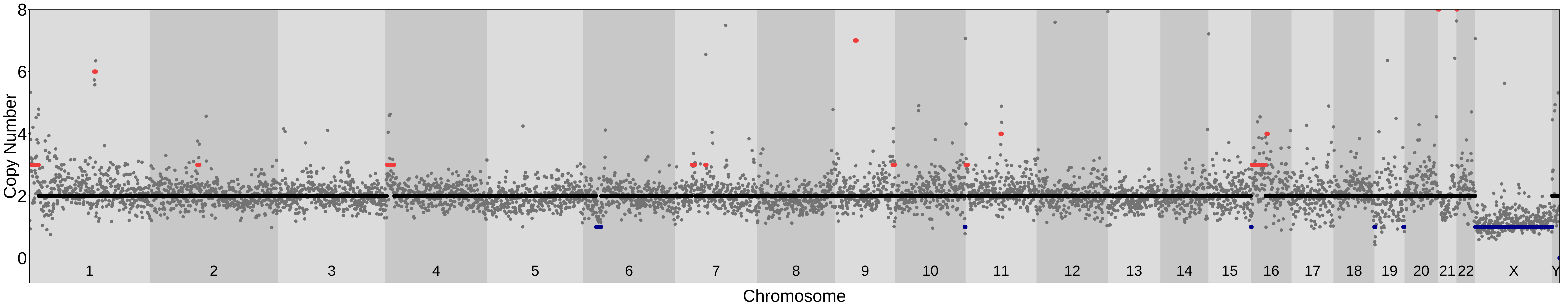

### PTA_101bp_500kb_Liftover; MAD= 0.19 Confidence_score= 0.75

| samples | ID | shared_ind_number | chr | cn | cn_median | start | end | width |
|---|---|---|---|---|---|---|---|---|
| MSA−1 | A5_PTA_Exp9.2_sn9 | 3 | chr1 | 5 | 5.47 | 119614360 | 150230984 | 30616625 |
| | | 1 | chr3 | 1 | 1.45 | 31506731 | 35630883 | 4124153 |
| | | 2 | chr3 | 1 | 1.44 | 46480569 | 49561276 | 3080708 |
| | | 1 | chr4 | 3 | 2.68 | 3146119 | 13003037 | 9856919 |
| | | 1 | chr5 | 3 | 2.56 | 143757999 | 149455361 | 5697363 |
| | | 2 | chr6 | 1 | 1.40 | 30174593 | 33335086 | 3160494 |
| | | 1 | chr13 | 1 | 1.49 | 80184213 | 85819272 | 5635060 |
| | | 1 | chr17 | 3 | 2.72 | 51160513 | 57819122 | 6658610 |
| | | 1 | chr18 | 1 | 1.25 | 21079922 | 24159709 | 3079788 |
| | | 2 | chr18 | 1 | 1.43 | 73059886 | 80373285 | 7313400 |
| | | 1 | chr21 | 1 | 1.33 | 18717138 | 26935752 | 8218615 |
| | | 1 | chrX | 1 | 1.25 | 1 | 88791987 | 88791987 |
| | | 1 | chrX | 1 | 1.07 | 93394653 | 156040895 | 62646243 |

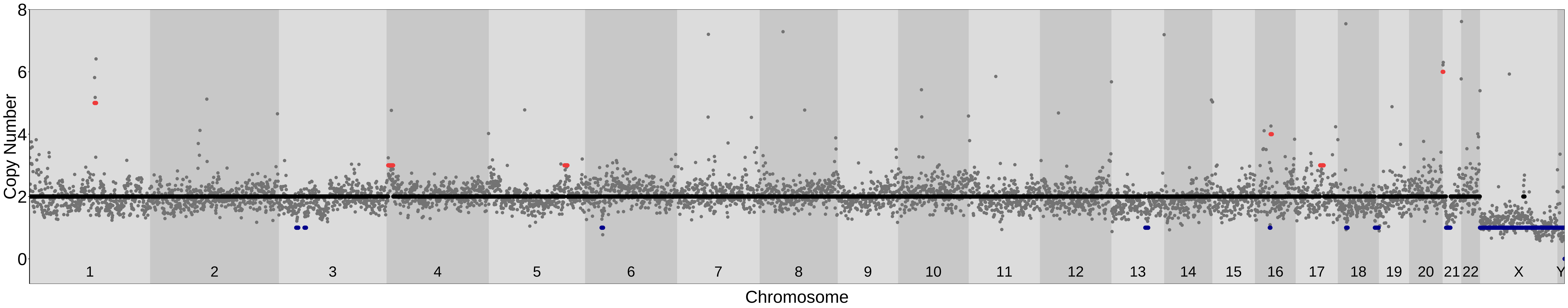

### PTA_101bp_500kb_Liftover; MAD= 0.18 Confidence_score= 0.88

| samples | ID | shared_ind_number | chr | cn | cn_median | start | end | width |
|---|---|---|---|---|---|---|---|---|
| MSA−1 | A6_PTA_Exp9.2_sn10 | 3 | chr1 | 5 | 5.02 | 119614360 | 150230984 | 30616625 |
| | | 1 | chr1 | 3 | 2.51 | 202731123 | 206522437 | 3791315 |
| | | 2 | chr2 | 3 | 3.24 | 86384595 | 97657618 | 11273024 |
| | | 2 | chr6 | 1 | 1.44 | 28610557 | 33335086 | 4724530 |
| | | 1 | chr12 | 3 | 2.71 | 58089213 | 72125291 | 14036079 |
| | | 1 | chr12 | 3 | 2.69 | 82975877 | 108705299 | 25729423 |
| | | 2 | chr12 | 3 | 3.01 | 124992315 | 131699400 | 6707086 |
| | | 1 | chr16 | 3 | 2.74 | 46937887 | 56197936 | 9260050 |
| | | 1 | chr19 | 1 | 1.20 | 1 | 3867248 | 3867248 |
| | | 2 | chr20 | 3 | 2.67 | 22207497 | 31460957 | 9253461 |
| | | 1 | chrX | 1 | 1.16 | 30378209 | 47603026 | 17224818 |
| | | 1 | chrX | 1 | 1.36 | 69776382 | 89468166 | 19691785 |
| | | 1 | chrX | 1 | 1.14 | 93394653 | 156040895 | 62646243 |

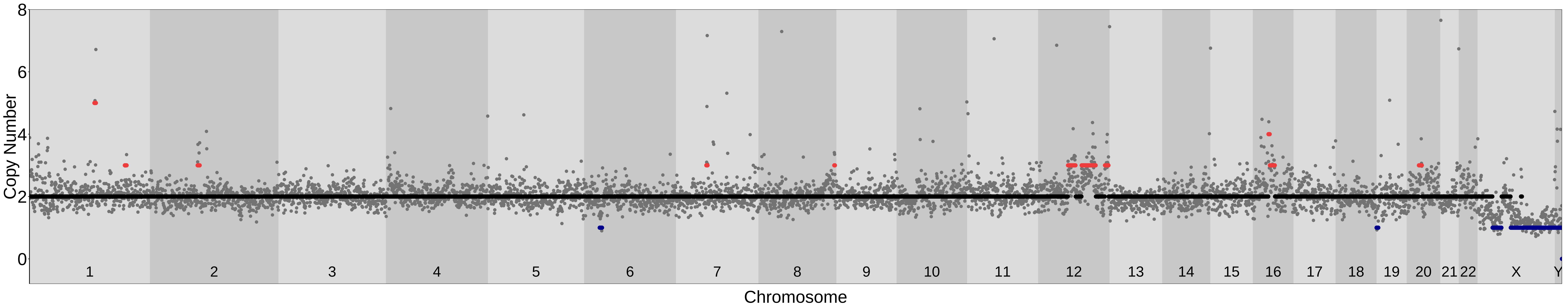

### PTA_101bp_500kb_Liftover; MAD= 0.28 Confidence_score= 0.9

| samples | ID | shared_ind_number | chr | cn | cn_median | start | end | width |
|---|---|---|---|---|---|---|---|---|
| NA12878 | A17_PTA_Test_PTA2__sn3_NA12878_S17_R | 1 | chr1 | 1 | 1.49 | 114409561 | 119614359 | 5204799 |
| | | 3 | chr1 | 5 | 5.38 | 119614360 | 150230984 | 30616625 |
| | | 2 | chr3 | 1 | 1.23 | 47001194 | 50073941 | 3072748 |
| | | 1 | chr3 | 1 | 1.35 | 155786628 | 158900220 | 3113593 |
| | | 1 | chr5 | 3 | 3.25 | 1 | 14477391 | 14477391 |
| | | 1 | chr9 | 1 | 1.22 | 99741651 | 104407794 | 4666144 |
| | | 2 | chr11 | 3 | 3.26 | 1768941 | 22853992 | 21085052 |
| | | 1 | chr15 | 1 | 1.29 | 92623323 | 96699959 | 4076637 |
| | | 1 | chr16 | 1 | 1.41 | 71538201 | 76244271 | 4706071 |
| | | 1 | chr16 | 3 | 2.60 | 85910770 | 90338345 | 4427576 |
| | | 1 | chr18 | 1 | 1.34 | 31853792 | 34954552 | 3100761 |
| | | 1 | chr18 | 1 | 1.31 | 52499763 | 58158591 | 5658829 |
| | | 1 | chr21 | 1 | 1.45 | 27447734 | 31107244 | 3659511 |

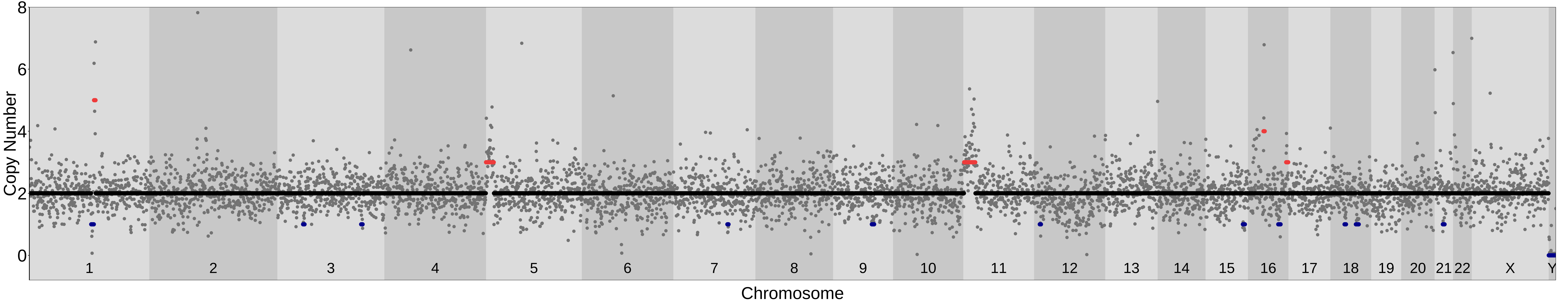

### PTA_101bp_500kb_Liftover; MAD= 0.18 Confidence_score= 0.92

| samples | ID | shared_ind_number | chr | cn | cn_median | start | end | width |
|---|---|---|---|---|---|---|---|---|
| NA12878 | A20_PTA_Test__PTA2__sn7_NA12878_S20_R | 1 | chr1 | 1 | 1.43 | 61739132 | 64810523 | 3071392 |
| | | 1 | chr1 | 1 | 1.46 | 91362207 | 94468384 | 3106178 |
| | | 3 | chr1 | 5 | 4.97 | 119614360 | 150230984 | 30616625 |
| | | 1 | chr2 | 1 | 1.36 | 42098252 | 47707496 | 5609245 |
| | | 1 | chr5 | 1 | 1.28 | 34625011 | 37738328 | 3113318 |
| | | 1 | chr6 | 1 | 1.27 | 126829227 | 135582087 | 8752861 |
| | | 1 | chr8 | 1 | 1.45 | 91086597 | 100932912 | 9846316 |
| | | 1 | chr9 | 1 | 1.36 | 123387616 | 136618464 | 13230849 |
| | | 1 | chr11 | 3 | 2.66 | 26013212 | 29670212 | 3657001 |
| | | 2 | chr11 | 3 | 2.70 | 46771101 | 61319458 | 14548358 |

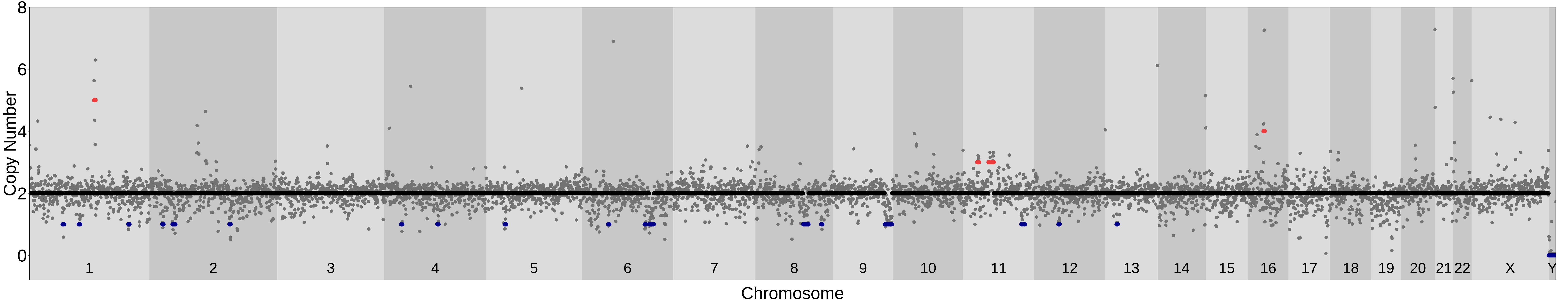

### PTA_101bp_500kb_Liftover; MAD= 0.16 Confidence_score= 0.8

| samples | ID | shared_ind_number | chr | cn | cn_median | start | end | width |
|---|---|---|---|---|---|---|---|---|
| Control | A16_PTA_Exp8.1.sn7_ | 3 | chr1 | 6 | 5.55 | 119614360 | 150230984 | 30616625 |
| | | 1 | chr1 | 3 | 2.66 | 170102062 | 173747866 | 3645805 |
| | | 2 | chr2 | 3 | 2.76 | 86384595 | 109674601 | 23290007 |
| | | 1 | chr2 | 3 | 2.59 | 113611098 | 157248033 | 43636936 |
| | | 1 | chr2 | 3 | 2.68 | 176836137 | 182508059 | 5671923 |
| | | 1 | chr2 | 3 | 2.68 | 186134892 | 196442558 | 10307667 |
| | | 2 | chr10 | 3 | 2.53 | 39332371 | 48303554 | 8971184 |
| | | 2 | chr20 | 3 | 2.69 | 18584152 | 31460957 | 12876806 |

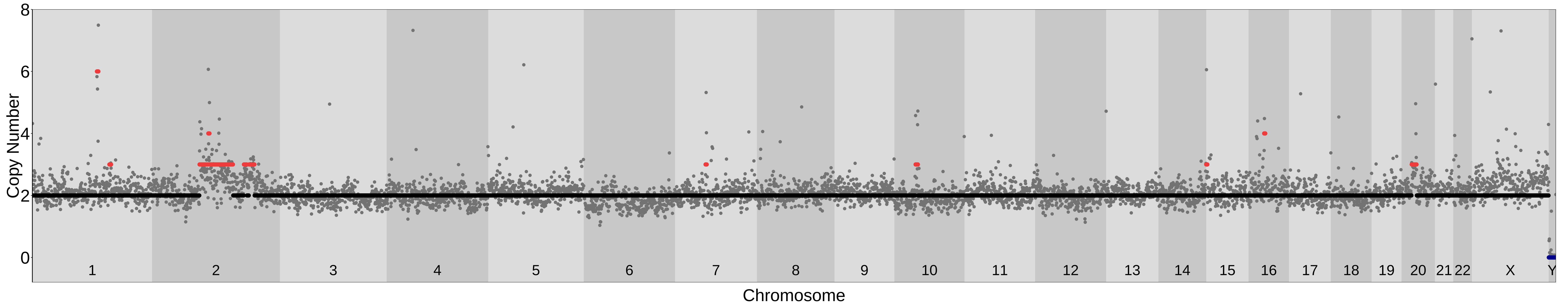

### PTA_101bp_500kb_Liftover; MAD= 0.2 Confidence_score= 0.92

| samples | ID | shared_ind_number | chr | cn | cn_median | start | end | width |
|---|---|---|---|---|---|---|---|---|
| Control | A1_PTA_Exp8.2_sn18 | 3 | chr1 | 5 | 5.13 | 119614360 | 150230984 | 30616625 |
| | | 1 | chr3 | 3 | 2.85 | 3627448 | 14951221 | 11323774 |
| | | 2 | chr6 | 1 | 1.32 | 29652898 | 33335086 | 3682189 |
| | | 1 | chr8 | 3 | 2.78 | 84674985 | 89534368 | 4859384 |
| | | 3 | chr9 | 6 | 6.24 | 37235975 | 68523573 | 31287599 |
| | | 2 | chr9 | 3 | 2.83 | 133563179 | 138394717 | 4831539 |

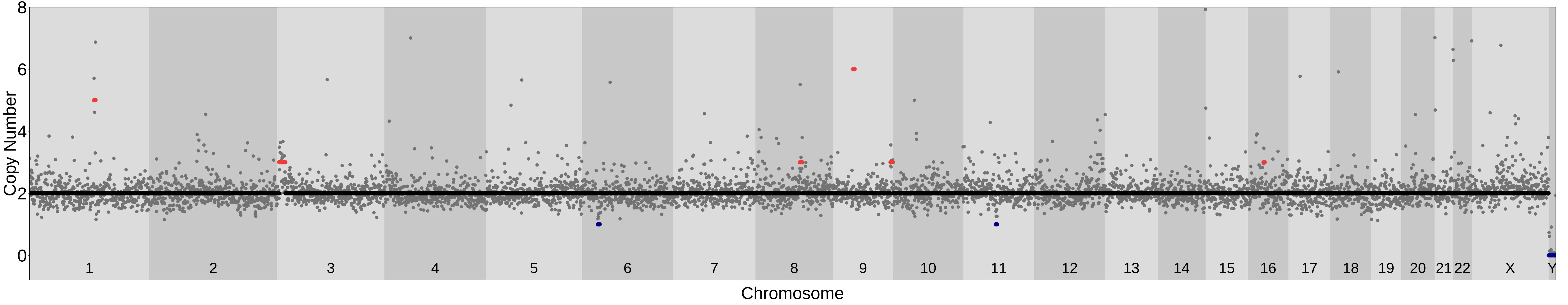

### PTA_48bp_500kb_Liftover; MAD= 0.15 Confidence_score= 0.91

| samples | ID | shared_ind_number | chr | cn | cn_median | start | end | width |
|---|---|---|---|---|---|---|---|---|
| Control | L17Exp8_1sn6_S20_R_L32Exp8_1sn6_S21_R | 2 | chr2 | 3 | 3.36 | 86594188 | 97674585 | 11080398 |
| | | 2 | chr10 | 3 | 2.79 | 38192350 | 48335174 | 10142825 |
| | | 3 | chr16 | 3 | 3.17 | 28179893 | 47026091 | 18846199 |
| | | 2 | chr17 | 3 | 2.74 | 18024196 | 27426130 | 9401935 |

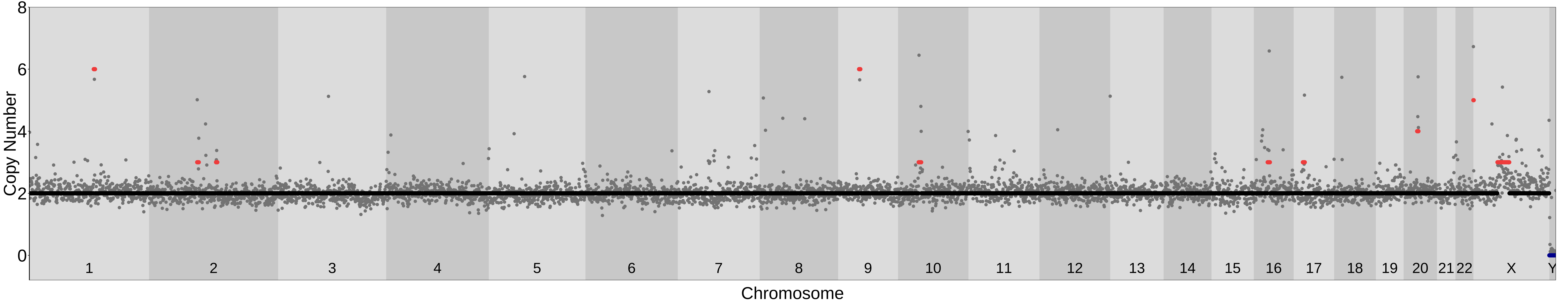

### PTA_48bp_500kb_Liftover; MAD= 0.21 Confidence_score= 0.79

| samples | ID | shared_ind_number | chr | cn | cn_median | start | end | width |
|---|---|---|---|---|---|---|---|---|
| Control | L18Exp8_1sn5_S14_R | 2 | chr2 | 3 | 3.39 | 86594188 | 97674585 | 11080398 |
| | | 1 | chr6 | 3 | 2.77 | 73685813 | 88640634 | 14954822 |
| | | 2 | chr7 | 4 | 3.70 | 56124416 | 65925737 | 9801322 |
| | | 1 | chr12 | 3 | 2.80 | 32335508 | 39787903 | 7452396 |
| | | 2 | chr17 | 3 | 2.97 | 18024196 | 27426130 | 9401935 |

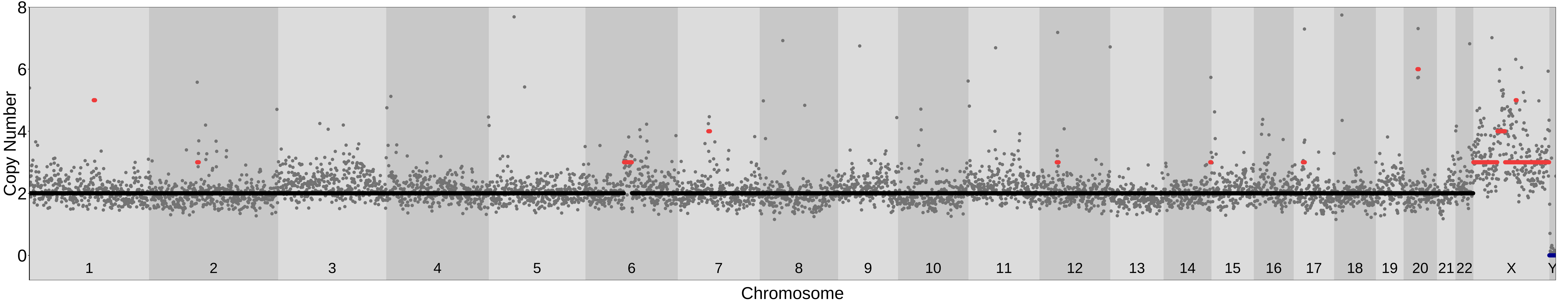

### PTA_48bp_500kb_Liftover; MAD= 0.19 Confidence_score= 0.8

| samples | ID | shared_ind_number | chr | cn | cn_median | start | end | width |
|---|---|---|---|---|---|---|---|---|
| Control | L20Exp8_2sn13_S4_R | 1 | chr4 | 1 | 1.42 | 152249878 | 157687866 | 5437989 |
| | | 2 | chr10 | 3 | 2.91 | 36903723 | 50816428 | 13912706 |
| | | 2 | chr12 | 3 | 2.74 | 128155351 | 133275309 | 5119959 |
| | | 1 | chr15 | 3 | 2.58 | 1 | 30965552 | 30965552 |

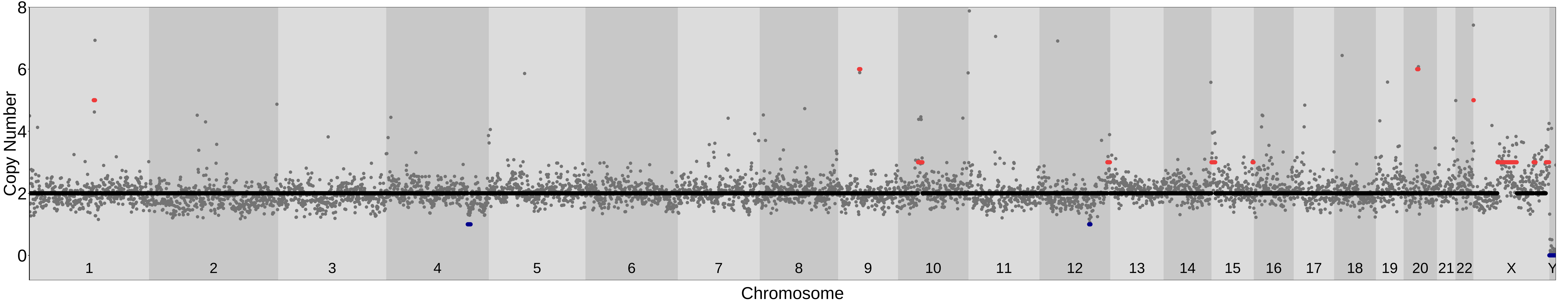

### PTA_48bp_500kb_Liftover; MAD= 0.28 Confidence_score= 0.85

| samples | ID | shared_ind_number | chr | cn | cn_median | start | end | width |
|---|---|---|---|---|---|---|---|---|
| MSA−1 | L21Exp9_1sn4_S5_R | 1 | chr1 | 3 | 2.90 | 11152572 | 22712622 | 11560051 |
| | | 1 | chr4 | 1 | 1.40 | 182023992 | 190214555 | 8190564 |
| | | 2 | chr6 | 1 | 1.42 | 25635361 | 33994564 | 8359204 |
| | | 1 | chr6 | 1 | 1.23 | 161264702 | 170805979 | 9541278 |
| | | 2 | chr7 | 3 | 3.08 | 56124416 | 77353642 | 21229227 |
| | | 2 | chr11 | 5 | 5.36 | 1 | 3757888 | 3757888 |
| | | 2 | chr11 | 3 | 2.68 | 48332652 | 65215597 | 16882946 |
| | | 1 | chr12 | 1 | 1.46 | 119509680 | 124864560 | 5354881 |
| | | 1 | chr13 | 1 | 1.25 | 110783956 | 114364328 | 3580373 |
| | | 1 | chr16 | 3 | 2.75 | 1 | 30909308 | 30909308 |
| | | 1 | chrX | 1 | 1.10 | 1 | 47580361 | 47580361 |
| | | 1 | chrX | 1 | 1.29 | 93441828 | 156040895 | 62599068 |

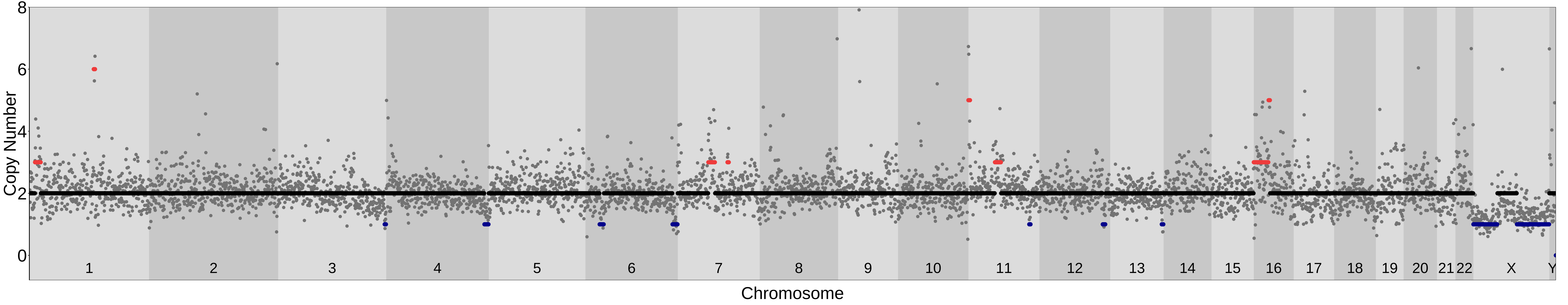

### PTA_48bp_500kb_Liftover; MAD= 0.2 Confidence_score= 0.77

| samples | ID | shared_ind_number | chr | cn | cn_median | start | end | width |
|---|---|---|---|---|---|---|---|---|
| MSA−1 | L23Exp9_1sn7_S6_R | 2 | chr2 | 3 | 3.30 | 86594188 | 97674585 | 11080398 |
| | | 1 | chr4 | 1 | 1.50 | 24691087 | 40757729 | 16066643 |
| | | 1 | chr8 | 3 | 2.66 | 32812883 | 36709997 | 3897115 |
| | | 2 | chr10 | 3 | 2.50 | 36903723 | 50816428 | 13912706 |
| | | 1 | chr10 | 3 | 2.78 | 104622572 | 110129637 | 5507066 |
| | | 2 | chr12 | 3 | 2.57 | 127606291 | 133275309 | 5669019 |
| | | 1 | chr13 | 3 | 2.62 | 53707864 | 57529239 | 3821376 |
| | | 2 | chr17 | 4 | 3.55 | 18024196 | 27426130 | 9401935 |
| | | 2 | chr21 | 3 | 2.56 | 1 | 46709983 | 46709983 |
| | | 1 | chrX | 1 | 1.19 | 1 | 48226474 | 48226474 |
| | | 1 | chrX | 1 | 1.25 | 96234700 | 156040895 | 59806196 |

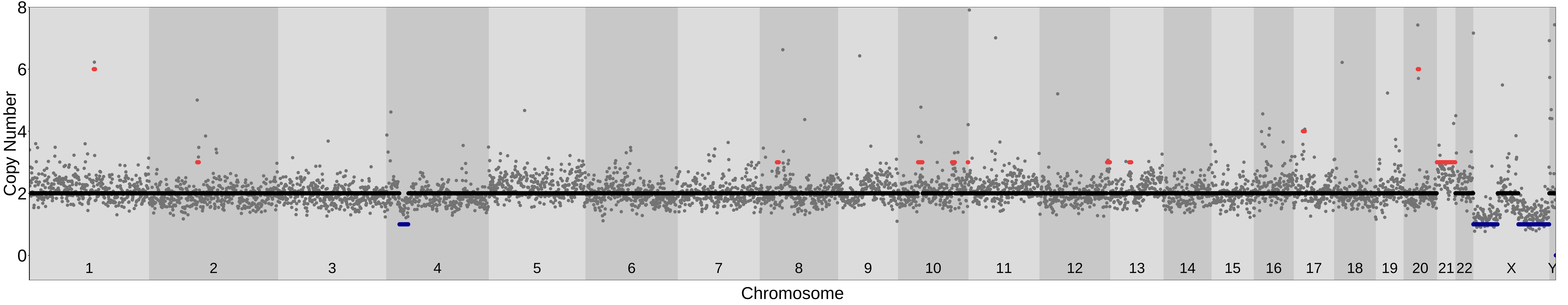

### PTA_48bp_500kb_Liftover; MAD= 0.21 Confidence_score= 0.91

| samples | ID | shared_ind_number | chr | cn | cn_median | start | end | width |
|---|---|---|---|---|---|---|---|---|
| NA12878 | L2TestPTA_2singlecell_8_S1_R | 1 | chr2 | 1 | 1.04 | 150076773 | 156700365 | 6623593 |
| | | 1 | chr3 | 1 | 1.09 | 75978290 | 83535110 | 7556821 |
| | | 1 | chr3 | 1 | 1.10 | 176053690 | 182208322 | 6154633 |
| | | 1 | chr4 | 1 | 1.16 | 17524711 | 21401853 | 3877143 |
| | | 1 | chr5 | 1 | 1.22 | 137823825 | 142975465 | 5151641 |
| | | 1 | chr7 | 1 | 1.50 | 1 | 5234863 | 5234863 |
| | | 1 | chr9 | 1 | 0.76 | 90847044 | 104007098 | 13160055 |
| | | 1 | chr11 | 1 | 0.96 | 5574009 | 14511395 | 8937387 |
| | | 1 | chr12 | 1 | 1.21 | 108772049 | 116689224 | 7917176 |
| | | 1 | chr16 | 1 | 1.30 | 55282897 | 83813102 | 28530206 |
| | | 2 | chr18 | 1 | 1.03 | 68907706 | 73239732 | 4332027 |

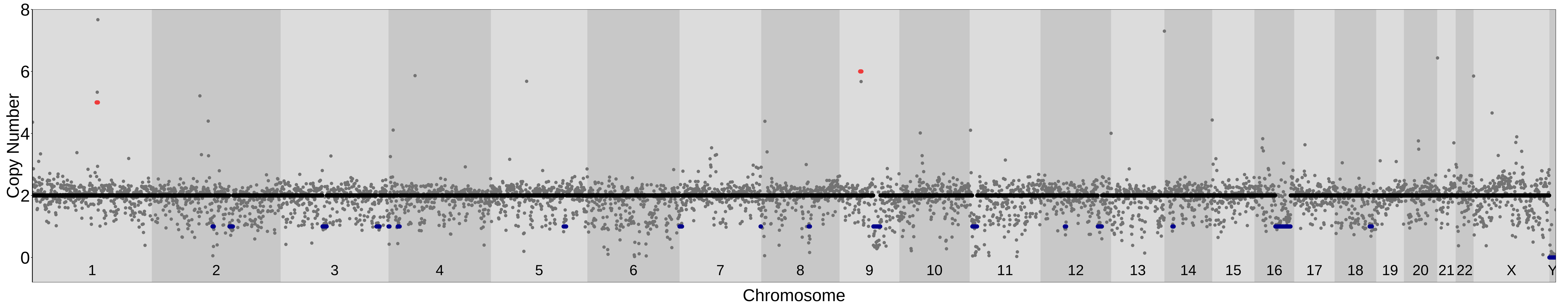

### PTA_48bp_500kb_Liftover; MAD= 0.22 Confidence_score= 0.9

| samples | ID | shared_ind_number | chr | cn | cn_median | start | end | width |
|---|---|---|---|---|---|---|---|---|
| Control | L4Exp8_1sn10_S18_R_L34Exp8_1sn10_S19_R | 2 | chr2 | 3 | 2.77 | 83220012 | 97674585 | 14454574 |
| | | 2 | chr7 | 3 | 3.05 | 56124416 | 65925737 | 9801322 |
| | | 2 | chr10 | 3 | 2.73 | 38192350 | 50816428 | 12624079 |

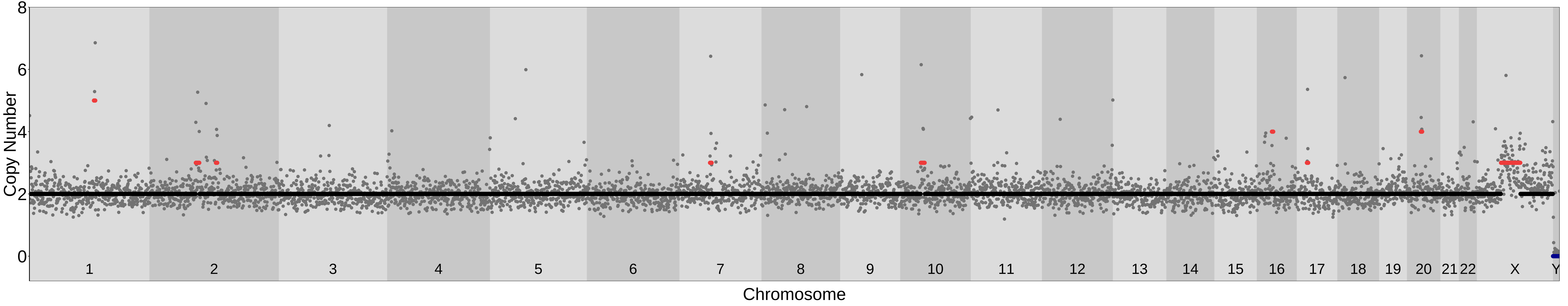
